# Morphological and functional convergence of visual projections neurons from diverse neurogenic origins in *Drosophila*

**DOI:** 10.1101/2024.04.01.587522

**Authors:** Rana Naja El-Danaf, Katarina Kapuralin, Raghuvanshi Rajesh, Félix Simon, Nizar Drou, Filipe Pinto-Teixeira, Mehmet Neset Özel, Claude Desplan

**Author notes:** These authors contributed equally (Rana Naja El-Danaf, Katarina Kapuralin).

## Abstract

The *Drosophila* visual system is a powerful model to study the development of neural circuits. Projection neurons that relay visual information from the lobula part of the optic lobe to the central brain (the lobula columnar neurons-LCNs), are thought to encode different visual features relevant to natural behavior. There are ∼20 classes of LCNs whose projections form highly specific, non-overlapping synaptic domains in the brain called optic glomeruli. Although functional investigations of several LCN circuits have been carried out, very little is known about their developmental origin and the stem cell lineages that generate the LCN subtypes. To address their origin, we used single-cell mRNA sequencing to define the transcriptome of each LCN subtype and identified driver lines that are expressed in specific LCNs throughout development. We show that LCNs originate from neural stem cells in four distinct regions in the fly brain that exhibit different modes of neurogenesis, including the ventral and dorsal tips of the outer proliferation center (tOPC), the ventral tips of the inner proliferation center (vtIPC) and the central brain (CB). This convergence of similar neurons illustrates the complexity of generating neuronal diversity in the brain and likely reflects the evolutionary origin of each LCN subtype that detects a highly specific visual feature and influence behaviors that might be specific to each species.

## Introduction

The complexity of the brain is due to the presence of a vast pool of neurons that assemble into circuits that perform different functions and control various behaviors. *Drosophila* has emerged as a prime model for understanding how neuronal diversity is generated during development. Neural stem cells (called neuroblasts, NBs) from different brain structures use distinct patterning mechanisms to produce a large diversity of neurons. This is best understood in the optic lobes that constitute about two thirds of the adult fly brain and contain ∼250 well-defined neuronal types (Doe, 2017; Holguera & Desplan, 2018; Özel et al., 2021). The fly optic lobes are made of four neuropils that mediate different aspects of visual processing: lamina, medulla, lobula and lobula plate. Most optic lobes neurons originate from two neuroepithelial regions: the outer proliferation center (OPC) produces neurons that mostly innervate the lamina and medulla, while the inner proliferation center (IPC) produces neurons that project to the lobula plate or to the medulla (Ngo et al., 2017) .

Neuronal diversity is generated by several patterning mechanisms: Temporal mechanisms, wherein NBs sequentially express a series of temporal transcription factors (tTFs), thereby producing distinct neuronal types at each division (Konstantinides et al., 2022; Li et al., 2013; Sato et al., 2013, 2019; Suzuki et al., 2013; Zhu et al., 2022), reviewed in El-Danaf et al., 2023. Spatial patterning - although NBs originating from different regions of the OPC undergo the same series of tTFs, the neurons they produce from a given temporal window may differ depending on the spatial origin of the NBs (Erclik et al., 2017). Indeed, the OPC is subdivided into at least 8 domains by the expression of the spatial transcription factors (sTFs): Visual system homeobox 1 (Vsx1), Optix, Brinker (Brk) and Retinal Homeobox (Rx), with the latter domain further subdivided by the expression of signaling molecules Decapentaplegic (Dpp) and Wingless (Wg) (Dearbon & Kunes, 2004; Gold & Brand, 2014; Islam et al., 2021; Malin et al., 2023; Perez & Steller, 1996). Two other sTFs act orthogonally and pattern the dorso-ventral axis. Disco marks the ventral domain, which is also marked by the expression of Hedgehog (Hh) during early development of the optic lobes (Hh expression is not maintained) while Spalt labels the dorsal OPC (Chang et al., 2001; Erclik et al., 2017; Valentino & Erclik, 2022). Finally, further diversity is achieved through a Notch binary cell fate decision that generates two distinct sister neurons from the single division of an intermediate progenitor (ganglion mother cell) that are produced by asymmetric divisions of NBs. The main part of the OPC (mOPC) generates most neurons of the medulla, while the tips of the OPC crescent (tOPC) are marked by Wg expression and exhibit a largely similar series of tTFs. They generate a variety of neurons that innervate several neuropils such as medulla, lobula and lobula plate (Bertet et al., 2014).

The main part of the IPC (main IPC) is divided into three domains, two domains that express Dpp and one domain expressing Brk. It generates migrating cells that become NBs upon reaching the distal part of the IPC, where they produce a short lineage and generate neurons involved in the global motion pathways that project to the lobula plate, the lamina and the medulla (Apitz & Salecker, 2015; Oliva et al., 2014; Pinto-Teixeira et al., 2018). The ventral tip of the IPC (vtIPC), marked by Wg expression, has an uncharacterized mode of neurogenesis that generates neurons whose fate has also not been determined (Apitz & Salecker, 2018).

In contrast to the development of neurons in the lamina, medulla and lobula plate, little is known about the generation of lobula projection neurons, which represent one of the major outputs of the visual system towards the central brain. Each type of lobula columnar neurons (LCNs) integrates different visual cues in the lobula to detect specific visual features and relays this information to the CB to control appropriate behavioral responses (Fischbach & Dittrich, 1989;

Otsuna & Ito, 2006; Panser et al., 2016; Scott et al., 2016; Wu et al., 2016). LCNs comprise at least 20 morphological and functional subtypes that each collect visual information from distinct layers of the lobula and projects to one of ∼20 ‘optic glomeruli’ that are highly specific, neighboring synaptic domains in the PVLP (posterior ventrolateral protocerebrum) and PLP (posterior lateral protocerebrum) regions of the CB (Fischbach & Dittrich, 1989; Otsuna & Ito, 2006; Panser et al., 2016; Wu et al., 2016). Different LCNs play a role in the detection of looming stimuli (LC4, LC6, LC16, LPLC2) (Ache et al., 2019; Klapoetke et al., 2017; Sen et al., 2017; von Reyn et al., 2017; Wu et al., 2016), figure-ground discrimination (LC9, LC12) (Aptekar et al., 2015), detection of small dark moving objects (LC11) (Keleş et al., 2020; Keleş & Frye, 2017; Tanaka & Clark, 2020), detection of small moving objects (LC18) (Klapoetke et al., 2022), or tracking of mates during courtship (LC10 group) (Ribeiro et al., 2018; Sten et al., 2021). As these neurons detect highly specific visual features, they are of critical importance for the adaptive behavior of *Drosophila*, and they have been the subject of extensive functional investigations (Morimoto et al., 2020; Städele et al., 2020). However, very little is known about their origin and development.

The identification of highly specific split-Gal4 lines that mark each of the ∼20 types of LCNs has allowed a description of how each of these LCNs collects distinct visual information in different layers of the lobula and relays it to a single glomerulus (Wu et al., 2016). However, although all LCNs appear to follow very similar rules of projection (i.e. projecting to lobula and CB), their cell bodies are in several different locations near the optic lobes, suggesting that they might originate from different progenitor pools (Fischbach & Dittrich, 1989; Otsuna & Ito, 2006; Panser et al., 2016; Wu et al., 2016) (Figure 1A).

**Figure 1:**
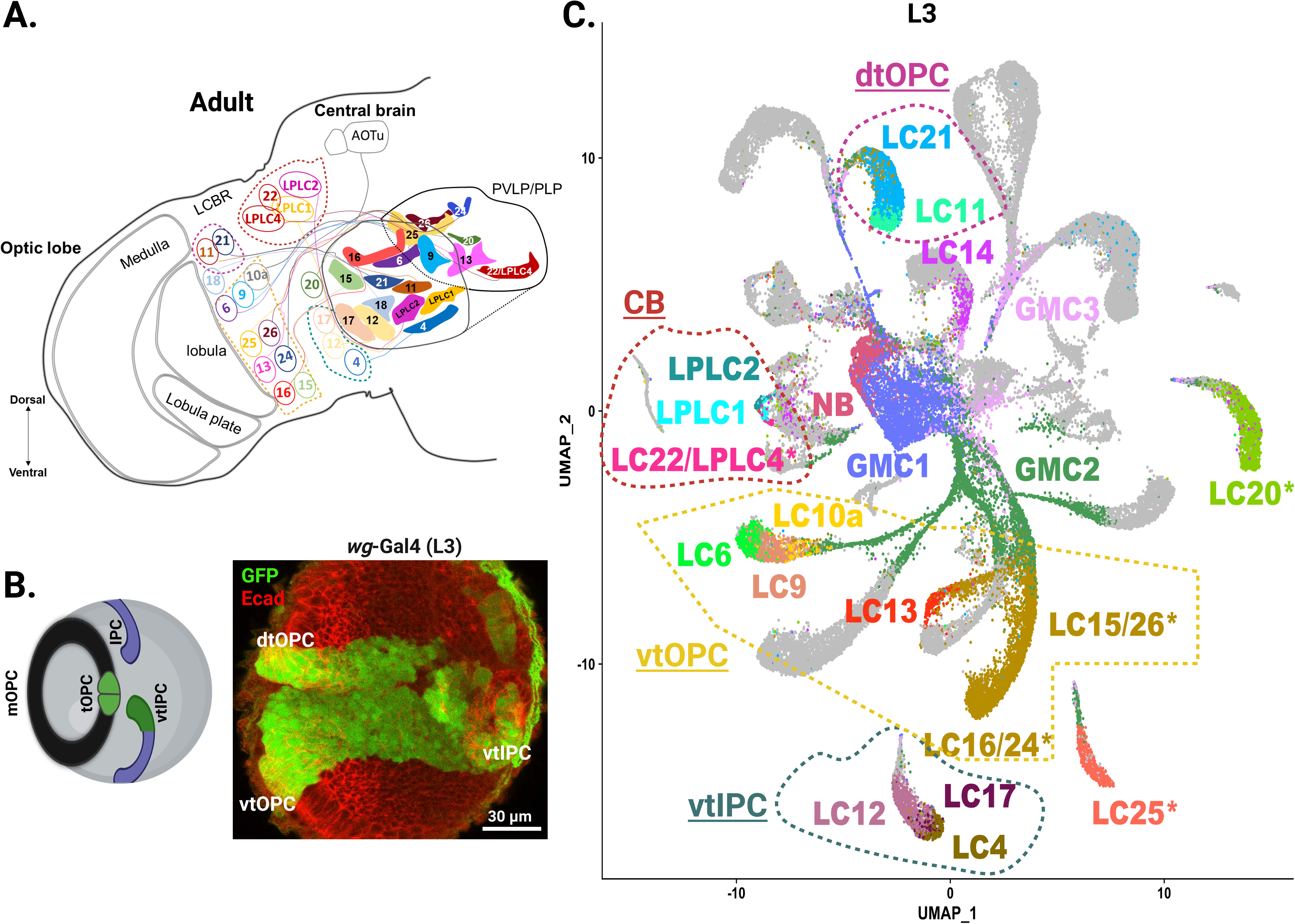
Identification of LCN subtype origin. A. Schematic of the adult brain showing the distribution of LCNs in the adult brain (based on Wu et al., 2016). Colored dashed lines delineate the origin of each LCN subtype. PVLP: posterior ventrolateral protocerebrum; PLP: posterior lateral protocerebrum; LCBR: lateral cell body rind. B. *wg-*Gal4 is expressed in the tOPC (green, tips of the outer proliferation center) and vtIPC (green, ventral tip of the inner proliferation center) in the L3 brain with Ecad (red). mOPC: main OPC; dtOPC: dorsal tip of OPC; vtOPC: ventral tip of OPC; vtIPC; ventral tip of IPC. C. Two-dimensional UMAP plot showing all LCN and progenitor clusters in L3 *wg-*Gal4 single-cell RNA sequencing datasets. Asterisk delineates heterogeneous clusters. Colored dashed lines delineate the origin of each LCN subtype.

We investigated the developmental origin of LCNs that project to the PVLP and PLP regions of the CB (Figure 1A) (Fischbach & Dittrich, 1989; Otsuna & Ito, 2006; Panser et al., 2016; Wu et al., 2016). We tested the existing split-Gal4 lines in early development, but none were expressed at late L3 stage when these neurons are produced (Wu et al., 2016). We then annotated LCNs in our developmental single-cell transcriptomic atlas (Özel et al., 2021). We found that the tOPC is the putative origin of many LCNs and thus, profiled the tOPC using single cell mRNA sequencing (scRNA-seq) at the late L3 stage to understand the mode of neurogenesis for LCNs. We used a visual system cell type classifier to identify LCN clusters in this dataset (Özel et al., 2021). We identified the clusters corresponding to all 12 LCNs that were not previously annotated in our scRNA-seq atlas (Supp Figure 1-16; Annex 1); thus, specific developmental transcriptomes are now available for all known LCNs projecting to the PVLP/PLP (Supp Figure 1-16; Annex 1). We used this information to screen Janelia Gal4 and MiMIC lines that correspond to genes specifically expressed in each of the different clusters and identified drivers that were expressed in specific LCNs throughout development (Jenett et al., 2012; Nagarkar-Jaiswal et al., 2016). This allowed us to discover that LCNs originate from various NB pools across the brain, including the dorsal and ventral tips of the OPC, the vtIPC and the CB. Interestingly, these NBs exhibit different modes of neurogenesis. LCNs that originate from the same location are related transcriptionally but often do not share related functions. This convergence of LCNs towards highly similar morphology may reflect the evolution of visual functions by co-opting and recruiting different NBs in different regions of the brain.

## Results

### LCN cluster annotation

The first step towards understanding the origin of LCNs was to annotate all LCN clusters in our scRNA-seq to define their transcriptomes throughout development (Özel et al., 2021). We previously identified several LCN clusters including: LC4, LC6, LC10 group, LC12, LC14, LC16, LC17, LPLC1 and LPLC2 (Özel et al., 2021). To annotate the remaining ones, we used the split-Gal4 lines that had been generated to label each subtype of adult LCNs (Wu et al., 2016). We combined their expression pattern with antibody staining against marker genes (such as TFs) in order to define a combination of genes specific to each LCN. During our analyses, we noticed that, in some cases, several LCNs matched to a single cluster indicating under-clustering in the original dataset (see Methods). We therefore re-clustered all cells that belonged to the clusters consistent with any of the known LCNs, and were able to identify specific clusters for each of these cell types. Detailed analysis and markers for each of the LCNs and their corresponding clusters are shown in Supplementary text (1), Supplementary figures (1-16) and Annex (1).

### scRNAseq of tOPC and vtIPC during neurogenesis

In the adult, LCN cell bodies are distributed across several distinct clusters near the optic lobes (Figure 1A) (Fischbach & Dittrich, 1989; Otsuna & Ito, 2006; Panser et al., 2016; Wu et al., 2016), suggesting that these cells are produced from different progenitors. We previously found that LC6 originates from NBs expressing the tTF Sloppy-paired (Slp) in the tOPC, the region of the OPC that expresses Wg (Fig 1B) (Bertet et al., 2014). The cell bodies of some other LCNs (LC9, LC11, LC18, LC21) (Wu et al., 2016) reside near LC6, suggesting that they might originate from the tOPC as well (Fig 1A). We therefore performed scRNA-seq of FACSed neurons labeled by a *wg-*Gal4 driver that is expressed in the tOPC and vtIPC, as well as in some neurons in the central brain (Apitz & Salecker, 2018) (Figure 1B). We dissociated brains at the late larval stage (wandering L3), during neurogenesis, and obtained 65,851 single cell transcriptomes using the Chromium system (10X Genomics). A uniform manifold approximation and projection (UMAP) plot was used to visualize distinct clusters for neuroepithelial cells, neuroblasts, GMCs, and the many neuron types of the tOPC and vtIPC (Figure 1C; Supp Figure 17; Supp Figure 18).

We adopted a supervised approach, using a neural network classifier trained on our reference atlas, to identify these cells according to the annotations we previously defined from P15 to adult stages (Özel et al., 2021). Interestingly, this classifier identified all LCNs projecting to the PVLP/PLP in this dataset except for LC18 (Figure 1C, Supp Figure 17 and Supp Table 1), which suggested that most LCNs are of tOPC or vtIPC origin, or might come from a central brain NB that expresses Wg. Many of the cell-types originating from the tOPC and vtIPC did not correspond to known LCNs and were not further studied here (Supp Figure 17 and Supp Table 1). The NBs expressing known tOPC tTFs were sequentially organized in the UMAP plot in the order of the sequential expression of the tTFs expressed: Distal-less (Dll), Eyeless (Ey), Slp and Dichaete (D) (Supp Figure 18C; Figure 5C) (Bertet et al., 2014).

### LC6, LC9 and LC10a originate from ventral tOPC neuroblasts at the Slp temporal window

We had previously shown that LC6 originates from tOPC NBs at the Slp temporal window (Bertet et al., 2014). On UMAP, LC9 cluster mapped very close to the LC6 cluster, suggesting that they share many common genes and likely a common origin (Figure 1C). Indeed, the cell bodies of LC6 and LC9 are also located near each other in adults (Figure 1A) and their axonal projections follow the same trajectory to the central brain via the anterior optic tract (AOT) and terminate at the anterior PVLP (Wu et al., 2016) (Figure 1A). Moreover, both LC6 and LC9 express *twin of eyeless (toy),* which encodes a TF marking neurons generated from Slp-expressing NBs in the tOPC (Figure 2A; Supp Figure 2) (Bertet et al., 2014). To confirm these observations, we used FLEXAMP (Flip-out LEXA AMPlification-a memory cassette tool used to identify adult neuronal subtypes by immortalization of the reporter from its expression at larval stages (Bertet et al., 2014) with the *slp*-Gal4 driver line (*R35A08*-Gal4). We recovered LC6 as expected (Figure 3A, B), but also LC9 (Figure 3B, C) that presumably could not be distinguished from LC6 in the previous study (Bertet et al., 2014) due to their extensive morphological similarities. In order to identify whether LC6 and LC9 originated from the dorsal or the ventral tOPC, we looked at *slp*-Gal4 expression in late L3, and observed two neuronal populations: the dorsal population extended processes into the medulla and central brain and were likely medulla tangential neurons (Bertet et al., 2014), while the ventral neurons extended processes into the lobula and central brain, indicative of an LCN, which are very likely LC6 and LC9 (Figure 3A). Interestingly, we also recovered neurons belonging to the LC10a-d group in these FLEXAMP experiments (Figure 3D).

**Figure 2:**
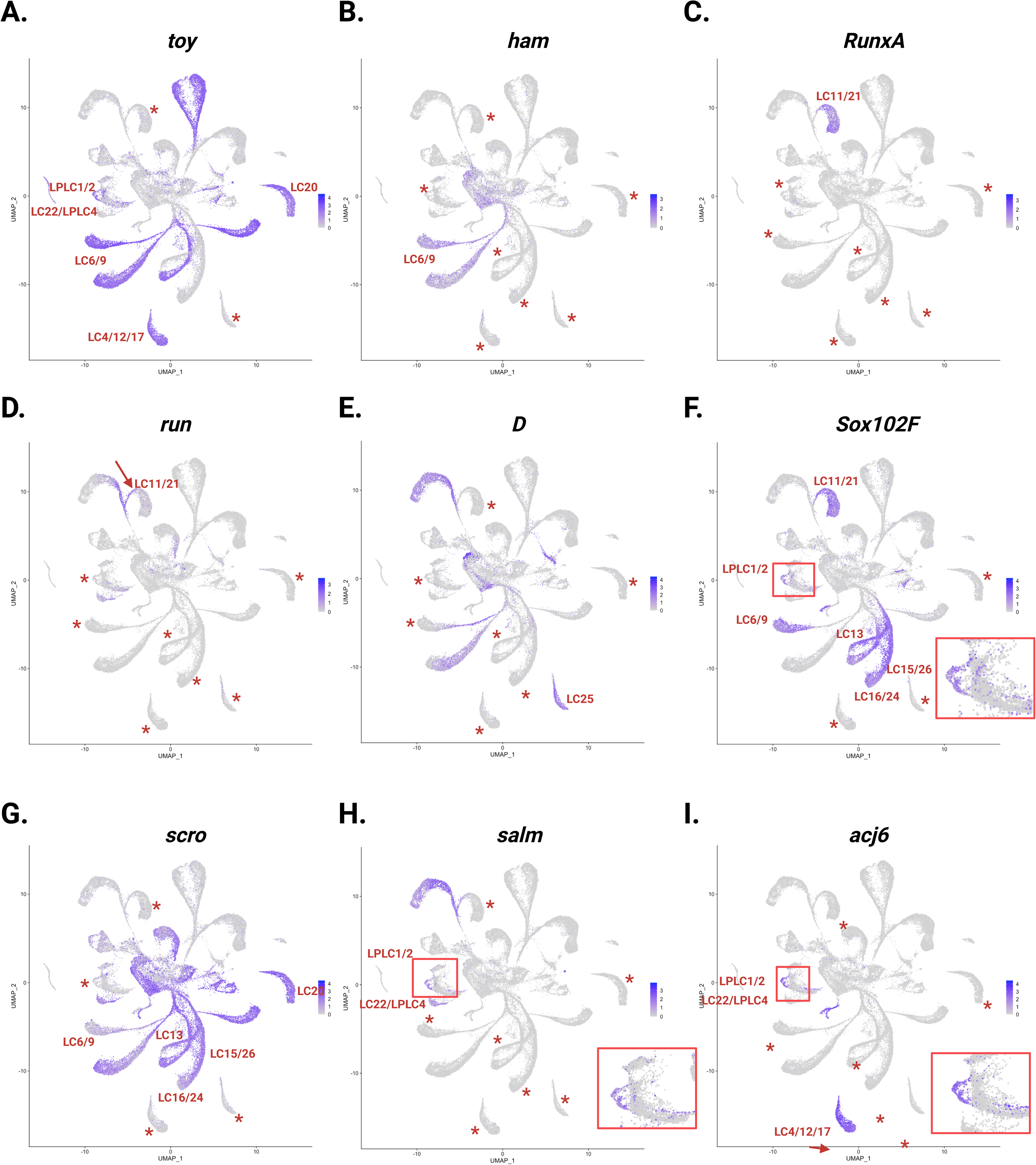
Expression of markers genes in LCN clusters of the developing Wg region. Two-dimensional UMAP plots showing the expression of different marker genes that allowed the annotation of the different LCN clusters in the Wg domain: *toy* (A), *ham* (B), *RunxA* (C), *run* (D), *D* (E), *Sox102F* (F), *scro* (G), *salm* (H), *acj6* (I). Red asterisk depicts LCN clusters where the specific gene is not expressed.

**Figure 3:**
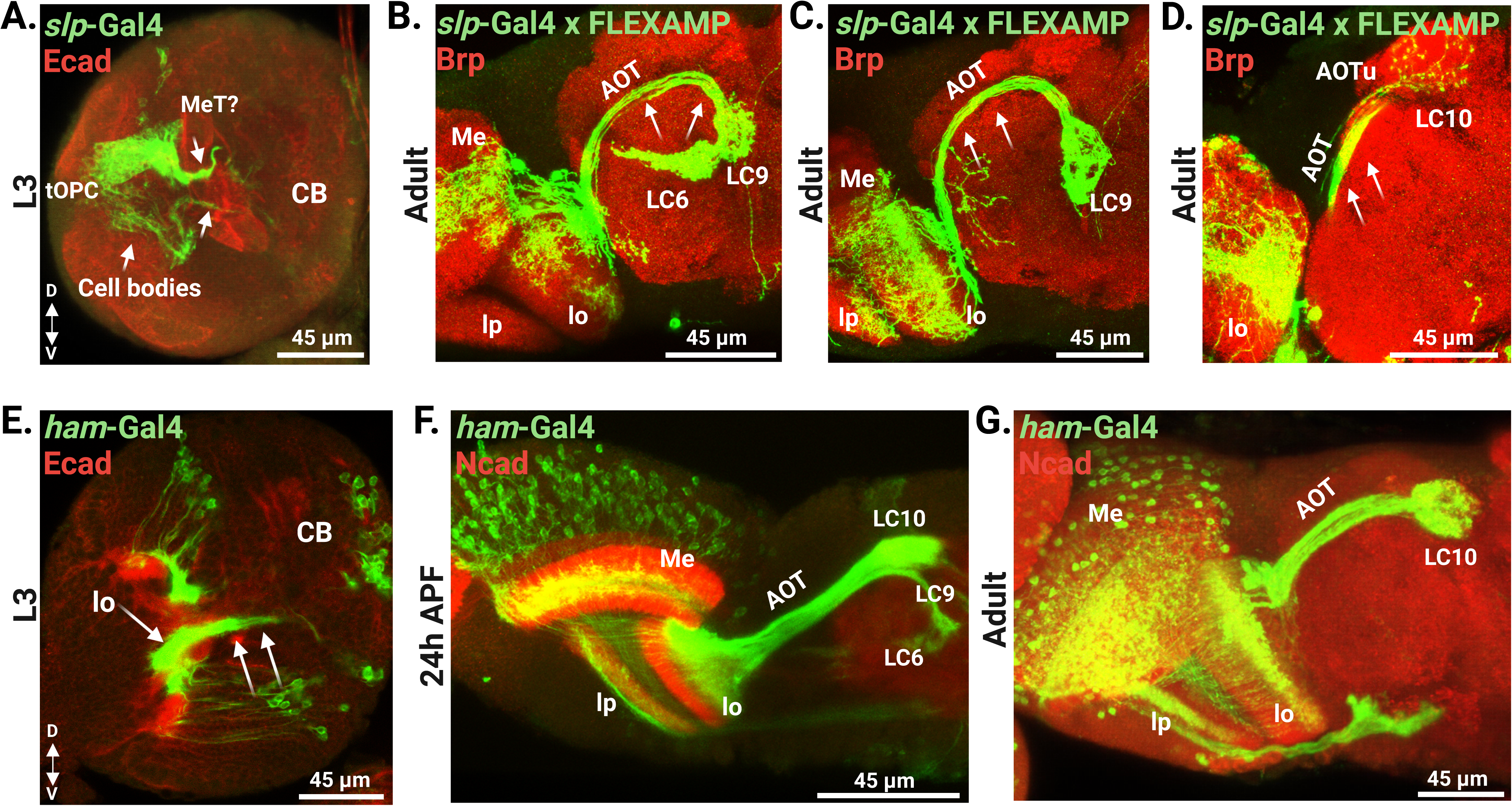
LC6, LC9 and LC10 originate from the Slp domain of the ventral tOPC. A. Expression pattern of the *slp*-Gal4 line (green) in the L3 brain with Ecad (red). Note the presence of two neuronal populations with projections extending to the central brain (CB). The dorsal population is reminiscent of medulla tangential neurons (MeT), while the ventral population resembles that of an LCN population. D: dorsal; V: ventral. B-D. FLEXAMP experiment in the *slp*-Gal4 line reveals the presence of the LC6 (B; green), LC9 (B,C; green) and LC10 (D; green) populations in the adult brain (red: Brp), demonstrated by their unique projection patterns (arrows) to the central brain via the AOT (anterior optic tract). Me: medulla; lp: lobula plate; lo: lobula; AOTu: anterior optic tubercule. E-G. Developmental expression pattern of *ham*-Gal4 in L3 (E), 24h APF (after pupal formation; F) and adult (G) brains. E. Note the presence of a ventral LCN neuronal population (green) extending towards the lobula (lo) and into the central brain (CB), similar to *slp*-Gal4 (A) (red: Ecad). F. In early pupal stages, LC6, LC9 and LC10 populations (green) can be seen extending via the anterior optic tract (AOT) into the central brain (red: Ncad). G. In the adult, only LC10s can be detected in the brain (green; red: Ncad). Me: medulla; lp: lobula plate; lo: lobula.

The gene for the transcription factor *hamlet (ham)* is expressed in the LC6 and LC9 clusters (Figure 2B). *ham*-Gal4 (*R80G09*-Gal4) recapitulated the expression of *slp*-Gal4 in L3 with both dorsal and ventral populations of the tOPC (Figure 3A, E). As the ventral population projected to the lobula and central brain (Figure 3E), they were likely LC6 and LC9. *ham*-Gal4 expression was maintained at early pupal stages in LC6 and LC9 (Figure 3F). Interestingly, it was also expressed in LC10a from early-late pupal and into adult stages (Figure 3F, G). The cluster for LC10a maps closely to LC6 and LC9 on the UMAP (Figure 1C) and the neurons share a similar trajectory towards the central brain in the adult. LC10a project from the lobula to the central brain via the same track (AOT), but terminate in the anterior optic tubercule (AOTu) as opposed to LC6 and LC9 that project in the PVLP (Wu et al., 2016) (Figure 1A). This observation suggests that LC10a also originate from the tOPC. However, the LC10 group includes multiple subtypes (LC10 a-d), which all project to the same glomerulus in the AOTu (Wu et al., 2016).

Yet, our *wg*-Gal4 scRNA-seq dataset only included LC10a that was annotated with relatively low confidence (Figure 1C; Supp Figure 17, Supp Table 1), although neural network classifications can be less accurate at very early stages. As our current analyses are focused on LCNs that project to the PVLP/PLP domains, future investigations will be needed to understand the identities of the other members of the LC10 group. Altogether, our data suggests that LC6, LC9 and LC10a emerge from close temporal windows of the ventral tOPC as they all share the expression of many genes.

### LC11 and LC21 originate from dorsal tOPC at the Dll temporal window

The LC11 and LC21 (Figure 1C) are the only clusters that express the gene encoding the transcription factor *Runt related A (RunxA*) (Figure 2C; Supp Figures 3, 7). We therefore stained the *wg-*Gal4 reporter line with RunxA antibody at L3 (Figure 4A): LC11 and LC21 cell bodies were located in the dorsal tOPC, and were produced from the earliest, Dll temporal window (Figure 4A). Indeed, LC11 and LC21 clusters also expressed *runt (run)* - a marker of neurons generated from Dll expressing NBs of the tOPC (Figure 2D) - but not *D* (Figure 2E), a marker of ventral neurons originating from the Dll temporal window (Bertet et al., 2014). Additionally, LC11 and LC21 clusters expressed *Sox102F* (Figure 2F) but not *toy* (Figure 2A) or *scarecrow* (*scro*) (Figure 2G), and anti-Sox102F labelled the LC11 and LC21 neurons in the dorsal tOPC (Figure 4B). To further confirm this annotation, we examined the expression patterns of a *RunxA* MiMIC line that marked a population of neurons in the dorsal tOPC similar to the subpopulation found with *wg*-Gal4 in L3 (Figure 4C). We confirmed that it was an LCN population since their processes extended to both the lobula and central brain (Figure 4C). This confirmed that dorsally located LC11 and LC21 are transcriptionally related, project to neighboring optic glomeruli and originate from NBs in the Dll temporal window in the dorsal tOPC (Figure 1A, C).

**Figure 4:**
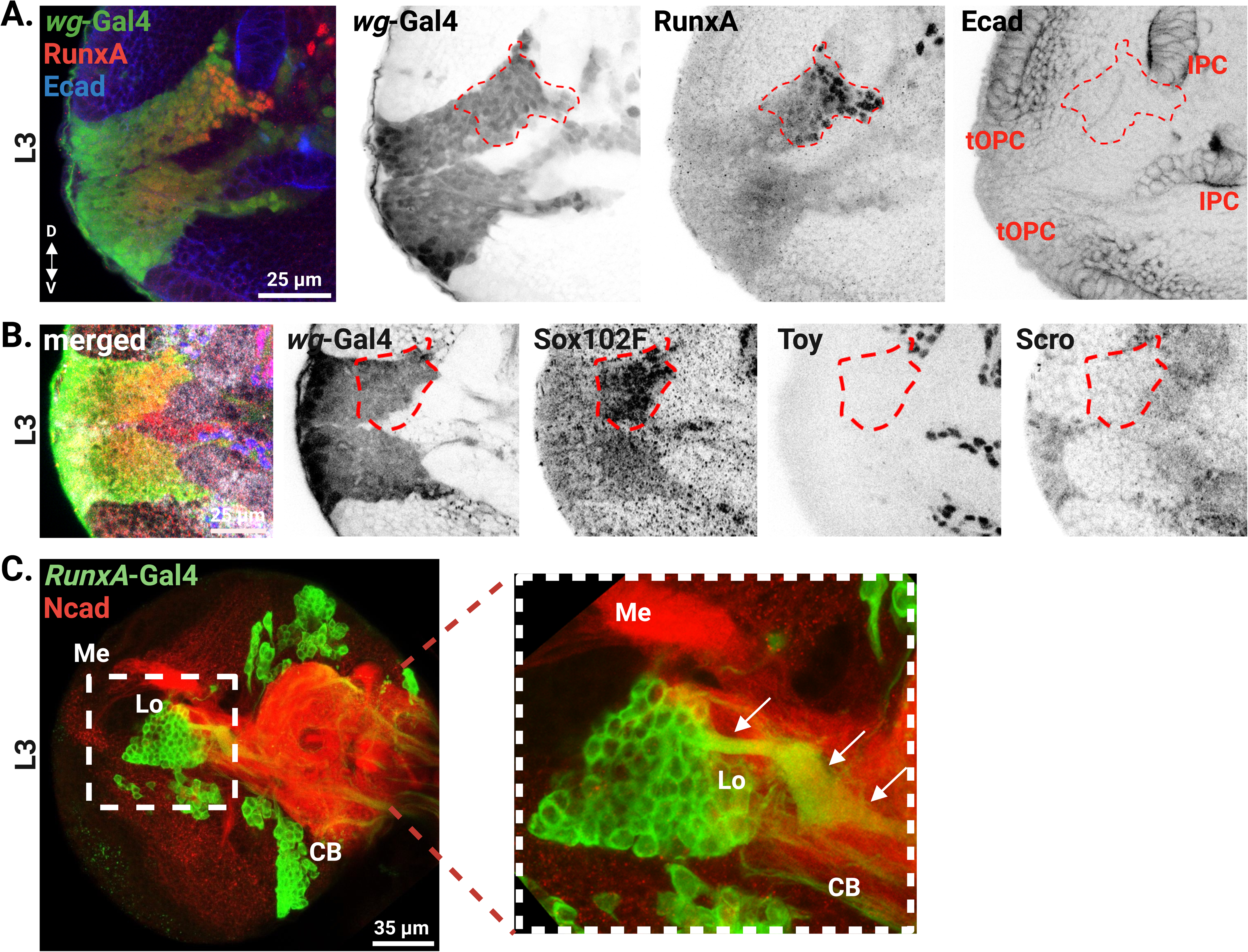
LC11 and LC21 originate from the Dll domain of the dorsal tOPC. A. Expression of RunxA (red) in the *wg-*Gal4 (green) in the L3 brain (blue: Ecad). Note that RunxA is expressed in a population of neurons located in the dorsal tOPC (dashed line). D:dorsal; V:ventral. B. LC11 and LC21 express Sox10F (red), while they don’t express Toy (blue) nor Scro (grey) in the *wg*-Gal4 line (green) in L3. Note that the Sox102F positive population (dashed line) is located in the dorsal tOPC, similar to the RunxA population (A). C. Expression pattern of the *RunxA*-Gal4 line (green) shows the presence of a LCN population (inset) with projections (arrows) to the lobula (Lo) and central brain (CB). Red: Ncad; Me: medulla.

### LC13 and LC15, LC16, LC24, LC26 originate from the ventral tOPC

The clusters corresponding to LC15/LC16/LC24/LC26 and LC13 are close to each other in the UMAP (Figure 1C). These clusters expressed *Sox102F* that is also expressed in the clusters for the other LCNs from the tOPC, ventral LC6-LC9 and dorsal LC11-LC21(Figure 2F, 4B). In the *wg-*Gal4 line, Sox102F was expressed in neurons in the dorsal and ventral tOPC (Figure 4B, 5A). *Scro* was expressed in the LC6, LC9, LC15/16/24/26 and LC13 clusters but not in LC11-LC21 (Figure 2G) and was only found in neurons in the ventral (but not dorsal) tOPC, confirming that LC11 and LC21 are produced dorsally (Figure 4B, 5B). In order to distinguish amongst neurons in the ventral groups, we used *toy* that was expressed in the LC6 and LC9 clusters but absent from LC15/16/24/26 and LC13 (Figure 2A). Co-staining for Toy and Sox102F showed that LC15/16/24/26 and LC13 neurons were located ventrally in a cluster of neurons in close proximity to LC6 and LC9 (Figure 5A).

**Figure 5:**
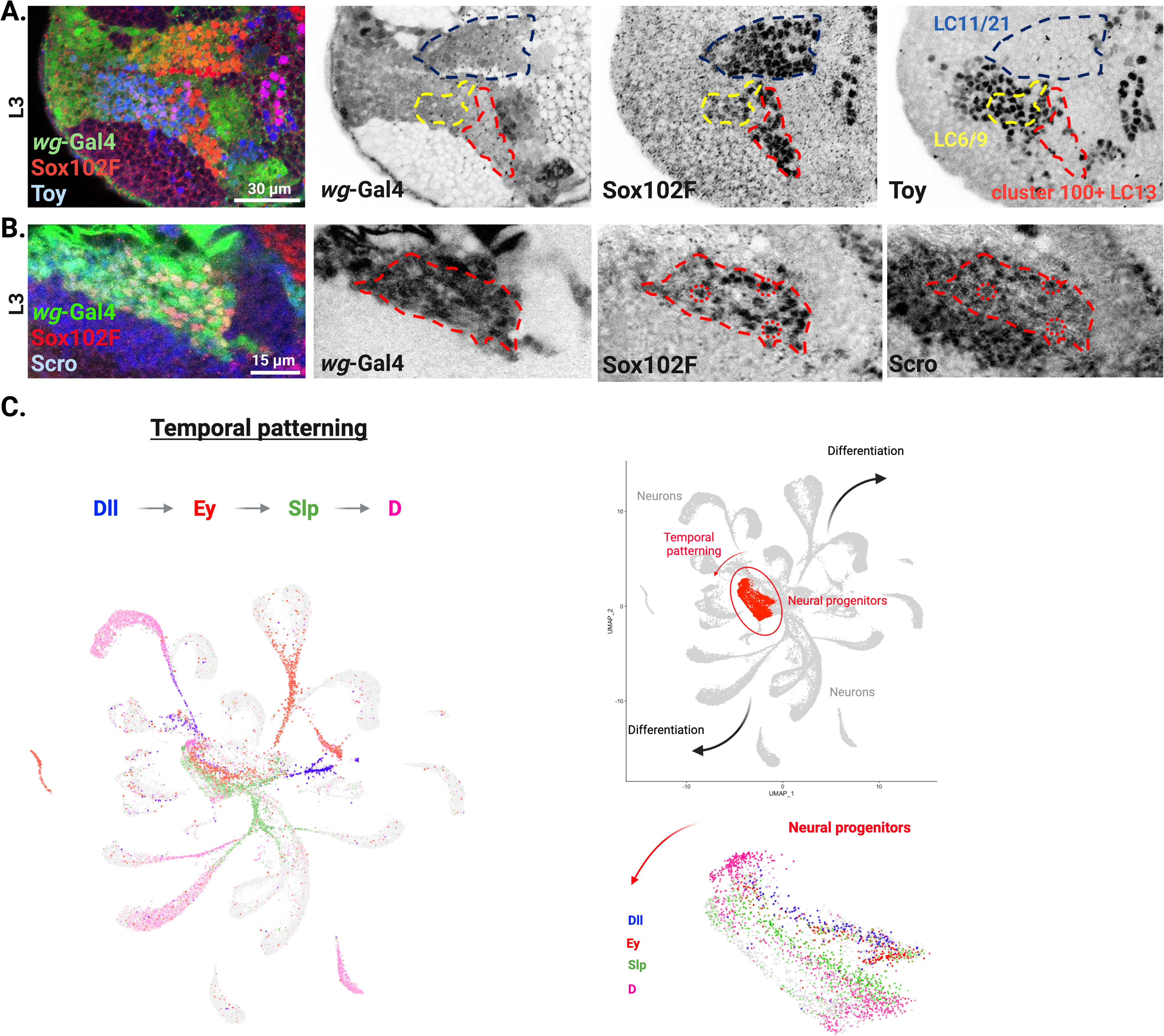
LC13, LC15, LC16, LC24 and LC26 originate from a novel domain of ventral tOPC. A. LC15, 16, 24, 26 (cluster 100) and LC13 express Sox102F (red), but not Toy (blue) in L3 of the *wg*-Gal4 line (green). These LCNs are located in the ventral tOPC (red dashed line), in close proximity to LC6/9 clusters (yellow dashed line) where they express both Toy and Sox102F. While LC11/21 clusters (blue dashed line) are located in the dorsal tOPC (see also Figure 4). B. In the ventral tOPC, cluster 100 and LC6/9 populations co-express Sox102F (red) and Scro (blue) in the *wg*-Gal4 line (green). C. Left panel: UMAP plot showing the expression of temporal transcription factors in the developing *Drosophila* tOPC. tOPC neuroblasts sequentially express Dll (blue), Ey (red), Slp (green) and D (magenta). Right panel, top: UMAP plot showing neural progenitors (neuroblasts and GMCs) in red color. Right panel, bottom: Close-up of the UMAP plot showing neural progenitors. The known tTFs (Dll Ey, Slp, D) are expressed in sets of neuroblasts organized in the plot from right to left (red arrow). The neuroblasts that do not express a known tTF are shown in gray color, suggesting that previously known temporal cascade for the tOPC may be incomplete.

Although LC15/16/24/26 and LC13 originate from the ventral tOPC, they do not express any known markers of the Dll, Ey, Slp or D temporal windows (Bertet et al., 2014) (Figure 5C; Supp Figure 18). They must therefore originate from other, yet unknown temporal windows. Using the UMAP, we examined the expression of the different tTFs in the NBs from the tOPC. In the UMAP of the L3 *wg-*Gal4 scRNA-seq, distinct subsets of NBs were arranged in a way that reflected the trajectory of their progression along the temporal series. The previously described temporal factors (Dll, Ey, Slp and D) were expressed in NBs that were organized along pseudotime while intermediate rows of neuroblasts did not express any known tTFs (Figure 5C). This indicates that the known temporal cascade for the tOPC is incomplete. The analysis of other temporal windows will be the topic of another publication.

### LC4/LC12/LC17 originate from the ventral tip of the IPC

LC4, LC12 and LC17 clusters are next to each other in the UMAP (Figure 1C) and their cell bodies are also situated close together in the lateral cell body rind (LCBR) in the adult (Figure 1A) (Wu et al., 2016) with all three subtypes co-expressing *abnormal chemosensory jump 6 (acj6) and toy* (Figure 2A, I) (Erclik et al., 2008; Özel et al., 2021). *Acj6* is not expressed in any neurons from the tOPC but it is found in the vtIPC, a domain where *wg-*Gal4 is also expressed (Figure 1B, 6A). To validate the vtIPC origin of LC4, LC12 and LC17, we screened for Gal4 lines that were expressed in the vtIPC between larval and adult stages. R65C05-Gal4 was expressed in a population of neurons in the vtIPC at late L3 stage (Figure 6B). Processes of these neurons project towards the lobula and the central brain, indicating that they are likely LCNs (Figure 6B). Furthermore, these neurons co-express Acj6 and Toy (Figure 6C). To distinguish between these three subtypes, we looked for differentially expressed genes among their clusters. *Knot (kn)* was expressed in LC4 and LC17 but not LC12 (Figure 6D) (Özel et al., 2021), while *jim lovell (lov*) was only expressed in LC4 (Figure 6E). Antibody staining at L3 and early pupal stages confirmed the presence of these three populations.

**Figure 6:**
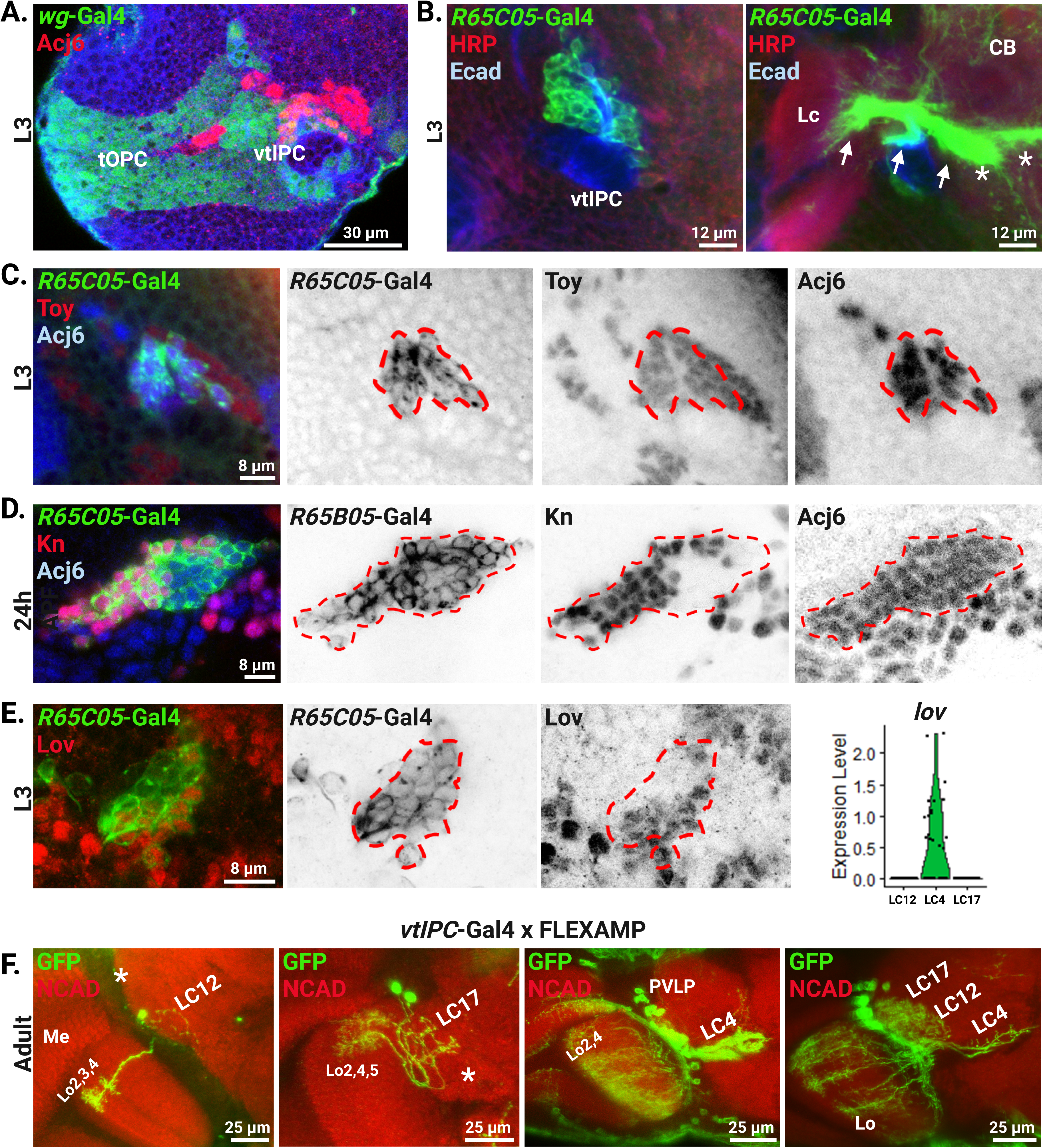
LC4, LC12 and LC17 originate from the vtIPC. A. Acj6 (red) is expressed in the vtIPC (green; *wg*-Gal4) but not in the tOPC in L3. B. Expression pattern of the *R65C05*-Gal4 (green) in L3, reveals the presence of a LCN neuronal population in the vtIPC (left panel; green). Right panel: These neurons extend projections (arrows) into the lobula complex (Lc) and central brain (CB). Note the bifurcation of the projections in the CB indicating multiple LCN subtypes (asterisk). C. LCNs in *R65C05*-Gal4 (green) co-express Toy (red) and Acj6 (blue) in L3. D. During early pupal stages (24h AFP), a population of LCNs in *R65C05*-Gal4 (green) co-express Kn (red) and Acj6 (blue), markers of LC4. E. Lov (red) a marker of LC4 is also expressed in a population of neurons in *R65C05*-Gal4 (green) confirming LC4’s vtIPC origin. F. FLEXAMP in a vtIPC-Gal4 (specific to vtIPC) reveals the presence of LC12 (green; first panel), LC17 (green; second panel) and LC4 (green; third panel). While all three LCNs are present in the fourth panel (green). These LCNs are distinguished by their unique projection pattern within the different lobula layers (Lo) and in the central brain. Asterisk represent optic glomeruli of the LC12 (first panel) and LC17 (second panel). Me: medulla; PVLP: posterior ventrolateral protocerebrum.

We performed FLEXAMP using a vtIPC-specific Gal4 line at larval stages. Since Dll is expressed in the tOPC but not in the vtIPC, we used *dll-frag3-Gal80* (driven by a *dll* enhancer) (Galindo et al., 2011) to suppress Gal4 expression in the tOPC when recombined with *wg*-Gal4 (Filipe Pinto-Teixeira; in preparation). This combination labeled LC4, LC12 and LC17 (Figure 6F) in adult brains. Altogether, these data confirm that LC4,12,17 share a common origin from the vtIPC. Unlike the OPC, neurogenesis in the vtIPC is not well understood. It is yet to be established whether LC4,12,17 originate from different temporal windows or from distinct spatial sub-domains within the vtIPC.

### LPLC1, LPLC2, LPLC4 and LC22 originate from the central brain

LPLC1, LPLC2, LPLC4 and LC22 cluster closely together in our *wg-*Gal4 scRNA-seq UMAP (Figure 1C). A common feature of these clusters is that they all co-express *acj6* and *toy* (Figure 2 A, I). Acj6 is absent from the tOPC (Figure 6A) but present in the vtIPC. However, unlike LC4, LC12 and LC17, these neurons were not recovered from the vtIPC-Gal4 FLEXAMP experiment (see above) and are thus unlikely to originate from the vtIPC.

To explore the origin of these neurons, we examined Acj6 in the larval brain where its expression was confined to a few populations, including neurons from the vtIPC (LC4,12,17) (Fig 6) and the -neurons originating from the main IPC (Pinto-Teixeira et al., 2018). The remaining large population of Acj6 expressing neurons were in the dorsal and ventral central brain, suggesting LPLC1, LPLC2, LC22 and LPLC4 originate from central brain NBs (Supp Figure 19A). Staining of the *wg-*Gal4 line at L3 for Sox102F, a gene that is expressed in LPLC1 and LPLC2 (Figure 2F inset), revealed a population of neurons in the dorsal central brain that likely included LPLC1 and LPLC2 (Figure 7A).

**Figure 7:**
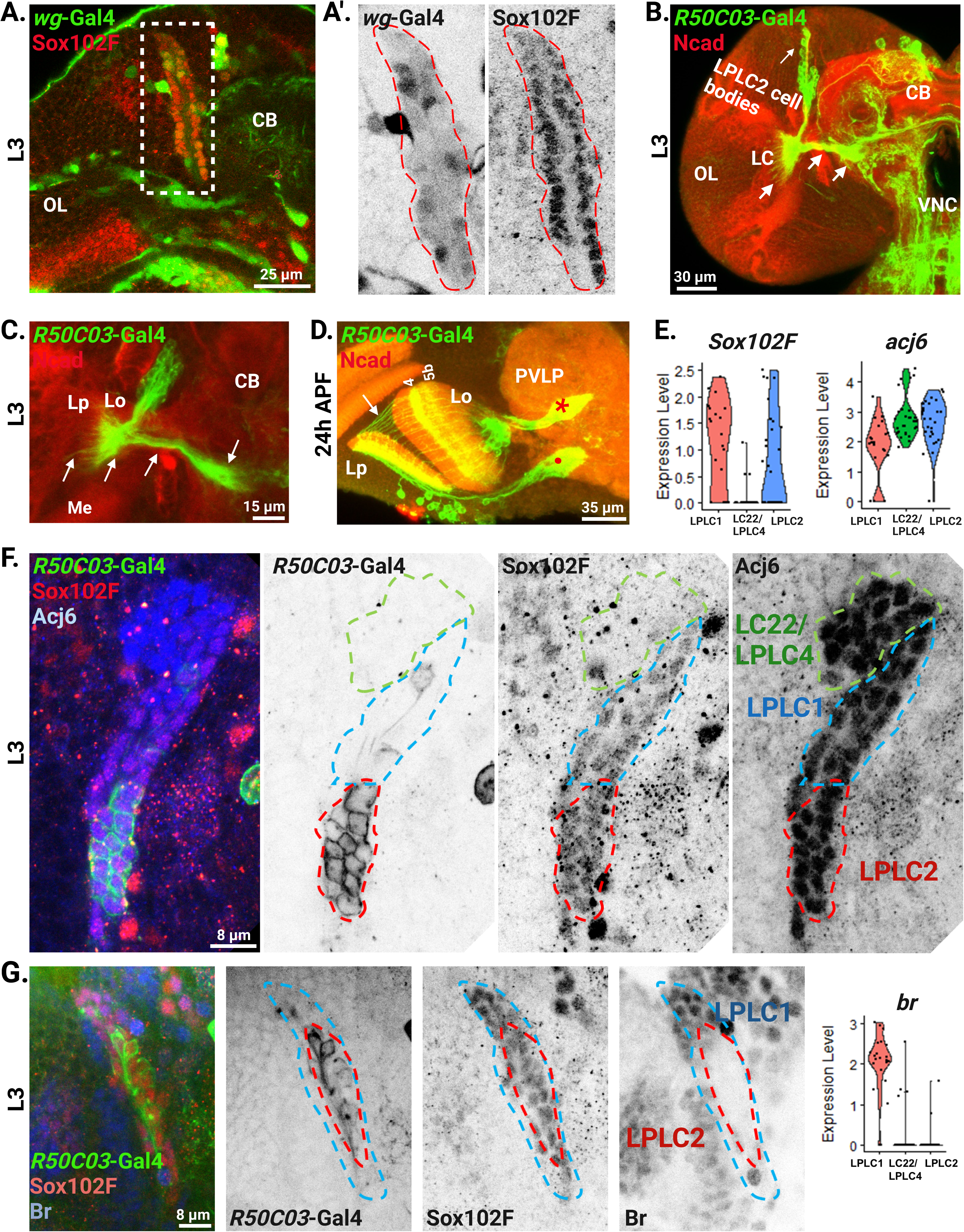
LPLC1, LPLC2, LPLC4 and LC22 originate from the dorsal CB. A. A population of central brain neurons (CB) in L3 of the *wg*-Gal4 line express Sox102F (red; Inset: A’). Ol: optic lobe. B. Expression pattern of *R50C03*-Gal4 (green) with Ncad (red) showing the presence of LPLC2 population (cell bodies in the dorsal CB), with projections (arrows) extending into the lobula complex (LC) of the optic lobe (OL), as well as the central brain (CB). VNC: ventral nerve cord. C. High magnification image of *R50C03*-Gal4 (green) with Ncad (red) in L3 showing LPLC2 neurons extending (arrows) to lobula plate (Lp), lobula (Lo) and central brain (CB). D. Expression of *R50C03*-Gal4 (green) with Ncad (red) at 24h APF, showing LPLC2 population projecting to lobula plate (Lp), and lobula (Lo) layers 4 and 5b, with a distinct optic glomerulus (asterisk) in the PVLP (posterior ventrolateral protocerebrum). Note the presence of another lobula plate neuronal population (dot). E. Violin plots showing the expression of *Sox102F* (left panel) and *acj6* (right panel) in the clusters of LPLC1, LPLC2 and LPLC4/LC22. All three neuronal populations express *acj6*. Note the absence of *Sox102F* in the LPLC4/LC22 cluster. F. Expression pattern of Sox102F (red) and Acj6 (blue) in *R50C03*-Gal4 (green) in L3. Note the presence of three populations: LPLC2 (red dashed line; GFP+, Sox102F+, Acj6+); LPLC1 (blue dashed line: GFP-; Sox102F+, Acj6+) and LPLC4/LC22 (green dashed line; GFP-, Sox102F-, Acj6+). G. Br (blue) can be used to distinguish between LPLC1 (blue dashed line) and LPLC2 (red dashed line) in *R50C03*-Gal4 (green) in L3. Both populations are Sox102F positive (red). Left panel: violin plots showing that *br* is exclusively expressed in LPLC1 but not LPLC2 nor LPLC4/LC22 clusters.

We screened for Gal4 lines with expression in the dorsal central brain similar to Acj6: *R41C07*-Gal4 and *R50C03*-Gal4 marked a central brain neuronal population whose cell bodies lied dorsally in the larval brain, expressed Acj6 and Toy, and projected to the central brain, the lobula as well as the lobula plate, indicating that they were LPLC neurons (Figure 7B, C; Supp Figure 19B). Driving the FLEXAMP reporter using *R41C07*-Gal4 line identified LPLC2, suggesting that it originates from the dorsal central brain (Supp Figure 19C). In addition, as the R50C03-Gal4 line was expressed from L3 until adult stage (Figure 7D), this allowed us to identify that population as LPLC2 based on the adult projection patterns within the lobula, lobula plate, and the distinct optic glomerulus.

To address whether LPLC1, LC22 and LPLC4, whose cell bodies were positioned close to LPLC2, were also originating from a central brain NB (Wu et al., 2016), we screened for genes differentially expressed between LPLC1, LPLC2, LPLC4 and LC22. *Broad* (*br*) was only expressed in LPLC1 while *Sox102F* was expressed in LPLC1 and LPLC2 but not LC22/LPLC4 (Figure 7E, F, G). All four neurons expressed *acj6* (Figure 7E; Supp figure 7, 8). We stained for these markers in the R50C03-Gal4 line. We identified LPLC1 that expressed Br+ /Sox102F+/ GFP- and were located near the GFP+ LPLC2 neurons (Figure 7F, G). Additionally, the Acj6+/GFP-/Sox102F-population which were in close proximity were likely LC22/LPLC4 (Figure 7F).

The arrangement of these VPNs suggest that they originate from the same NB. Their cell bodies were ordered in a way that likely reflects their birth order (Figure 7F, G) (Spindler and Hartenstein, 2010). LPLC2 cell bodies were found deepest in the brain and might be early born, followed by LPLC1 that were sandwiched in between LPLC2 and LC22/LPLC4 that were in the top layer and thus the last-born neurons in this group. They are likely to be produced by the VPNd1 NB (Yu et al., 2013). Altogether, our data indicate that LPLC2, LPLC1 and LC22/LPLC4 neurons are generated from a single NB in the central brain NBs instead of the optic lobes.

### LC18, LC25 and LC20

We were unable to identify driver lines that were expressed early in development in LC18, LC20 and LC25, and therefore could not define their origin with certitude.

We did not recover LC18 in our *slp*-FLEXAMP experiment (Figure 3A-D) or in the *wg*-Gal4 scRNA-seq dataset (Figure 1C). These neurons might be in too small numbers to be recovered in non-supervised clustering although the number of LC18 is comparable to that of LC6 and LC9 (Wu et al., 2016). However, in the adult, LC18 cell bodies were in close proximity to LC6,9,11 and 21 (Figure 1A) (Wu et al., 2016). They also expressed *toy/Sox102F/scro* that are markers shared with LC6 and LC9, but not *RunxA* (a marker of LC11 and LC21) (Figure 2A, C, F, G). We therefore suggest that they are derived from the ventral tOPC at a temporal window close to that of LC6 and LC9.

In the adult, LC25 are located in a similar position as LC13,15,16,24 and 26 that originate from the ventral tOPC (Figure 1A) (Wu et al., 2016). LC25 did not express any markers of the neurons originating from the Dll, Ey, Slp or D temporal windows (Figure 5C; Supp figure 18). Additionally, they did not express *Sox102F or acj6* that are markers of ventral tOPC, CB and vtIPC LCNs (Figure 2F, I). We conclude that LC25 likely originate from the tOPC from a temporal window that is different from all the other LCNs described here.

### Potential convergence between the transcriptomes of LCNs of different origins

The LCNs described here are generated from different origins but converge to form a common class of cell types with similar pre- and post-synaptic target regions in the brain (lobula and optic glomeruli of the PLP/PVLP). We asked whether there were any genes that could dictate this convergence to a common morphology. We analyzed markers that might be common between all LCNs (Figure 8; Supp Figure 16A). Although many transcription factors were expressed in multiple LCN types, none was expressed in all LCNs. *Sox102F*, which was restricted to the majority of LCN populations in the tOPC (Figure 2F, 4B, 5A), was also expressed in LCNs originating from the central brain such as LPLC1 and LPLC2 (Figure 2F inset, 7F, G), but not in vtIPC LCNs. *Acj6* was expressed in the central brain and vtIPC LCNs, but not tOPC LCNs (Figure 2I, 6A, C, D, 7 E, F). This lack of common TF markers suggests that shared anatomical features of LCNs have likely evolved independently in different neurogenic pools downstream of different identity TFs (Özel et al., 2022).

**Figure 8:**
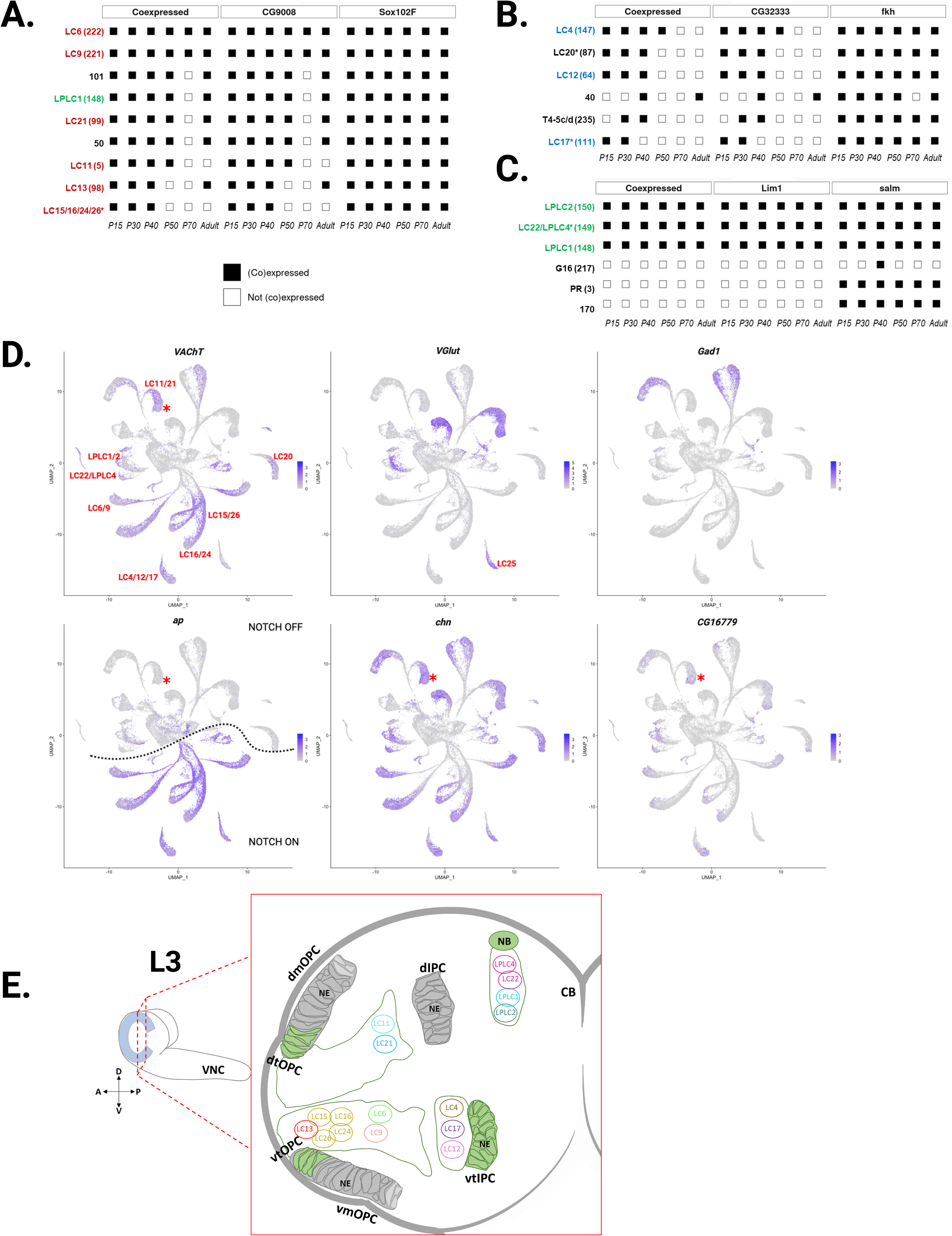
Analysis for common LCN marker genes. A-C. Mixture modeling plots for cluster marker genes organized by co-expression in P15, P30, P40, P50, P70 and Adult stages (left to right) from Özel and Simon et al. 2021. Plots were made using <u>scMarco</u> from Chen et al., 2023. Non-LCN clusters are annotated as shown in Supplementary Table 2. A. tOPC LCNs (in red) all co-express *CG9008* and *Sox102F* from P15 to P40. Co-expression is lost in some clusters after P40. B. vtIPC LCNs (in blue) all co-express *CG32333* and *fkh* from P15 to P30. Co-expression is lost in some clusters after P30. C. Central brain LCNs (in green) all co-express *Lim1* and *salm* in all developmental stages. D. Upper panel: Expression of VAChT, VGlut and Gad1, serving as markers for cholinergic, glutamatergic and GABAergic cells respectively, indicates that all LCNs (with the exception of LC 25) are cholinergic. Lower panel: Cholinergic identity is highly correlated with *ap* expression, that is Notch^ON^ (dashed line). Almost all *ap*-positive LC clusters in our dataset also express VAChT, confirming their cholinergic nature. The only exception is LC11/LC21 cluster (asterisk) that is VAChT-positive, but ap-negative. This cluster is, however, positive for *chn* and *CG16779*, two genes known to be involved in the regulation of VAChT expression. E. Schematic of the L3 brain showing the distribution of different LCN clusters based on their origin. (VNC: ventral nerve cord; dtOPC: dorsal tip of OPC; vtOPC: ventral tip of OPC; dmOPC: dorsal main OPC; vmOPC: ventral main OPC; NE: neuroepithelium; dIPC: dorsal IPC; vtIPC: ventral tip of IPC; NB: neuroblast; CB: central brain).

We then grouped LCNs based on their origin: tOPC LCNs; vtIPC LCNs and CB LCNs. Using the mixture modeling-based co-expression analysis (see Methods), we found that all tOPC LCNs expressed C*G9008 and Sox102F* until P40 (Figure 8A). vtIPC LCNs co-expressed CG32333 and *fork head (fkh)* until P30 (Figure 8B) and CB LCNs expressed *LIM homeobox 1 (lim1)* and *spalt major (salm*) throughout development (Figure 8C) and this co-expression was not seen in any other cell type. The co-expression of these marker genes specific to LCNs with common neurogenic origin might contribute to their morphological convergence.

Our investigation to understand the genetic determinants of LCN identity yielded a particularly intriguing observation: all LCNs expressed Vesicular acetylcholine transporter (VAChT), except LC25 that expressed Vesicular glutamate transporter (VGlut). LC25 is also the only LCN subtype that projects to a single layer in the lobula (Wu et al., 2016). Other neurotransmitter-related genes, such as Glutamic acid decarboxylase 1 (Gad1) were not expressed in LCN clusters (Figure 8D). Next, we checked whether *apterous (ap*) was expressed in all cholinergic LCN cells, as it is responsible for the generation of the cholinergic identity of many medulla neurons while *charlatan (chn) and CG16779* have been implicated in the regulation of VAChT in neurons where *ap* is not expressed (Konstantinides et al., 2018). Not all LCNs were *ap* positive, but all of them expressed *chn* while some expressed *CG16779*. For example, LC11 and 21 did not express *ap*, but expressed *chn* and only a subset of these cells expressed *CG16779* (Figure 8D). In the tOPC, *ap* is expressed in Notch^ON^ neurons (Bertet et al, 2014). LCNs originating from the Dll temporal window from the dorsal tOPC (LC11 and 21) did not express *ap* and were Notch^OFF^. This is consistent with previous findings that the neurons coming from Dll NBs do not undergo a terminal division (Type 0 NBs) and are thus Notch independent (Bertet et al, 2014).

All the other LCNs originating from the tOPC (LCNs originating from Slp time window and neurons coming from unknown temporal window), as well as neurons coming from vtIPC expressed *ap* and were therefore Notch^ON^. Neurons originating from the central brain and LC20, coming from yet unidentified domain, were *ap* negative, thus probably Notch^OFF^. This suggests that LCNs coming from the same spatial domains share the same neurotransmitter identity and Notch status.

## Discussion

We have identified the transcriptome throughout development of nearly all known LCNs projecting to optic glomeruli in the PVLP/PLP domains of the central brain. These neurons have at least four different origins, from both the ventral and dorsal tips of the OPC, the ventral tip of the IPC and the central brain (Figure 8E). We also showed that LCNs that share a common origin are transcriptionally similar and cluster closely together in early stages of development (Figure 1A, C; Figure 8E; Supp Figure 1). In addition, their cell bodies cluster together in the adult brain (Figure 1A). To our knowledge, this is the first description of a class of morphologically similar neurons originating from multiple distinct progenitor regions. The majority of LCNs were thought to have a central brain origin due to the location of their cell bodies in the lateral cell body rind between the optic lobes and the central brain (Fischbach & Dittrich, 1989; Yu et al., 2013), although our previous research had shown that LC6 originate from the tOPC (Bertet et al., 2014). Here, we show that, although a few LCNs do have a central brain origin, many others originate from optic lobe NBs, including ventral and dorsal tOPC and the vtIPC. This is also the first report of neurons that originate from the vtIPC. These findings add to the growing knowledge of the generation of neuronal diversity in the *Drosophila* brain and shed light on the different types of neurogenesis of this class of neurons with common anatomical and functional features.

It was previously proposed that neurogenesis in the tOPC comprises four temporal windows (Dll-Ey-Slp-D), giving rise to neurons that populate the medulla, lobula and lobula plate (Bertet et al., 2014). Our findings demonstrate that within the tOPC, some LCNs (LC11 and LC21) originate from the Dll temporal windows, others (LC6 and LC9) from the Slp window, while others (LC15/16/24/26 and LC13) originate from yet unidentified temporal domains. This is consistent with findings in the mOPC where additional tTFs were recently discovered (Konstantinides et al., 2022; Zhu et al., 2022). The identification of new tTFs in the tOPC is currently being investigated for another study. Other LCNs may originate from the vtIPC, but it is not yet known whether vtIPC NBs undergo temporal patterning, or whether they are spatially patterned. The presence of multiple LCN subtypes indicates that one or both patterning mechanisms are likely to occur in the vtIPC.

LCNs constitute a unique group of neurons in the *Drosophila* optic lobes, due to their convergence onto cell-type-specific optic glomeruli in the central brain. The optic glomeruli resemble the 52 olfactory glomeruli in the *Drosophila* antennal lobes, as each mediates a specific response by receiving information from different olfactory receptors before conveying it to the central brain. However, the development of olfactory glomeruli is much simpler: each type of olfactory sensory neurons (OSN) expressing a given olfactory receptor gene projects to one specific glomerulus in a highly stereotyped manner. OSNs undergo differentiation in the antennal disc and extend projections to the antennal lobe, where they form synapses with projection neurons in unique glomeruli. Specific OSNs expressing a particular olfactory receptor (Or) gene are precisely paired with projection neurons of distinct identities, originating at specific developmental time points (Gao et al., 2000; Gao & Chess, 1999; Jhaveri et al., 2000; Laissue et al., 1999; Vosshall et al., 2000). Thus, unlike for LCNs, the neurogenesis of olfactory neurons is homogeneous.

Does the projection pattern of an LCN within the lobula correlate with its origin? For example, do LCNs that share a common origin innervate the same lobula layers while avoiding layers where LCNs of different origin reside? Anatomical data reveal that this is not the case (Fischbach & Dittrich, 1989; Otsuna & Ito, 2006; Panser et al., 2016; Wu et al., 2016): LCNs from different origins are found across different layers that are shared with LCNs from other origins. Therefore, each LCN type has a unique innervation pattern with no apparent rules based on origin. This organization indicates that LCNs of different origins may receive input from the same presynaptic partners. Recent connectome and functional studies confirmed this. Tm5Y provides input to LC17 (vtIPC), LC6 (vtOPC) and LPLC1/2 (dCB) (Tanaka & Clark, 2022), while T3 medulla neurons connect to LC17 (vtIPC), LC11 (dtOPC) and LPLC1 (dCB) (Keleş et al., 2020; Tanaka & Clark, 2020, 2022). Similarly, T2 innervates LC4 (vtIPC), LC11 (dtOPC) and LPLC1 (dCB) (Keleş et al., 2020; Tanaka & Clark, 2020, 2022). This organization of LCNs of different origins may be essential for the development of parallel visual pathways to different stimuli to ensure the survival of *Drosophila*.

Do LCNs that cluster together and share a similar origin also share similar functional properties? LC4, LC6, LC16 and LPLC2 all can detect looming stimuli. However, they each originate from a different location in the brain (vtIPC, ventral Slp-tOPC, novel ventral tOPC temporal window and dorsal CB, respectively) (Ache et al., 2019; Klapoetke et al., 2017; Sen et al., 2017; von Reyn et al., 2017; Wu et al., 2016). Moreover, LCNs sharing similar origin can have different functional properties. For example, LC12 that detect edge generated motion onset (Städele et al., 2020), originate from the vtIPC, similar to the LC4 that detect looming speed (Ache et al., 2019; von Reyn et al., 2017). Similarly, LC6 (looming/fly take off) and LC9 (ground discrimination) both originate from the Slp tOPC (Morimoto et al., 2020; Wu et al., 2016). Finally, LC16, detect looming/backward walking and LC15 detects moving bars while sharing a ventral tOPC origin (Sen et al., 2017; Städele et al., 2020; Wu et al., 2016). Thus, the functional organization of these LCNs must result from an evolutionary adaptation of the species to specific environment and visual function whose mechanisms should be further investigated.

In the mammalian retina, retinal ganglion cells (RGCs) show many similarities to LCNs. They are also visual projection neurons comprising multiple subtypes that receive different information from other retinal neurons and transmit those to the brain for further processing (Dhande et al., 2015). Similar to LCNs, different RGCs control different behaviors that are essential for the survival of the species such as detecting looming stimuli and avoiding predators, as well as photoentrainment, pupillary light reflex, among other behaviors (Dhande et al., 2015; Farrow et al., 2013; Münch et al., 2009; Yilmaz & Meister, 2013; Zhang et al., 2012). Although they do not form optic glomeruli, RGCs terminate in different nuclei/layers in the brain in a cell type specific manner such as in the lateral genicular nucleus (LGN) and superior colliculus (Seabrook et al., 2017). For example, direction selective RGCs innervate the outer shell of the LGN, while alpha, non-direction tuned RGCs innervate the core of the LGN (Cruz-Martín et al., 2014). RGCs are early born neurons in the mammalian retina, in which neurogenesis follows a temporal patterning mechanism similar to *Drosophila* neurons (Cepko, 2014; Konstantinides et al., 2015; Nguyen-ba-charvet & Rebsam, 2020). Additionally, RGCs include more than 30 subtypes (Rheaume et al., 2018; Sanes & Masland, 2015; Shekhar et al., 2022), with evidence of different birthdates for distinct RGC subtypes (Osterhout et al., 2014), a similar ordered birthdate has been shown for mammalian retinal bipolar cells (West et al., 2022). However, the mechanisms of RGC subtype neurogenesis are still not well understood: it is not yet known whether different RGC subtypes are born from similar progenitor pools, or whether convergence in RGCs neurogenesis occurs similar to LCNs. This study may shed light on this question. RGCs and LCNs both evolve rapidly as species need to generate new neurons to adjust to novel environment (Hahn et al., 2023).

## Supporting information

Annex 1

## Acknowledgements

We would like to thank all members of the Desplan laboratory in Abu Dhabi and in New York for helpful discussions. We especially thank Mehar Sultana for her valuable technical support with FACS and 10X Genomics experiments and Marc Arnoux for his assistance with sequencing. Part of this work was carried out using Core Technology Platform resources at NYUAD with the help of Rachid Rezgui. We thank Isabel Holguera, Yen-Chung Chen and Yu-Chieh David Chen for critical reading of the manuscript. Figures in this article were created with BioRender.com.

## Methods

### Genetics and Immunohistochemistry

Male and female flies were dissected at different developmental stages: late larval stage (L3), 24 hours (h) after puparium formation (APF), 48h APF, 72h APF, 96 h APF and 3-5 days after eclosion (adult stage). Flies were maintained at 18°C-25°C, while crosses were maintained at 25°C. For FLEXAMP experiments, crosses were maintained at 18°C, moved to 29°C for heat shock (12h-24h), and then maintained at 18°C until eclosion. All genotypes used in this study are detailed in Supplementary table (Supp table 2).

Brains were dissected in cold 1X phosphate buffer saline (PBS) and fixed in 4% paraformaldehyde in PBS (PFA) at room temperature for 25-30 minutes. Brains were then washed in PBS for 30 minutes, followed by a 30 minutes wash in 0.5% PBST (PBS + 0.5% Triton X100). Brains were then incubated for 30 minutes in blocking solution (PBST + 0.5% goat serum), which was used to dilute primary and secondary antibodies. Brains were incubated for 2 nights in primary antibody at 4°C, followed by 3x washes in PBST (30 minutes each) at room temperature, and incubated for 2 nights in secondary antibody at 4°C. Brains were washed 3x in PBST (30 minutes), followed by 1x wash in PBS (30 minutes) at room temperature. Brains were mounted with Vectashield and imaged using a Leica SP8 confocal microscope or a Leica Stellaris-8 microscope, using a 40x oil lens (NA=1.4) and a 40x glycerol lens (NA=1.25), with a 1024x1024 pixel resolution. Images were analyzed and processed using ImageJ.

The following primary antibodies and concentrations were used: mouse anti-Acj6 (1:20); rat-anti Bab2 (1:1000); mouse-anti Br (1:500); mouse-anti Brp (1:30); rabbit-anti Caps 1:200; mouse-anti Con (1:100); mouse-anti Ct (1:20); guinea pig-anti Dicheate (1:50); guinea pig-anti Disco (1:100); Rat-anti E-cad (1:20); guinea pig-anti Fred (1:100); rabbit-anti GFP (1:250); sheep-anti GFP (1:250); guinea pig-anti Kn (1:100); rabbit-anti Kn (1:50); guinea pig-anti Lov (1:500); rabbit-anti Mirr (1:500); rat-anti Ncad (1:20); rat-anti pdm3 (1:1000); mouse-anti Pros (1:200); mouse-anti RFP (1:1000); rabbit-anti RFP (1:1000); rabbit-anti RunxA (1:500); guinea pig-anti Rx (1:500); guinea pig-anti Scro (1:100); rabbit-anti Sox102F (1:400); rabbit-anti SoxN (1:100); mouse-anti Ten-m (1:500); rabbit-anti Toy (1:50).

The following secondary antibodies were used at a dilution of 1:200 for all: donkey-anti guinea pig (555, 647); donkey-anti mouse (555, 647); donkey-anti rabbit (488, 555); donkey-anti rat (405, 555, 647, cy5); donkey-anti sheep (488).

Origin of all antibodies and fly lines used are detailed in Supplementary table 2.

### Single cell sample preparation

Male flies of the *wg-*Gal4 line were crossed to virgin females from *UAS-Red Stinger* (BL#8547) to label nuclei of specific *wg-*Gal4-positive cell types. Brains were dissected from *wg-*Gal4 wandering third instar larvae in ice-cold Schneider’s medium (Gibco), complemented with 10% Fetal Bovine Serum, 1% Penicillin-Streptomycin, 200mM L-Glutamin, 4mg/ml L-Glutathione and 20ug/ml insulin (all from Sigma-Aldrich). Dissected brains were dissociated into a single cell suspension by incubating in 2mg/ml collagenase and 2mg/ml papain (Sigma-Aldrich) in complemented Schneider’s medium for 30 min at 25°C. The enzymes were then removed and replaced with complemented Schneider’s medium. The brains were further mechanically dissociated by gently pipetting up and down the content of the tube 300 times until most large chunks of tissue are dissociated. The sample was then filtered using 20 μm cell strainers (pluriSelect) and kept cold by putting tube on ice. The cells expressing the transgene were sorted on the basis of their fluorescent signal using Fluorescence-Activated Cell Sorting (FACS Aria III). The cells were sorted in a tube containing PBS + 0.1%BSA. The concentration of cells was calculated based on FACS parameters.

### Library preparation, sequencing, and processing

Towards the end of sorting, we placed all appropriate reagents (as indicated in the manufacturer’s protocol, Chromium Single Cell 3’ Reagent Kits v3 chemistry) to equilibrate at room temperature. Droplet-based purification, amplification and barcoding of single-cell transcriptomes were performed using Chromium Single Cell 3’ Reagent Kit v3 (10x Genomics) as described in the manufacturer’s manual, with a target recovery of 8,000 cells per experiment. We prepared 15 libraries which were subjected to paired-end sequencing (26 × 8 × 98) using the Illumina NovaSeq 6000 system (New York University Abu Dhabi) to an average 50,000 reads per cell sequenced. We mapped the sequenced libraries to the *Drosophila melanogaster* genome assembly BDGP6.88 using CellRanger 3.0.1. For the downstream analysis, we used Seurat v4. We kept all cell barcodes that have unique feature counts between 200 and 2,500, with more than 1,000 UMIs and less than 5% of UMIs corresponding to mitochondrial genes. After processing, the dataset comprised 65,851 cells passing quality filters, with a median of 1,812 genes per cell.

To cluster our integrated single-cell transcriptomes to identify groups of cells with similar gene expression patterns, we used the functions FindNeighbours and FindClusters with default parameters except for the number of principal components (PCs), which are used to reduce the dataset dimensionality and calculate the distance between all pairs of cells, and the resolution, which is used to compensate for the tendency of modularity optimization algorithms to merge small clusters. The dataset was clustered with a dimensionality of 30 and a resolution of 2,5.

#### Neural network classification

The final Seurat object consisting of all 15 libraries was classified using our existing P15 stage neural network model with the original cluster identities (Özel et al., 2021) and the docker image: <u>yenchungchen/optic-lobe-nn</u>. We used 496 marker genes to run the classifier and obtained a median prediction confidence of 0.58 (Supp Figure 17) and an average median confidence of 0.51 per cluster.

#### Mixture modelling analysis

Transcript levels are not directly indicative of protein expression (Davis et al., 2020). To overcome this, we used a mixture modelling approach to assign a probability of expression for each gene and in each cluster of our scRNA-seq atlas. Briefly, mixture modeling approach assumes that each gene exists in two states, ON or OFF, and by comparing transcript levels across multiple cell types: those with higher transcripts levels will be considered more likely to be ON, while those with lower (but not necessarily null) transcript levels will be considered OFF (Davis et al., 2020). To create the mixture modelling plots, we used the values we previously calculated in (Chen et al., 2023; Özel et al., 2021) with a mixture modeling threshold of 0.5 for the plots in Supp Figure 17.

### Re-analysis of the existing scRNA-seq data

Cluster analysis protocols were done as previously described in (Özel et al., 2021). As we sought to assign each LCN type to specific clusters in our previously published scRNA-seq atlas (Özel et al., 2021), we found that the transcriptomic profiles of some clusters were consistent with the marker gene expression pattern of multiple LCN types (see Supp Data, cluster annotations). For instance, LC15, 16, 24, 26 were all consistent with cluster 100 (Supp Figure 15) while LC22 and LPLC4 were both mapped to cluster 149 (Supp Figure 8, 9).

The relatively low abundance and similarity of LCNs could make them particularly susceptible to such under-clustering in the scRNA-seq atlas. We therefore isolated the cells belonging to the 19 clusters consistent with any of the known LCNs (including those we had previously annotated) in the adult stage and reanalyzed them using Seurat v4.1. The data was re-normalized using SCTransform (variable.features.n = 2000) and batch corrected using the standard Seurat integration workflow with default parameters. Clustering was then performed on the integrated data using the first 50 principal components (resolution = 4), which revealed that clusters 27 (LC14), 57 (LC25), 79 (LC10b), 87 (LC20), 100 (LC15/16/24/26) and 149 (LC22 and LPLC4) contained multiple cell types (Supp Figure 1). Each LCN type could then be unambiguously assigned to a single subcluster using marker gene expression patterns. As LC14 is the only cell-type that expresses *ato* in adult optic lobes, and we could observe *ato* expression in both subgroups of cluster 27, we annotated these as LC14a and LC14b. Note that one of the two subclusters of clusters 79 and 87, one of the five subclusters of the cluster 100, as well as three of the four subclusters of cluster 57 remained unannotated and might represent yet undescribed LCN or LCN-like neurons.

We then quantified the relative transcriptomic similarity of adult LCN clusters with BuildClusterTree() function based on the same 50 PCs as above. In order to perform the same analysis during the early development of LCNs, we similarly isolated the LCN clusters in the P15 scRNA-seq data (Supp Figure 15A). However, the fact that neurons have not yet fully differentiated at this stage (shortly after neurogenesis) as well as the relatively lower number of cells sequenced compared to the adult dataset have prevented clustering P15 LCNs to the same resolution as in adult. Thus, we performed the cluster tree analysis on the original clusters, using 50 PCs calculated specifically on the LCN clusters at P15 (Supp Figure 16B).

## Supplementary data

### Cluster annotations

We adhere to the original cluster numbering/nomenclature used by (Özel et al., 2021) for the purpose of these annotations (Supp Figure 1-16; Annex 1).

#### LC9/Cluster 221

LC9 cell bodies are situated in the dorsal lateral cell body rind (LCBR), extend processes along the anterior optic tubercle (AOT), and terminate in the anterior PVLP (dorso-medial) in close proximity to the LC6 optic glomerulus (Wu et al., 2016). They were found to play a role in figure ground discrimination, and their activation caused a backward walking behavior in flies (Wu et al., 2016). By screening various markers using antibody staining, as well as several gene trap lines, we identified the following positive markers for LC9: Ac76E MiMIC line n= 5 optic lobes (Ol)), Toy (n= 7 Ol), Sox102F (n= 4 Ol), Con (n= 3 Ol; Timaeus et al., 2020), Caps (Timaeus et al., 2020), and Ten-m (n= 3 Ol). Additionally, LC9s were negative for: Acj6 (n= 7 Ol), Kn (n= 4 Ol; MiMIC), and Sns (n= 3 Ol; MiMIC). The best match for LC9 is cluster 221 (Supp Figure 2).

#### LC11/Cluster 5

LC11 cell bodies are located in the dorsal LCBR adjacent to the surface of lobula, and have an extended optic glomerulus sandwiched between the dorsal and ventral glomeruli, next to LC21 cell bodies (Wu et al., 2016). LC11’s function has been studied recently and were shown to be involved in identifying the movement of small dark objects (Wu et al., 2016). We identified cluster 5 as that belonging to LC11 using the following markers (Supp Figure 3): AChRα1, nAChRα6, nAChRα7 (all 3 (Keleş et al., 2020)); Rdl; Beat-Ic (n= 3 Ol; MiMIC); Con (n= 3 Ol; antibody staining); RunxA (n= 5 Ol; antibody staining); Sox102F (n= 3 Ol; antibody staining). While LC11 were negative for the following: nAChRα5 (Keleş et al., 2020); Acj6 (n= 7 Ol), Toy (n= 7 Ol), Ten-m (n= 3 Ol; antibody staining); Kn (n= 4 Ol) (MiMIC). It is important to note that RunxA expression is restricted to a few clusters alone, with the highest expression profile seen for cluster 5.

#### LC13/Cluster 98

LC13 cell bodies are located in the ventral LCBR with their optic glomerulus position in the posterior PVLP (Wu et al., 2016). Currently, no specific function has been reported for LC13. Based on antibody staining, LC13 were found to express: Sox102F (n=5 Ol), Pros (n= 4 Ol), Lov (n=4 Ol). Whereas they are negative for: Bab2 (n= 5 Ol), Rx (n= 3 Ol), br (n= 3 Ol), Pdm3 (n= 5 Ol), Acj6 (n=5 Ol), Scro (n= 3 Ol), RunxA (n= 3 Ol), D (n= 3 Ol), Toy (n= 3 Ol). Analyses identified cluster 98 as the candidate for LC13 (Supp Figure 4).

#### LC18/Cluster 139

LC18 cell bodies are located on the dorsal-posterior shell of the lobula, with their optic glomerulus neighboring LC12 (ventral-anterior PVLP) (Wu et al., 2016). Based on antibody staining, the following markers were identified for LC18, consistent with cluster 139 (Supp Figure 5): Toy (n=4 Ol), Sox102F (n= 3 Ol), Ct (n=5 Ol), Scro (n= 4 Ol), Lov (n= 7 Ol). Conversely, no expression was detected in LC18s for: Acj6 (n=3 Ol), SoxN (n= 5 Ol), Pdm3 (n=5 Ol), Br (n= 4 Ol), Rx (n=4 Ol), RunxA (n= 3 Ol).

#### LC20/Cluster 87

LC20 cells bodies can be found in the dorsal anterior LCBR, with processes terminating in the posterior PVLP (Wu et al., 2016). LC20s were found to express several transcription factor markers including: Lov (n= 7 Ol), Ct (n= 4 Ol), Pros (n= 3 Ol), Toy (n= 4 Ol), Scro (n= 5 Ol), and were negative for: Acj6 (n= 5 Ol), Sox102F (n= 5 Ol), Br (n= 6 Ol), Bab2 (n= 5 Ol), Rx (n= 5 Ol), Pdm3 (n= 6 Ol), SoxN (n= 3 Ol), D (n= 5 Ol), RunxA (n= 4 Ol), Kn (n= 4 Ol). This indicated that these cells belong to cluster 87 which is a heterogenous cluster (See Methods; Supp Figure 6).

#### LC21/Cluster 99

LC21 cell bodies are positioned on the surface of the lobula (posterior-medial), with their optic glomerulus adjacent to LC11 (Wu et al., 2016). Interestingly, LC21 expressed RunxA (n= 5 Ol) similar to LC11. Additionally, Sox102F (n= 3 Ol) and Pros (n= 3 Ol) were positive markers for LC21. Moreover, they were negative for: Br (n= 3 Ol), Pdm3 (n= 3 Ol), Acj6 (n= 3 Ol), Bab2 (n= 3 Ol), Scro (n= 3 Ol), Toy (n= 3 Ol), Rx (n= 4 Ol). This combination of positive and negative markers is unique to cluster 99 (Supp Figure 7).

#### LC22 and LPLC4/ Cluster 149

LC22 cell bodies are situated in the dorsal LCBR, while their axons terminate in the posterior PVLP. They were found to elicit backward walking with optogenetic activation (Wu et al., 2016). Screening for several transcription factors showed that LC22 fall in cluster 149 as they express Acj6 (n= 5 Ol) and toy (n= 5 Ol), while they are negative for: Sox102F (n= 5 Ol), SoxN (n= 4 Ol), Pdm3 (n= 4 Ol), Br (n= 3 Ol), Rx (n= 3 Ol), RunxA (n= 5 Ol), Mirr (n= 5 Ol), Lov (n= 3 Ol), Scro (n= 4 Ol). Our analysis revealed that this cluster is heterogeneous containing multiple subclusters including the one for LC22 (See Methods; Supp Figure 8).

Additionally, LPLC4 constituted another subcluster of 149(a). They were positive for: Acj6 (n= 5 Ol) and Salm (n= 4 Ol). LPLC4 were negative for: Sox102F (n= 5 Ol), Lov (n= 4 Ol), Br (n= 4 Ol), Rx (n= 6 Ol), D (n= 6 Ol), ct (n= 3 Ol), Bab2 (n= 4 Ol), Olid2 (n= 4 Ol), cro (n= 3 Ol), RunxA (n= 3 Ol), Kn (n= 3 Ol), Disco (n= 4 Ol). Although LPLC4 extend to innervate the lobula plate, it’s important to note that they share many similarities with LC22 (149b) (Supp Figure 1A) such as cell body location and optic glomerulus projection in the PVLP (See Methods; Supp Figure 9).

#### LC25/Cluster 57

LC25 cell bodies reside in the ventral LCBR and their optic glomerulus is located in the dorsal posterior PVLP (Wu et al., 2016). LC25 belong within cluster 57, another heterogeneous cluster (See Methods; Supp Figure 10), with the corresponding markers: Br (n= 8 Ol), Pdm3 (n= 6 Ol), D (n= 7 Ol), Pros (n= 6 Ol). They do not express: RunxA (n= 4 Ol), Scro (n= 6 Ol), Kn (n= 3 Ol), Acj6 (n= 5 Ol), Sox102F (n=5 Ol), Ct (n= 5 Ol), Lov (n= 4 Ol), Bab2 (n= 6 Ol), Rx (n= 5 Ol), Toy (n= 3 Ol).

#### LC15, LC16, LC24 and LC26/Cluster 100

LC15 cell bodies are located in the ventral LCBR with their optic glomerulus in the anterior PVLP (dorsal-lateral) adjacent to LC16 (Wu et al., 2016). Screening for LC15 markers revealed several hits, including: Ct (n=3 Ol), Fred (n= 4 Ol), Pdm3 (n= 4 Ol) (all antibody staining), and they were negative for Acj6 (n= 4 Ol), Toy (n= 4 Ol) and Rx (n= 5 Ol) (antibody staining); Kn (n= 6 OL) and Sns (3 OL) (MiMIC). This combination of markers is consistent with cluster 100 (Supp Figure 11). However, this particular cluster had been already identified as LC16 (Neset et al., 2021). Upon further examination, it was revealed that cluster 100 contains multiple subclusters (a-e) (Supp Figure 15), where LC15 makes up subcluster b (Scro negative (n= 4 Ol); Lov negative (n= 5 Ol)).

Thus, we set out to classify the identity of these subclusters. We went back to LC16 and checked several markers where they were positive for Lov (n= 4 Ol), Sox102F (n= 6 Ol), Pros (n= 6 Ol); and negative for Pdm3 (n= 4 Ol) and Scro (n= 4 Ol). Therefore, LC16 comprise subcluster a of cluster 100 (Supp Figure 12).

Additionally, we identified subcluster c as LC24 (Supp Figure 13). These have cell bodies in the ventral LCBR, with an optic glomerulus in posterior PVLP (posterior-dorsal) (Wu et al., 2016). LC24’s markers included: Sox102F (n= 6 Ol), Ct (n= 5 Ol), Scro (n= 6 Ol), Disco (n= 4 Ol) (all antibody staining). Furthermore, they do not express the following: Acj6 (n= 7 Ol), Rx (n= 4 Ol), Pdm3 (n= 7 Ol), Br (n= 6 Ol), Lov (n= 6 Ol), Mirr (n=3 Ol), Bab2 (n= 4 Ol), RunxA (n= 5 Ol), Pros (n= 3 Ol), D (n= 4 Ol), Kn (n= 5 Ol) (all antibody staining).

Lastly, here we show that subcluster d belongs to LC26 (Supp Figure 14). Similar to LC15, LC16 and LC24, LC26s cell bodies reside in the ventral LCBR with axons projecting to the dorsal posterior PVLP (Wu et al., 2016). LC26 express the following genes: Pdm3 (n= 6 Ol), Sox102F (n= 8 Ol), Scro (n= 3 Ol), Ct (n= 4 Ol), Pros (n= 3 Ol), whereas they are negative for: Acj6 (n= 4 Ol),Br (n= 6 Ol), Lov (n= 4 Ol), Bab2 (n= 4 Ol), D (n= 4 Ol), Toy (n= 4 Ol), Rx (n= 3 Ol), RunxA (n= 4 Ol).

Finally, we asked whether we could separate the original cluster 100 annotation into its constituents in our *wg-*Gal4 scRNAseq dataset which would further indicate a common origin for LC15, LC16, LC24 and LC26. We noticed that in our neural network classified dataset cells from cluster 100 mapped to a single UMAP trajectory with higher confidence. We ran unsupervised clustering on these cells and were able to separate this trajectory into two subclusters comprising of LC15/26 (Wnt4^+^ and Lov^+^) and LC16/24 (Pdm3^+^) respectively (Supp Figure 15B). Due to the highly similar transcriptomic identity of these cell types, we were unable to further separate the LC15/26 and LC16/24 subclusters in our late L3 stage *wg*-Gal4 scRNAseq dataset (Figure1C; Supp Figure 15B).

Lastly, subcluster e could not be labeled here, and its identity is yet to be determined. It didn’t fall under any known LCN subtype, as we have identified all the clusters for the previously published LCNs (Fischbach & Dittrich, 1989; Otsuna & Ito, 2006; Panser et al., 2016; Wu et al., 2016). They could belong to LCNs that haven’t been identified yet, or other lobula neurons which share similar genetic composition, cell body location or axonal trajectory to optic glomeruli similar to LC15, LC16, LC24 and LC26.

## References

Ache, J. M., Polsky, J., Alghailani, S., Parekh, R., Breads, P., Peek, M. Y., Bock, D. D., von Reyn, C. R., & Card, G. M. (2019). Neural Basis for Looming Size and Velocity Encoding in the Drosophila Giant Fiber Escape Pathway. Current Biology, 29(6), 1073–1081.e4. 10.1016/j.cub.2019.01.079

Apitz, H., & Salecker, I. (2015). A region-specific neurogenesis mode requires migratory progenitors in the Drosophila visual system. Nature Neuroscience, 18(1), 46–55. 10.1038/nn.3896

Apitz, H., & Salecker, I. (2018). Spatiooral relays control layer identity of direction-selective neuron subtypes in Drosophila. Nature Communications, 9(1). 10.1038/s41467-018-04592-z

Aptekar, J. W., Keleş, M. F., Lu, P. M., Zolotova, N. M., & Frye, M. A. (2015). Neurons forming optic glomeruli compute figure–ground discriminations in Drosophila. Journal of Neuroscience, 35(19), 7587–7599. 10.1523/JNEUROSCI.0652-15.2015

Bertet, C., Li, X., Erclik, T., Cavey, M., Wells, B., & Desplan, C. (2014). Temporal patterning of neuroblasts controls notch-mediated cell survival through regulation of hid or reaper. Cell, 158(5), 1173–1186. 10.1016/j.cell.2014.07.045

Cepko, C. (2014). Intrinsically different retinal progenitor cells produce specific types of progeny. In Nature Reviews Neuroscience (Vol. 15, Issue 9, pp. 615–627). Nature Publishing Group. 10.1038/nrn3767

Chen, Y. C. D., Chen, Y. C., Rajesh, R., Shoji, N., Jacy, M., Lacin, H., Erclik, T., & Desplan, C. (2023). Using single-cell RNA sequencing to generate predictive cell-type-specific split-GAL4 reagents throughout development. Proceedings of the National Academy of Sciences of the United States of America, 120(32). 10.1073/pnas.2307451120

Cruz-Martín, A., El-Danaf, R. N., Osakada, F., Sriram, B., Dhande, O. S., Nguyen, P. L., Callaway, E. M., Ghosh, A., & Huberman, A. D. (2014). A dedicated circuit links direction-selective retinal ganglion cells to the primary visual cortex. Nature, 507(7492), 358–361. 10.1038/nature12989

Davis, F. P., Nern, A., Picard, S., Reiser, M. B., Rubin, G. M., Eddy, S. R., & Henry, G. L. (2020). A genetic, genomic, and computational resource for exploring neural circuit function. ELife, 9. 10.7554/eLife.50901

Dearbon, R., & Kunes, S. (2004). An axon scaffold induced by retinal axons directs glia to destinations in the Drosophila optic lobe. Development, 131(10), 2291–2303. 10.1242/dev.01111

Dhande, O. S., Stafford, B. K., Lim, J.-H. A., & Huberman, A. D. (2015). Contributions of Retinal Ganglion Cells to Subcortical Visual Processing and Behaviors. Annual Review of Vision Science, 1(1), 291–328. 10.1146/annurev-vision-082114-035502

Doe, C. Q. (2017). Temporal Patterning in the Drosophila CNS. Annual Review of Cell and Developmental Biology, 12, 55. 10.1146/annurev-cellbio-111315

El-Danaf, R. N., Rajesh, R., & Desplan, C. (2023). Temporal regulation of neural diversity in Drosophila and vertebrates. In Seminars in Cell and Developmental Biology (Vol. 142, pp. 13–22). Elsevier Ltd. 10.1016/j.semcdb.2022.05.011

Erclik, T., Hartenstein, V., Lipshitz, H. D., & McInnes, R. R. (2008). Conserved Role of the Vsx Genes Supports a Monophyletic Origin for Bilaterian Visual Systems. Current Biology, 18(17), 1278–1287. 10.1016/j.cub.2008.07.076

Erclik, T., Li, X., Courgeon, M., Bertet, C., Chen, Z., Baumert, R., Ng, J., Koo, C., Arain, U., Behnia, R., Del Valle Rodriguez, A., Senderowicz, L., Negre, N., White, K. P., & Desplan, C. (2017). Integration of temporal and spatial patterning generates neural diversity. Nature, 541(7637), 365–370. 10.1038/nature20794

Farrow, K., Teixeira, M., Szikra, T., Viney, T. J., Balint, K., Yonehara, K., & Roska, B. (2013). Ambient illumination toggles a neuronal circuit switch in the retina and visual perception at cone threshold. Neuron, 78(2), 325–338. 10.1016/j.neuron.2013.02.014

Fischbach, K.-F., & Dittrich, A. P. M. (1989). Cell Tissue Res (1989) 258:441M75 The optic lobe of Drosophila melanogaster. I. A Golgi analysis of wild-type structure. Cell and Tissue Research, 258, 441–475.

Galindo, M. I., Fernández-Garza, D., Phillips, R., & Couso, J. P. (2011). Control of Distal-less expression in the Drosophila appendages by functional 3’ enhancers. Developmental Biology, 353(2), 396–410. 10.1016/j.ydbio.2011.02.005

Gao, Q., & Chess, A. (1999). Identification of Candidate Drosophila Olfactory Receptors from Genomic DNA Sequence. Genomics, 60, 31–39. http://CCR-081.mit.edu/GENSCAN.html

Gao, Q., Yuan, B., & Chess, A. (2000). Convergent projections of Drosophila olfactory neurons to specific glomeruli in the antennal lobe. Nature Neuroscience, 3(8), 780– 785. http://neurosci.nature.com

Gold, K. S., & Brand, A. H. (2014). Optix defines a neuroepithelial compartment in the optic lobe of the Drosophila brain. Neural Development, 9(1). 10.1186/1749-8104-9-18

Hahn, J., Monavarfeshani, A., Qiao, M., Kao, A., Kölsch, Y., Kumar, A., Kunze, V. P., Rasys, A. M., Richardson, R., Baier, H., Lucas, R. J., Li, W., Meister, M., Trachtenberg, J. T., Yan, W., Peng, Y.-R., Sanes, J. R., & Shekhar, K. (2023). Evolution of neuronal cell classes and types in the vertebrate retina. BioRxiv. 10.1101/2023.04.07.536039

Holguera, I., & Desplan, C. (2018). Neuronal specification in space and time. Science, 362, 176–180. http://science.sciencemag.org/

Islam, I. M., Ng, J., Valentino, P., & Erclik, T. (2021). Identification of enhancers that drive the spatially restricted expression of vsx1 and rx in the outer proliferation center of the developing drosophila optic lobe. Genome, 64(2), 109–117. 10.1139/gen-2020-0034

Jenett, A., Rubin, G. M., Ngo, T. T. B., Shepherd, D., Murphy, C., Dionne, H., Pfeiffer, B. D., Cavallaro, A., Hall, D., Jeter, J., Iyer, N., Fetter, D., Hausenfluck, J. H., Peng, H., Trautman, E. T., Svirskas, R. R., Myers, E. W., Iwinski, Z. R., Aso, Y., … Zugates, C. T. (2012). A GAL4-Driver Line Resource for Drosophila Neurobiology. Cell Reports, 2(4), 991–1001. 10.1016/j.celrep.2012.09.011

Jhaveri, D., Sen, A., & Rodrigues, V. (2000). Mechanisms underlying olfactory neuronal connectivity in Drosophila - The atonal lineage organizes the periphery while sensory neurons and glia pattern the olfactory lobe. Developmental Biology, 226(1), 73–87. 10.1006/dbio.2000.9855

Keleş, M. F., & Frye, M. A. (2017). Object-Detecting Neurons in Drosophila. Current Biology, 27(5), 680–687. 10.1016/j.cub.2017.01.012

Keleş, M. F., Hardcastle, B. J., Städele, C., Xiao, Q., & Frye, M. A. (2020). Inhibitory Interactions and Columnar Inputs to an Object Motion Detector in Drosophila. Cell Reports, 30(7), 2115–2124.e5. 10.1016/j.celrep.2020.01.061

Klapoetke, N. C., Nern, A., Peek, M. Y., Rogers, E. M., Breads, P., Rubin, G. M., Reiser, M. B., & Card, G. M. (2017). Ultra-selective looming detection from radial motion opponency. Nature, 551(7679), 237–241. 10.1038/nature24626

Klapoetke, N. C., Nern, A., Rogers, E. M., Rubin, G. M., Reiser, M. B., & Card, G. M. (2022). A functionally ordered visual feature map in the Drosophila brain. Neuron, 110(10), 1700–1711.e6. 10.1016/j.neuron.2022.02.013

Konstantinides, N., Holguera, I., Rossi, A. M., Escobar, A., Dudragne, L., Chen, Y. C., Tran, T. N., Martínez Jaimes, A. M., Özel, M. N., Simon, F., Shao, Z., Tsankova, N. M., Fullard, J. F., Walldorf, U., Roussos, P., & Desplan, C. (2022). A complete temporal transcription factor series in the fly visual system. Nature, 604(7905), 316–322. 10.1038/s41586-022-04564-w

Konstantinides, N., Kapuralin, K., Fadil, C., Barboza, L., Satija, R., & Desplan, C. (2018). Phenotypic Convergence: Distinct Transcription Factors Regulate Common Terminal Features. Cell, 174(3), 622–635.e13. 10.1016/j.cell.2018.05.021

Konstantinides, N., Rossi, A. M., & Desplan, C. (2015). Common temporal identity factors regulate neuronal diversity in fly ventral nerve cord and mouse retina. In Neuron (Vol. 85, Issue 3, pp. 447–449). Cell Press. 10.1016/j.neuron.2015.01.016

Kurmangaliyev, Y. Z., Yoo, J., Valdes-Aleman, J., Sanfilippo, P., & Zipursky, S. L. (2020). Transcriptional Programs of Circuit Assembly in the Drosophila Visual System. Neuron, 108(6), 1045–1057.e6. 10.1016/j.neuron.2020.10.006

Laissue, P. P., Reiter, C., Hiesinger, P. R., Halter, S., Fischbach, K. F., & Stocker, R. F. (1999). Three-dimensional reconstruction of the antennal lobe in Drosophila melanogaster. Journal of Comparative Neurology, 405(4), 543–552. 10.1002/(SICI)1096-9861(19990322)405:4<543::AID-CNE7>3.0.CO;2-A

Li, X., Erclik, T., Bertet, C., Chen, Z., Voutev, R., Venkatesh, S., Morante, J., Celik, A., & Desplan, C. (2013). Temporal patterning of Drosophila medulla neuroblasts controls neural fates. Nature, 498(7455), 456–462. 10.1038/nature12319

Malin, J. A., Chen, Y.-C., Simon, F., Keefer, E., & Desplan, C. (2023). Spatial patterning regulates neuron numbers in the Drosophila visual system 1. BiorXiv. 10.1101/2023.08.28.555170

Morimoto, M. M., Nern, A., Zhao, A., Rogers, E. M., Wong, A. M., Isaacson, M. D., Bock, D. D., Rubin, G. M., & Reiser, M. B. (2020). Spatial readout of visual looming in the central brain of drosophila. ELife, 9, 1–102. 10.7554/eLife.57685

Münch, T. A., Da Silveira, R. A., Siegert, S., Viney, T. J., Awatramani, G. B., & Roska, B. (2009). Approach sensitivity in the retina processed by a multifunctional neural circuit. Nature Neuroscience, 12(10), 1308–1316. 10.1038/nn.2389

Nagarkar-Jaiswal, S., DeLuca, S. Z., Lee, P.-T., Lin, W.-W., Pan, H., Zuo, Z., Lv, J., Spradling, A. C., & Bellen, H. J. (2016). A genetic toolkit for tagging intronic MiMIC containing genes. ELIFE. 10.7554/eLife.08469.001

Ngo, K. T., Andrade, I., & Hartenstein, V. (2017). Spatio-temporal pattern of neuronal differentiation in the Drosophila visual system: A user’s guide to the dynamic morphology of the developing optic lobe. Developmental Biology, 428(1), 1–24. 10.1016/j.ydbio.2017.05.008

Nguyen-ba-charvet, K. T., & Rebsam, A. (2020). Neurogenesis and specification of retinal ganglion cells. In International Journal of Molecular Sciences (Vol. 21, Issue 2). MDPI AG. 10.3390/ijms21020451

Oliva, C., Choi, C. M., Nicolai, L. J. J., Mora, N., De Geest, N., & Hassan, B. A. (2014). Proper connectivity of Drosophila motion detector neurons requires Atonal function in progenitor cells. Neural Development, 9(1). 10.1186/1749-8104-9-4

Osterhout, J. A., El-Danaf, R. N., Nguyen, P. L., & Huberman, A. D. (2014). Birthdate and outgrowth timing predict cellular mechanisms of axon target matching in the developing visual pathway. Cell Reports, 8(4), 1006–1017. 10.1016/j.celrep.2014.06.063

Otsuna, H., & Ito, K. (2006). Systematic analysis of the visual projection neurons of Drosophila melanogaster. I. Lobula-specific pathways. Journal of Comparative Neurology, 497(6), 928–958. 10.1002/cne.21015

Özel, M. N., Gibbs, C. S., Holguera, I., Soliman, M., Bonneau, R., & Desplan, C. (2022). Coordinated control of neuronal differentiation and wiring by sustained transcription factors. Science, 378(6626). 10.1126/science.add1884

Özel, M. N., Simon, F., Jafari, S., Holguera, I., Chen, Y. C., Benhra, N., El-Danaf, R. N., Kapuralin, K., Malin, J. A., Konstantinides, N., & Desplan, C. (2021). Neuronal diversity and convergence in a visual system developmental atlas. Nature, 589(7840), 88–95. 10.1038/s41586-020-2879-3

Panser, K., Tirian, L., Schulze, F., Villalba, S., Jefferis, G. S. X. E., Bühler, K., & Straw, A. D. (2016). Automatic Segmentation of Drosophila Neural Compartments Using GAL4 Expression Data Reveals Novel Visual Pathways. Current Biology, 26(15), 1943– 1954. 10.1016/j.cub.2016.05.052

Perez, S. E., & Steller, H. (1996). Migration of Glial Cells into Retinal Axon Target Field in Drosophila melanogaster. Journal of Neurobiology, 30(3), 359–373.

Pinto-Teixeira, F., Koo, C., Rossi, A. M., Neriec, N., Bertet, C., Li, X., Del-Valle-Rodriguez, A., & Desplan, C. (2018). Development of Concurrent Retinotopic Maps in the Fly Motion Detection Circuit. Cell, 173(2), 485–498.e11. 10.1016/j.cell.2018.02.053

Rheaume, B. A., Jereen, A., Bolisetty, M., Sajid, M. S., Yang, Y., Renna, K., Sun, L., Robson, P., & Trakhtenberg, E. F. (2018). Single cell transcriptome profiling of retinal ganglion cells identifies cellular subtypes. Nature Communications, 9(1). 10.1038/s41467-018-05134-3

Ribeiro, I. M. A., Drews, M., Bahl, A., Machacek, C., Borst, A., & Dickson, B. J. (2018). Visual Projection Neurons Mediating Directed Courtship in Drosophila. Cell, 174(3), 607–621.e18. 10.1016/j.cell.2018.06.020

Sanes, J. R., & Masland, R. H. (2015). The Types of Retinal Ganglion Cells: Current Status and Implications for Neuronal Classification. Annual Review of Neuroscience, 38, 221–246. 10.1146/annurev-neuro-071714-034120

Sato, M., Suzuki, T., & Nakai, Y. (2013). Waves of differentiation in the fly visual system. In Developmental Biology (Vol. 380, Issue 1, pp. 1–11). Academic Press Inc. 10.1016/j.ydbio.2013.04.007

Sato, M., Yasugi, T., & Trush, O. (2019). Temporal patterning of neurogenesis and neural wiring in the fly visual system. In Neuroscience Research (Vol. 138, pp. 49–58). Elsevier Ireland Ltd. 10.1016/j.neures.2018.09.009

Scott, K., Wu, M., Nern, A., Williamson, R., Morimoto, M. M., Reiser, M. B., Card, G. M., & Rubin, G. M. (2016). Visual projection neurons in the Drosophila lobula link feature detection to distinct behavioral programs. ELIFE. 10.7554/eLife.21022.001

Seabrook, T. A., Burbridge, T. J., Crair, M. C., & Huberman, A. D. (2017). Architecture, Function, and Assembly of the Mouse Visual System. Annual Review Neuroscience, 40, 499–538. 10.1146/annurev-neuro-071714

Sen, R., Wu, M., Branson, K., Robie, A., Rubin, G. M., & Dickson, B. J. (2017). Moonwalker Descending Neurons Mediate Visually Evoked Retreat in Drosophila. Current Biology, 27(5), 766–771. 10.1016/j.cub.2017.02.008

Shekhar, K., Whitney, I. E., Butrus, S., Peng, Y. R., & Sanes, J. R. (2022). Diversification of multipotential postmitotic mouse retinal ganglion cell precursors into discrete types. ELife, 11. 10.7554/ELIFE.73809

Städele, C., Keleş, M. F., Mongeau, J. M., & Frye, M. A. (2020). Non-canonical Receptive Field Properties and Neuromodulation of Feature-Detecting Neurons in Flies. Current Biology, 30(13), 2508–2519.e6. 10.1016/j.cub.2020.04.069

Sten, T. H., Li, R., Otopalik, A., & Ruta, V. (2021). Sexual arousal gates visual processing during Drosophila courtship. Nature, 595(7868), 549–553. 10.1038/s41586-021-03714-w

Suzuki, T., Kaido, M., Takayama, R., & Sato, M. (2013). A temporal mechanism that produces neuronal diversity in the Drosophila visual center. Developmental Biology, 380(1), 12–24. 10.1016/j.ydbio.2013.05.002

Tanaka, R., & Clark, D. A. (2020). Object-Displacement-Sensitive Visual Neurons Drive Freezing in Drosophila. Current Biology, 30(13), 2532–2550.e8. 10.1016/j.cub.2020.04.068

Tanaka, R., & Clark, D. A. (2022). Identifying Inputs to Visual Projection Neurons in Drosophila Lobula by Analyzing Connectomic Data. ENeuro, 9(2). 10.1523/ENEURO.0053-22.2022

Timaeus, L., Geid, L., Sancer, G., Wernet, M. F., & Hummel, T. (2020). Parallel Visual Pathways with Topographic versus Nontopographic Organization Connect the Drosophila Eyes to the Central Brain. IScience, 23(10). 10.1016/j.isci.2020.101590

Valentino, P., & Erclik, T. (2022). Spalt and disco define the dorsal-ventral neuroepithelial compartments of the developing Drosophila medulla. Genetics, 222(3). 10.1093/genetics/iyac145

von Reyn, C. R., Nern, A., Williamson, W. R., Breads, P., Wu, M., Namiki, S., & Card, G. M. (2017). Feature Integration Drives Probabilistic Behavior in the Drosophila Escape Response. Neuron, 94(6), 1190–1204.e6. 10.1016/j.neuron.2017.05.036

Vosshall, L. B., Wong, A. M., & Axel, R. (2000). An Olfactory Sensory Map in the Fly Brain. Cell, 102, 147–159.

West, E. R., Lapan, S. W., Lee, C. H., Kajderowicz, K. M., Li, X., & Cepko, C. L. (2022). Spatiotemporal patterns of neuronal subtype genesis suggest hierarchical development of retinal diversity. Cell Reports, 38(1). 10.1016/j.celrep.2021.110191

Wu, M., Nern, A., Williamson, R., Morimoto, M. M., Reiser, M. B., Card, G. M., & Rubin, G. M. (2016). Visual projection neurons in the Drosophila lobula link feature detection to distinct behavioral programs. ELife. 10.7554/eLife.21022.001

Yilmaz, M., & Meister, M. (2013). Rapid innate defensive responses of mice to looming visual stimuli. Current Biology, 23(20), 2011–2015. 10.1016/j.cub.2013.08.015

Yu, H. H., Awasaki, T., Schroeder, M. D., Long, F., Yang, J. S., He, Y., Ding, P., Kao, J. C., Wu, G. Y. Y., Peng, H., Myers, G., & Lee, T. (2013). Clonal development and organization of the adult Drosophila central brain. Current Biology, 23(8), 633–643. 10.1016/j.cub.2013.02.057

Zhang, Y., Kim, I. J., Sanes, J. R., & Meister, M. (2012). The most numerous ganglion cell type of the mouse retina is a selective feature detector. Proceedings of the National Academy of Sciences of the United States of America, 109(36), E2391–E2398. 10.1073/pnas.1211547109

Zhu, H., Zhao, S. D., Ray, A., Zhang, Y., & Li, X. (2022). A comprehensive temporal patterning gene network in Drosophila medulla neuroblasts revealed by single-cell RNA sequencing. Nature Communications, 13(1). 10.1038/s41467-022-28915-3

