## Supplementary material for "Morphological and functional convergence of visual projections neurons from diverse neurogenic origins in *Drosophila*": Annex 1

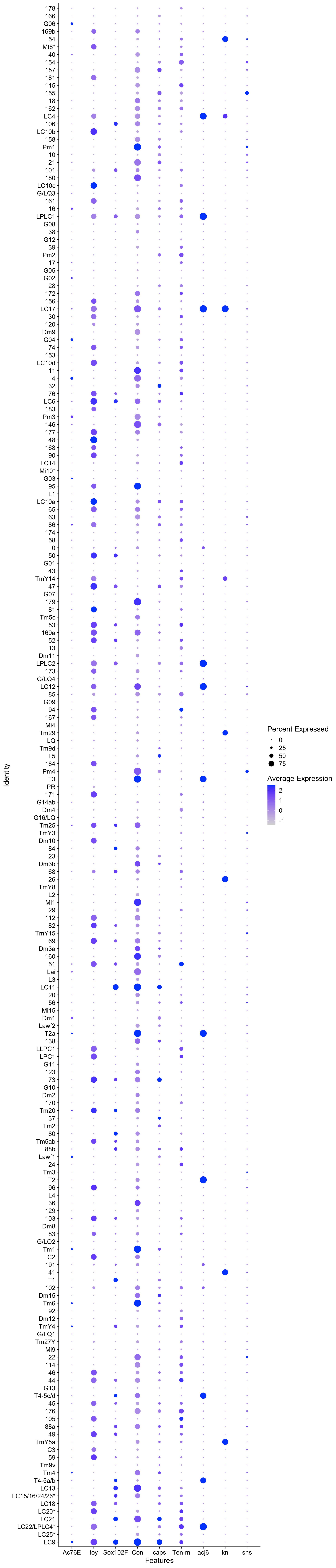

Percent Expressed
0
25
50
75
Average Expression
2
1
0
-1
Identity
Features
Ac76E
toy
Sox102F
Con
caps
Ten-m
acj6
kn
sns

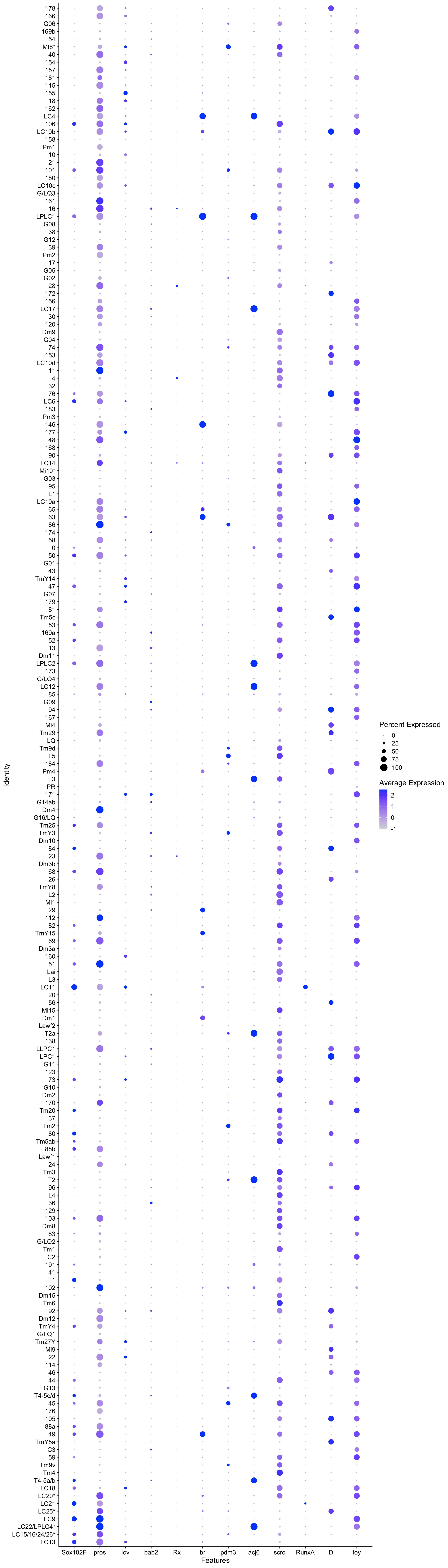

Identity
178
166
G06
169b
54
Mt8*
40
154
157
181
115
155
18
162
LC4
106
LC10b
158
Pm1
10
21
101
180
LC10c
G/LQ3
161
16
LPLC1
G08
38
G12
39
Pm2
17
G05
G02
28
172
156
LC17
30
120
Dm9
G04
74
153
LC10d
11
4
32
76
LC6
183
Pm3
146
177
48
168
90
LC14
Mi10*
G03
95
L1
LC10a
65
63
86
174
58
0
50
G01
43
TmY14
47
G07
179
81
Tm5c
53
169a
52
13
Dm11
LPLC2
173
G/LQ4
LC12
85
G09
94
167
Mi4
Tm29
LQ
Tm9d
L5
184
Pm4
T3
PR
171
G14ab
Dm4
G16/LQ
Tm25
TmY3
Dm10
84
23
Dm3b
68
26
TmY8
L2
Mi1
29
112
82
TmY15
69
Dm3a
160
51
Lai
L3
LC11
20
56
Mi15
Dm1
Lawf2
T2a
138
LLPC1
LPC1
G11
123
73
G10
Dm2
170
Tm20
37
Tm2
80
Tm5ab
88b
Lawf1
24
Tm3
T2
96
L4
36
129
103
Dm8
83
G/LQ2
Tm1
C2
191
41
T1
102
Dm15
Tm6
92
Dm12
TmY4
G/LQ1
Tm27Y
Mi9
22
114
46
44
G13
T4-5c/d
45
176
105
88a
49
TmY5a
C3
59
Tm9v
Tm4
T4-5a/b
LC18
LC20*
LC21
LC25*
LC9
LC22/LPLC4*
LC15/16/24/26*
LC13
Sox102F
pros
lov
bab2
Rx
br
pdm3
acj6
scro
RunxA
D
toy
Features
Percent Expressed
0
25
50
75
100
Average Expression
2
1
0
-1

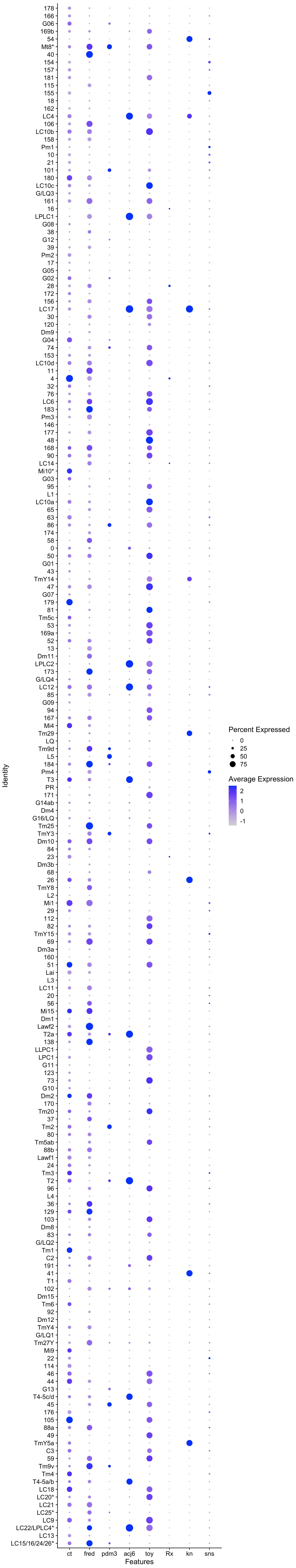
178
166
G06
169b
54
Mt8*
40
154
157
181
115
155
18
162
LC4
106
LC10b
158
Pm1
10
21
101
180
LC10c
G/LQ3
161
16
LPLC1
G08
38
G12
39
Pm2
17
G05
G02
28
172
156
LC17
30
120
Dm9
G04
74
153
LC10d
11
4
32
76
LC6
183
Pm3
146
177
48
168
90
LC14
Mi10*
G03
95
L1
LC10a
65
63
86
174
58
0
50
G01
43
TmY14
47
G07
179
81
Tm5c
53
169a
52
13
Dm11
LPLC2
173
G/LQ4
LC12
85
G09
94
167
Mi4
Tm29
LQ
Tm9d
L5
184
Pm4
T3
PR
171
G14ab
Dm4
G16/LQ
Tm25
TmY3
Dm10
84
23
Dm3b
68
26
TmY8
L2
Mi1
29
112
82
TmY15
69
Dm3a
160
51
Lai
L3
LC11
20
56
Mi15
Dm1
Lawf2
T2a
138
LLPC1
LPC1
G11
123
73
G10
Dm2
170
Tm20
37
Tm2
80
Tm5ab
88b
Lawf1
24
Tm3
T2
96
L4
36
129
103
Dm8
83
G/LQ2
Tm1
C2
191
41
T1
102
Dm15
Tm6
92
Dm12
TmY4
G/LQ1
Tm27Y
Mi9
22
114
46
44
G13
T4-5c/d
45
176
105
88a
49
TmY5a
C3
59
Tm9v
Tm4
T4-5a/b
LC18
LC20*
LC21
LC25*
LC9
LC22/LPLC4*
LC13
LC15/16/24/26*
Identity
ct
fred
pdm3
acj6
toy
Rx
kn
sns
Features
Percent Expressed
0
25
50
75
Average Expression
2
1
0
-1

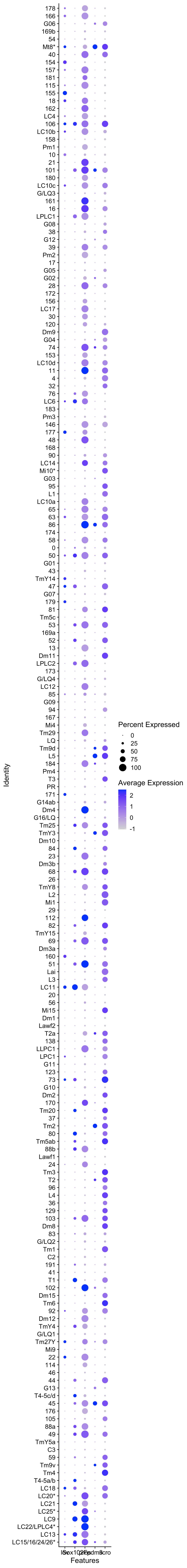

178
166
G06
169b
54
Mt8*
40
154
157
181
115
155
18
162
LC4
106
LC10b
158
Pm1
10
21
101
180
LC10c
G/LQ3
161
16
LPLC1
G08
38
G12
39
Pm2
17
G05
G02
28
172
156
LC17
30
120
Dm9
G04
74
153
LC10d
11
4
32
76
LC6
183
Pm3
146
177
48
168
90
LC14
Mi10*
G03
95
L1
LC10a
65
63
86
174
58
0
50
G01
43
TmY14
47
G07
179
81
Tm5c
53
169a
52
13
Dm11
LPLC2
173
G/LQ4
LC12
85
G09
94
167
Mi4
Tm29
LQ
Tm9d
L5
184
Pm4
T3
PR
171
G14ab
Dm4
G16/LQ
Tm25
TmY3
Dm10
84
23
Dm3b
68
26
TmY8
L2
Mi1
29
112
82
TmY15
69
Dm3a
160
51
Lai
L3
LC11
20
56
Mi15
Dm1
Lawf2
T2a
138
LLPC1
LPC1
G11
123
73
G10
Dm2
170
Tm20
37
Tm2
80
Tm5ab
88b
Lawf1
24
Tm3
T2
96
L4
36
129
103
Dm8
83
G/LQ2
Tm1
C2
191
41
T1
102
Dm15
Tm6
92
Dm12
TmY4
G/LQ1
Tm27Y
Mi9
22
114
46
44
G13
T4-5c/d
45
176
105
88a
49
TmY5a
C3
59
Tm9v
Tm4
T4-5a/b
LC18
LC20*
LC21
LC25*
LC9
LC22/LPLC4*
LC13
LC15/16/24/26*
Identity
Features
Percent Expressed
0
25
50
75
100
Average Expression
2
1
0
-1

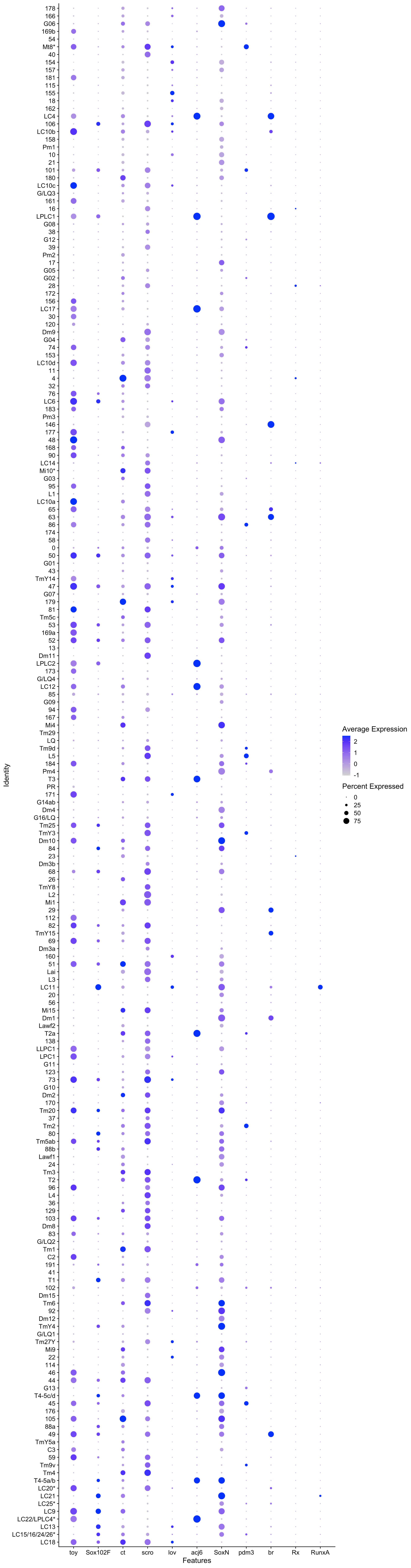
Identity
Features
toy
Sox102F
ct
scro
lov
acj6
SoxN
pdm3
br
Rx
RunxA
Average Expression
Percent Expressed
0
25
50
75

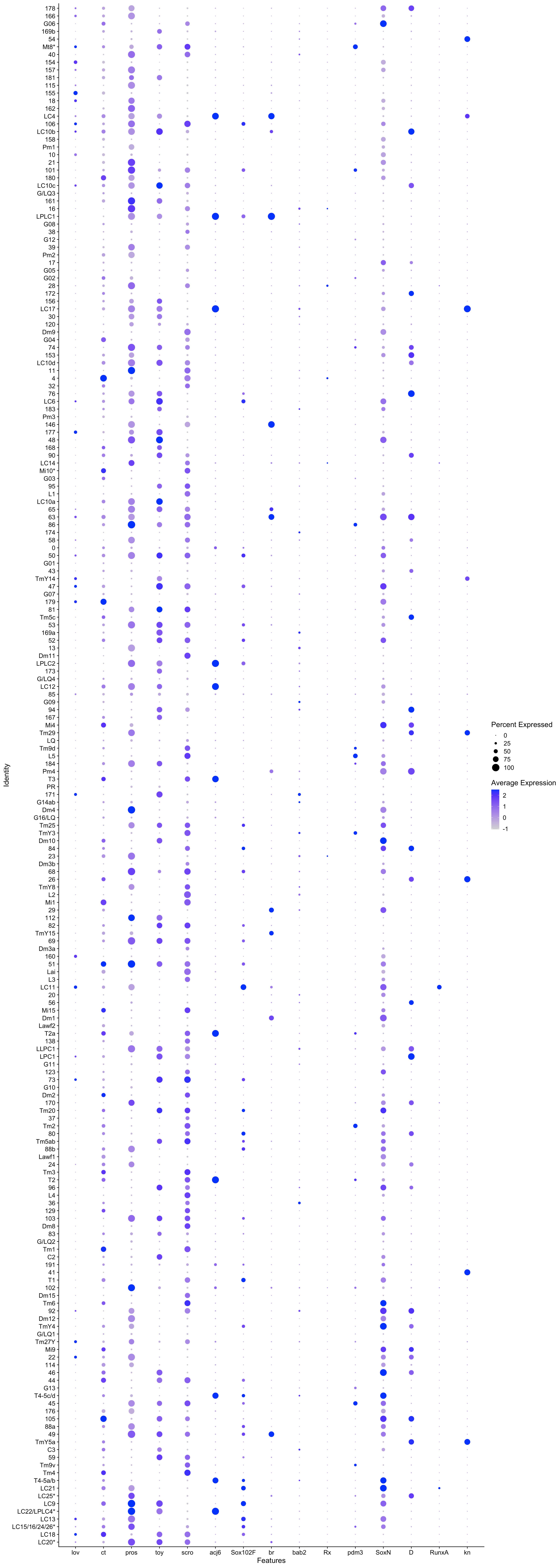
Identity
178
166
G06
169b
54
Mt8*
40
154
157
181
115
155
18
162
LC4
106
LC10b
158
Pm1
10
21
101
180
LC10c
G/LQ3
161
16
LPLC1
G08
38
G12
39
Pm2
17
G05
G02
28
172
156
LC17
30
120
Dm9
G04
74
153
LC10d
11
4
32
76
LC6
183
Pm3
146
177
48
168
90
LC14
Mi10*
G03
95
L1
LC10a
65
63
86
174
58
0
50
G01
43
TmY14
47
G07
179
81
Tm5c
53
169a
52
13
Dm11
LPLC2
173
G/LQ4
LC12
85
G09
94
167
Mi4
Tm29
LQ
Tm9d
L5
184
Pm4
T3
PR
171
G14ab
Dm4
G16/LQ
Tm25
TmY3
Dm10
84
23
Dm3b
68
26
TmY8
L2
Mi1
29
112
82
TmY15
69
Dm3a
160
51
Lai
L3
LC11
20
56
Mi15
Dm1
Lawf2
T2a
138
LLPC1
LPC1
G11
123
73
G10
Dm2
170
Tm20
37
Tm2
80
Tm5ab
88b
Lawf1
24
Tm3
T2
96
L4
36
129
103
Dm8
83
G/LQ2
Tm1
C2
191
41
T1
102
Dm15
Tm6
92
Dm12
TmY4
G/LQ1
Tm27Y
Mi9
22
114
46
44
G13
T4-5c/d
45
176
105
88a
49
TmY5a
C3
59
Tm9v
Tm4
T4-5a/b
LC21
LC25*
LC9
LC22/LPLC4*
LC13
LC15/16/24/26*
LC18
LC20*
lov
ct
pros
toy
scro
acj6
Sox102F
br
bab2
Rx
pdm3
SoxN
D
RunxA
kn
Features
Percent Expressed
0
25
50
75
100
Average Expression
2
1
0
-1

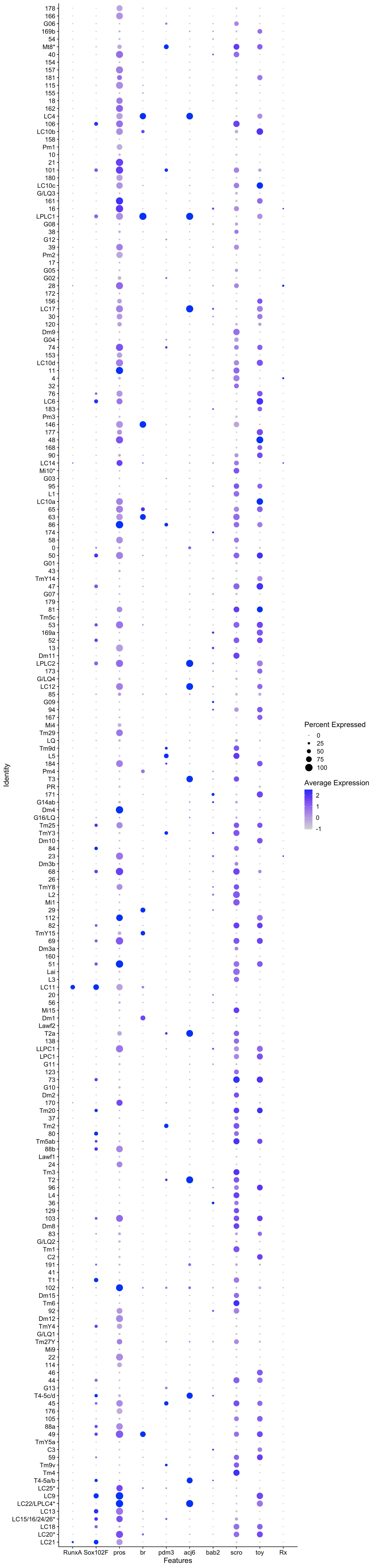
178
166
G06
169b
54
Mt8*
40
154
157
181
115
155
18
162
LC4
106
LC10b
158
Pm1
10
21
101
180
LC10c
G/LQ3
161
16
LPLC1
G08
38
G12
39
Pm2
17
G05
G02
28
172
156
LC17
30
120
Dm9
G04
74
153
LC10d
11
4
32
76
LC6
183
Pm3
146
177
48
168
90
LC14
Mi10*
G03
95
L1
LC10a
65
63
86
174
58
0
50
G01
43
TmY14
47
G07
179
81
Tm5c
53
169a
52
13
Dm11
LPLC2
173
G/LQ4
LC12
85
G09
94
167
Mi4
Tm29
LQ
Tm9d
L5
184
Pm4
T3
PR
171
G14ab
Dm4
G16/LQ
Tm25
TmY3
Dm10
84
23
Dm3b
68
26
TmY8
L2
Mi1
29
112
82
TmY15
69
Dm3a
160
51
Lai
L3
LC11
20
56
Mi15
Dm1
Lawf2
T2a
138
LLPC1
LPC1
G11
123
73
G10
Dm2
170
Tm20
37
Tm2
80
Tm5ab
88b
Lawf1
24
Tm3
T2
96
L4
36
129
103
Dm8
83
G/LQ2
Tm1
C2
191
41
T1
102
Dm15
Tm6
92
Dm12
TmY4
G/LQ1
Tm27Y
Mi9
22
114
46
44
G13
T4-5c/d
45
176
105
88a
49
TmY5a
C3
59
Tm9v
Tm4
T4-5a/b
LC25*
LC9
LC22/LPLC4*
LC13
LC15/16/24/26*
LC18
LC20*
LC21
Identity
RunxA
Sox102F
pros
br
pdm3
acj6
bab2
scro
toy
Rx
Features
Percent Expressed
0
25
50
75
100
Average Expression
2
1
0
-1

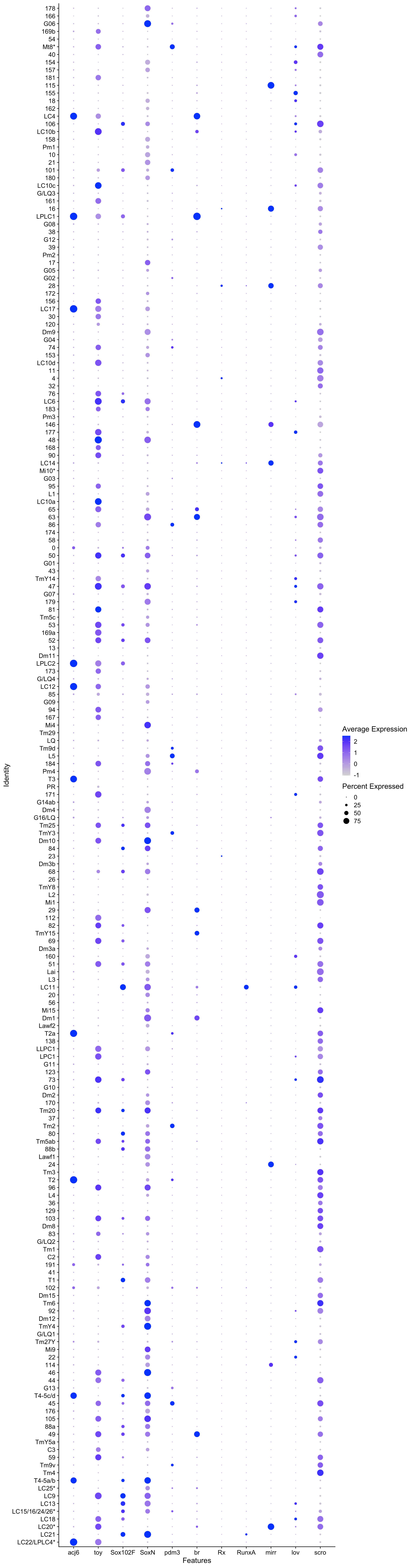
178
166
G06
169b
54
Mt8*
40
154
157
181
115
155
18
162
LC4
106
LC10b
158
Pm1
10
21
101
180
LC10c
G/LQ3
161
16
LPLC1
G08
38
G12
39
Pm2
17
G05
G02
28
172
156
LC17
30
120
Dm9
G04
74
153
LC10d
11
4
32
76
LC6
183
Pm3
146
177
48
168
90
LC14
Mi10*
G03
95
L1
LC10a
65
63
86
174
58
0
50
G01
43
TmY14
47
G07
179
81
Tm5c
53
169a
52
13
Dm11
LPLC2
173
G/LQ4
LC12
85
G09
94
167
Mi4
Tm29
LQ
Tm9d
L5
184
Pm4
T3
PR
171
G14ab
Dm4
G16/LQ
Tm25
TmY3
Dm10
84
23
Dm3b
68
26
TmY8
L2
Mi1
29
112
82
TmY15
69
Dm3a
160
51
Lai
L3
LC11
20
56
Mi15
Dm1
Lawf2
T2a
138
LLPC1
LPC1
G11
123
73
G10
Dm2
170
Tm20
37
Tm2
80
Tm5ab
88b
Lawf1
24
Tm3
T2
96
L4
36
129
103
Dm8
83
G/LQ2
Tm1
C2
191
41
T1
102
Dm15
Tm6
92
Dm12
TmY4
G/LQ1
Tm27Y
Mi9
22
114
46
44
G13
T4-5c/d
45
176
105
88a
49
TmY5a
C3
59
Tm9v
Tm4
T4-5a/b
LC25*
LC9
LC13
LC15/16/24/26*
LC18
LC20*
LC21
LC22/LPLC4*
Identity
acj6
toy
Sox102F
SoxN
pdm3
br
Rx
RunxA
mirr
lov
scro
Features
Average Expression
2
1
0
-1
Percent Expressed
0
25
50
75

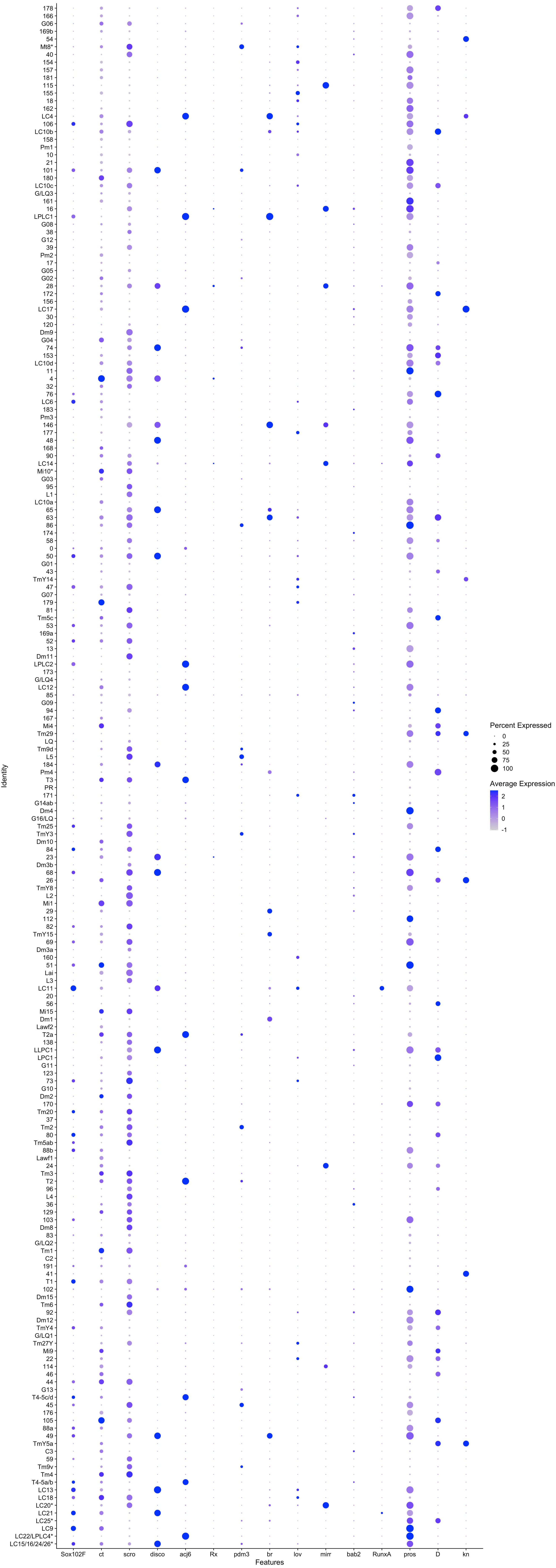

Identity
178
166
G06
169b
54
Mt8*
40
154
157
181
115
155
18
162
LC4
106
LC10b
158
Pm1
10
21
101
180
LC10c
G/LQ3
161
16
LPLC1
G08
38
G12
39
Pm2
17
G05
G02
28
172
156
LC17
30
120
Dm9
G04
74
153
LC10d
11
4
32
76
LC6
183
Pm3
146
177
48
168
90
LC14
Mi10*
G03
95
L1
LC10a
65
63
86
174
58
0
50
G01
43
TmY14
47
G07
179
81
Tm5c
53
169a
52
13
Dm11
LPLC2
173
G/LQ4
LC12
85
G09
94
167
Mi4
Tm29
LQ
Tm9d
L5
184
Pm4
T3
PR
171
G14ab
Dm4
G16/LQ
Tm25
TmY3
Dm10
84
23
Dm3b
68
26
TmY8
L2
Mi1
29
112
82
TmY15
69
Dm3a
160
51
Lai
L3
LC11
20
56
Mi15
Dm1
Lawf2
T2a
138
LLPC1
LPC1
G11
123
73
G10
Dm2
170
Tm20
37
Tm2
80
Tm5ab
88b
Lawf1
24
Tm3
T2
96
L4
36
129
103
Dm8
83
G/LQ2
Tm1
C2
191
41
T1
102
Dm15
Tm6
92
Dm12
TmY4
G/LQ1
Tm27Y
Mi9
22
114
46
44
G13
T4-5c/d
45
176
105
88a
49
TmY5a
C3
59
Tm9v
Tm4
T4-5a/b
LC13
LC18
LC20*
LC21
LC25*
LC9
LC22/LPLC4*
LC15/16/24/26*
Sox102F
ct
scro
disco
acj6
Rx
pdm3
br
lov
mirr
bab2
RunxA
pros
D
kn
Features
Percent Expressed
0
25
50
75
100
Average Expression
2
1
0
-1

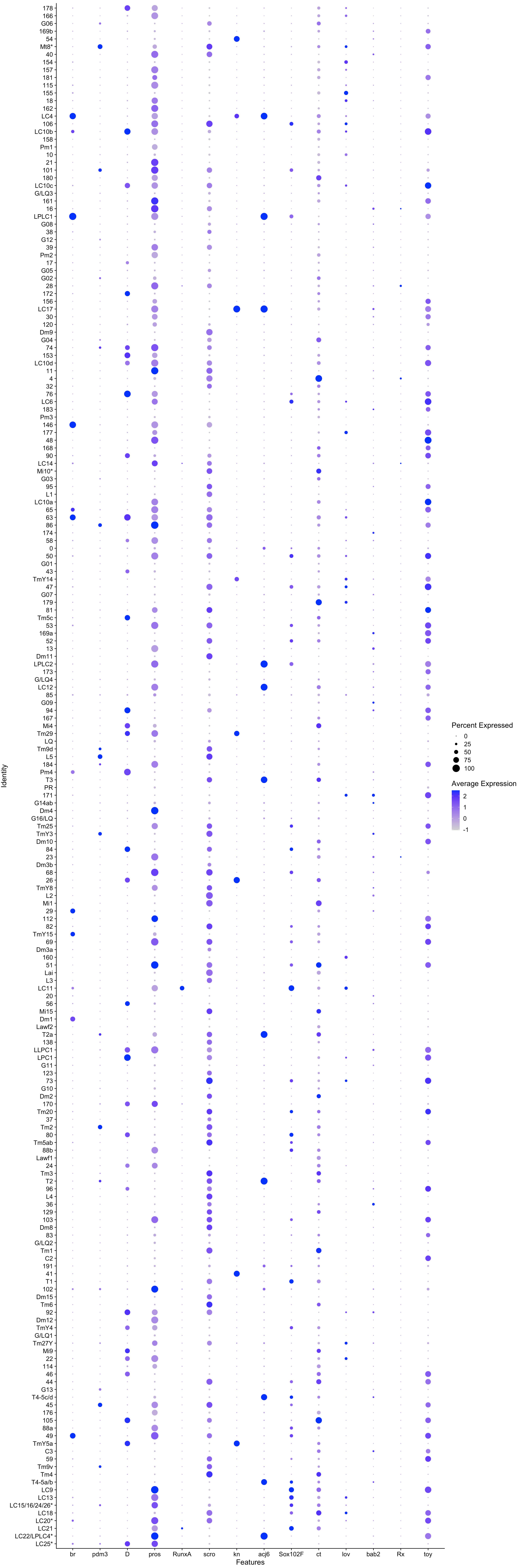
178
166
G06
169b
54
Mt8*
40
154
157
181
115
155
18
162
LC4
106
LC10b
158
Pm1
10
21
101
180
LC10c
G/LQ3
161
16
LPLC1
G08
38
G12
39
Pm2
17
G05
G02
28
172
156
LC17
30
120
Dm9
G04
74
153
LC10d
11
4
32
76
LC6
183
Pm3
146
177
48
168
90
LC14
Mi10*
G03
95
L1
LC10a
65
63
86
174
58
0
50
G01
43
TmY14
47
G07
179
81
Tm5c
53
169a
52
13
Dm11
LPLC2
173
G/LQ4
LC12
85
G09
94
167
Mi4
Tm29
LQ
Tm9d
L5
184
Pm4
T3
PR
171
G14ab
Dm4
G16/LQ
Tm25
TmY3
Dm10
84
23
Dm3b
68
26
TmY8
L2
Mi1
29
112
82
TmY15
69
Dm3a
160
51
Lai
L3
LC11
20
56
Mi15
Dm1
Lawf2
T2a
138
LLPC1
LPC1
G11
123
73
G10
Dm2
170
Tm20
37
Tm2
80
Tm5ab
88b
Lawf1
24
Tm3
T2
96
L4
36
129
103
Dm8
83
G/LQ2
Tm1
C2
191
41
T1
102
Dm15
Tm6
92
Dm12
TmY4
G/LQ1
Tm27Y
Mi9
22
114
46
44
G13
T4-5c/d
45
176
105
88a
49
TmY5a
C3
59
Tm9v
Tm4
T4-5a/b
LC9
LC13
LC15/16/24/26*
LC18
LC20*
LC21
LC22/LPLC4*
LC25*
Identity
br
pdm3
D
pros
RunxA
scro
kn
acj6
Sox102F
ct
lov
bab2
Rx
toy
Features
Percent Expressed
0
25
50
75
100
Average Expression
2
1
0
-1

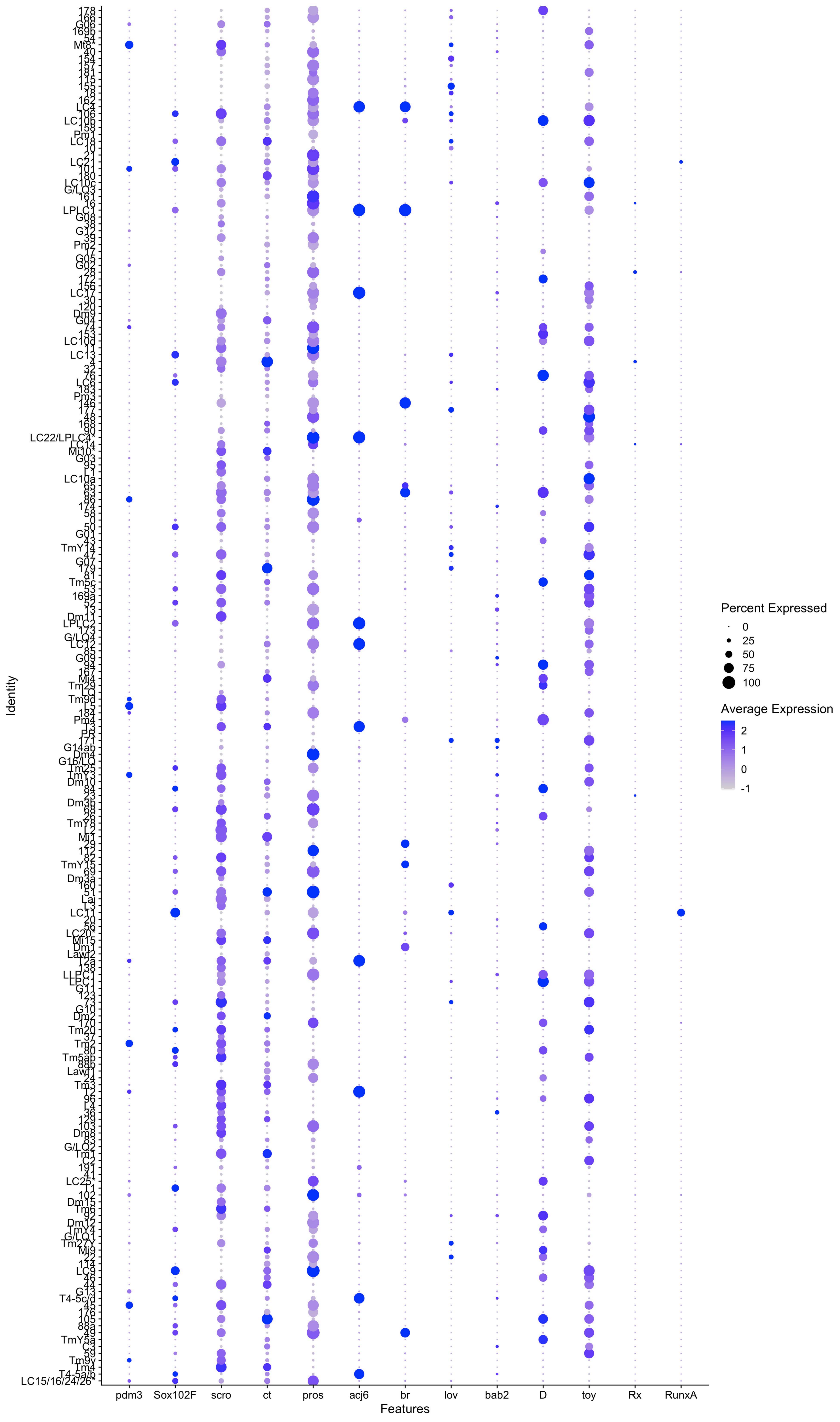
Identity
Features
pdm3
Sox102F
scro
ct
pros
acj6
br
lov
bab2
D
toy
Rx
RunxA
Percent Expressed
0
25
50
75
100
Average Expression
2
1
0
-1

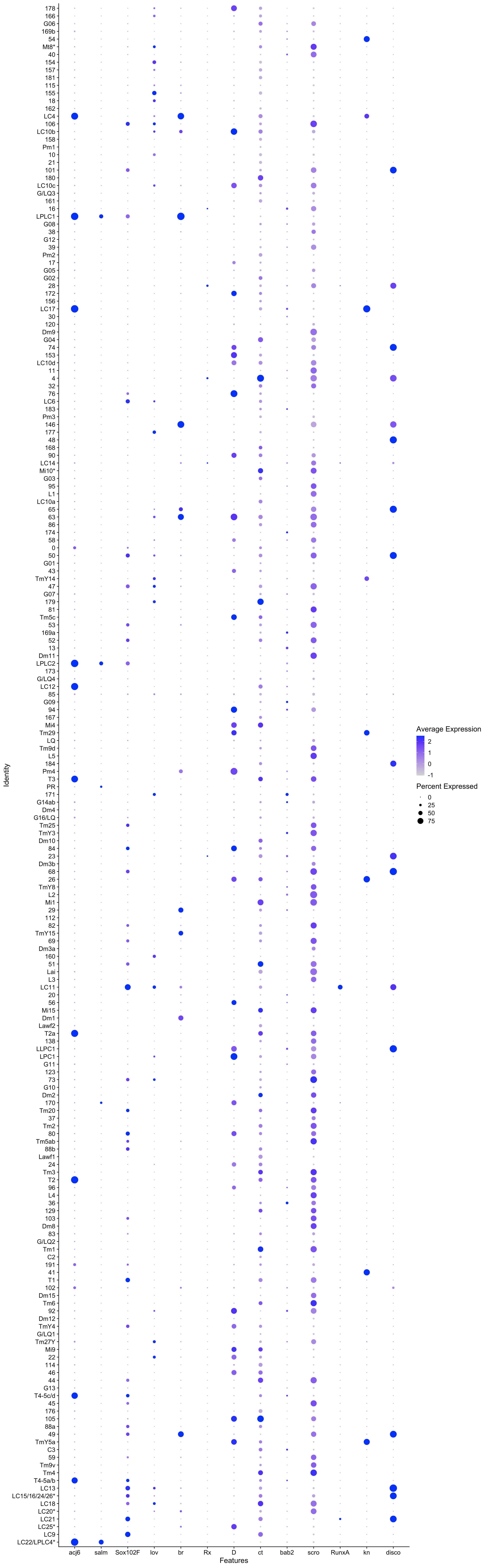
Identity
178
166
G06
169b
54
Mt8*
40
154
157
181
115
155
18
162
LC4
106
LC10b
158
Pm1
10
21
101
180
LC10c
G/LQ3
161
16
LPLC1
G08
38
G12
39
Pm2
17
G05
G02
28
172
156
LC17
30
120
Dm9
G04
74
153
LC10d
11
4
32
76
LC6
183
Pm3
146
177
48
168
90
LC14
Mi10*
G03
95
L1
LC10a
65
63
86
174
58
0
50
G01
43
TmY14
47
G07
179
81
Tm5c
53
169a
52
13
Dm11
LPLC2
173
G/LQ4
LC12
85
G09
94
167
Mi4
Tm29
LQ
Tm9d
L5
184
Pm4
T3
PR
171
G14ab
Dm4
G16/LQ
Tm25
TmY3
Dm10
84
23
Dm3b
68
26
TmY8
L2
Mi1
29
112
82
TmY15
69
Dm3a
160
51
Lai
L3
LC11
20
56
Mi15
Dm1
Lawf2
T2a
138
LLPC1
LPC1
G11
123
73
G10
Dm2
170
Tm20
37
Tm2
80
Tm5ab
88b
Lawf1
24
Tm3
T2
96
L4
36
129
103
Dm8
83
G/LQ2
Tm1
C2
191
41
T1
102
Dm15
Tm6
92
Dm12
TmY4
G/LQ1
Tm27Y
Mi9
22
114
46
44
G13
T4-5c/d
45
176
105
88a
49
TmY5a
C3
59
Tm9v
Tm4
T4-5a/b
LC13
LC15/16/24/26*
LC18
LC20*
LC21
LC25*
LC9
LC22/LPLC4*
acj6
salm
Sox102F
lov
br
Rx
D
ct
bab2
scro
RunxA
kn
disco
Features
Average Expression
2
1
0
-1
Percent Expressed
0
25
50
75
