## Supplementary Figures for "Morphological and functional convergence of visual projections neurons from diverse neurogenic origins in *Drosophila*"

### Supp Figure 1

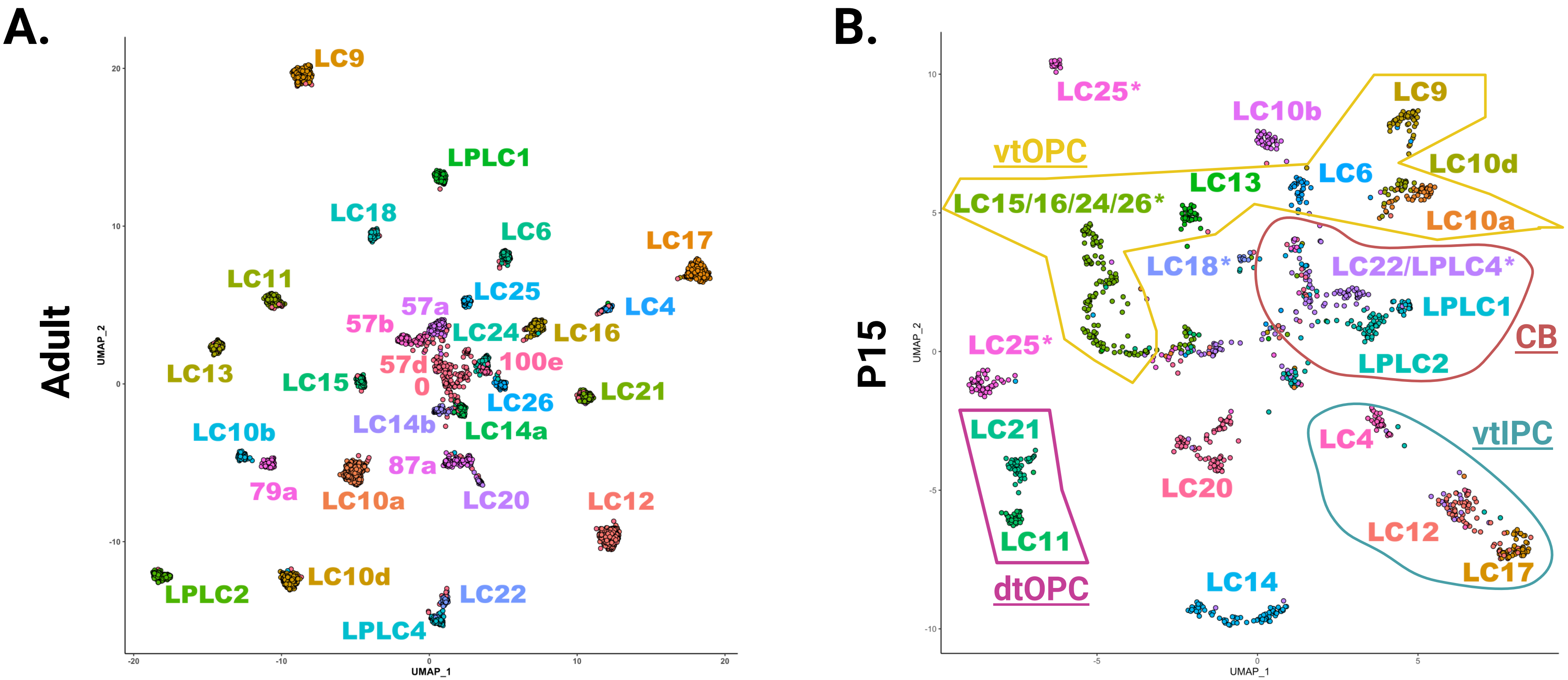

**Supp Figure 1: LCN cluster annotation across development.**

Two-dimensional UMAP plots showing all LCN and progenitor clusters in the Adult (A), and P15 (B) (Özel et al., 2021). Asterisk delineates heterogeneous clusters. Colored lines delineate LCN clusters with similar origins: yellow-OPC; red-CB; blue-vtIPC.

### Supp Figure 2

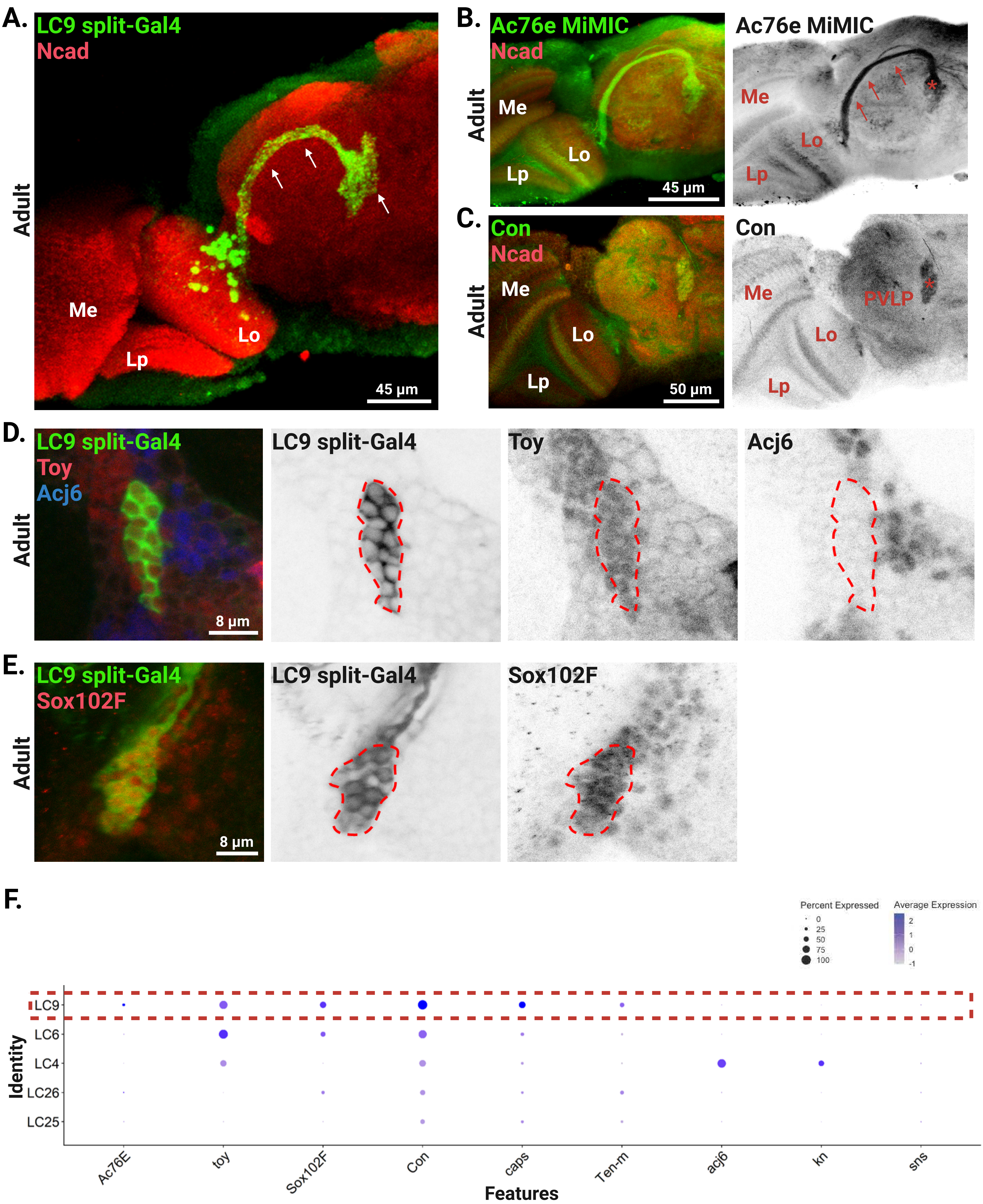

**Supp Figure 2: LC9 cluster annotation.**

A. Expression pattern of LC9 split-Gal4 (green) with Ncad (red) in the adult brain. Note the projection within the lobula (Lo), and the optic glomerulus in the central brain (arrows). Me: medulla; Lo: lobula; Lp: lobula plate.

B. Expression pattern of Ac76e MiMIC (green) with Ncad (red) in the adult brain. Note the expression of the axons (arrows) and optic glomerulus (asterisk) of LC9.

C. Expression pattern of Con (green) with Ncad (red) in the adult brain. Note the expression of Con in the optic glomerulus of LC9 (asterisk). PVLP: posterior ventrolateral protocerebrum.

D. Expression pattern of LC9 split-Gal4 (green) with Toy (red) and Acj6 (blue). Red dashed line delineates LC9 cell bodies. LC9 are Toy positive but Acj6 negative.

E. LC9 (green) express Sox102F (red). Red dashed line delineates LC9 cell bodies.

F. Dot plot showing the annotated cluster of LC9 (dashed line) showing several positive and negative markers.

### Supp Figure 3

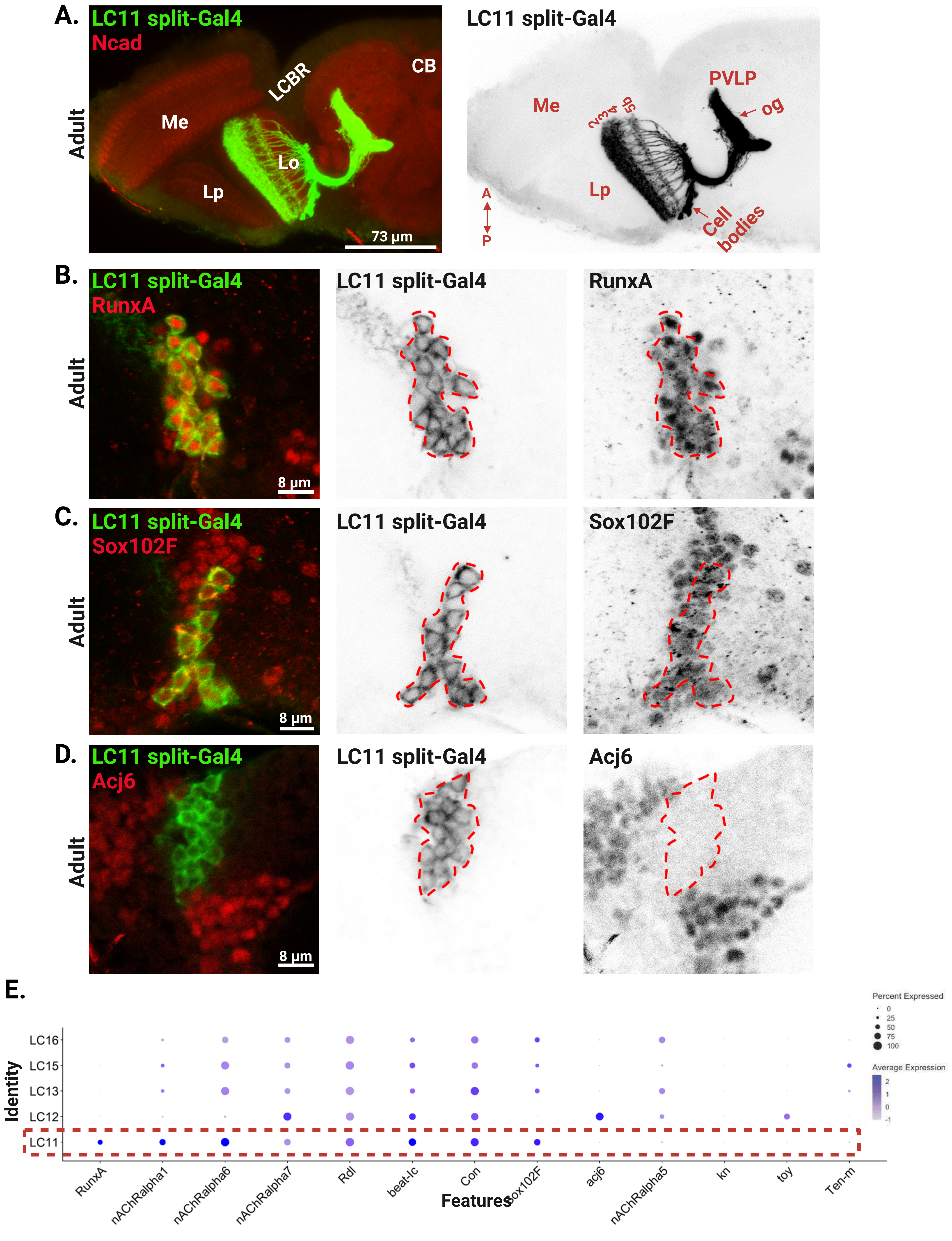

**Supp Figure 3: LC11 cluster annotation.**

A. Expression pattern of LC11 split-Gal4 (green) with Ncad (red) in the adult brain. Note the projection within the lobula (Lo) in layers 2, 3, 4 and 5b, and the optic glomerulus (og) in the central brain (arrow). Me: medulla; Lo: lobula; Lp: lobula plate; CB: central brain; LCBR: lateral cell body rind; PVLP: posterior ventrolateral protocerebrum.

B. Expression pattern of LC11 split-Gal4 (green) with RunxA (red) in the adult brain. Red dashed line delineates LC11 cell bodies. LC11 are RunxA positive.

C. LC11 (green) express Sox102F (red). Red dashed line delineates LC11 cell bodies.

D. LC11 (green) do not express Acj6 (red). Red dashed line delineates LC11 cell bodies.

E. Dot plot showing the annotated cluster of LC11 (dashed line) showing several positive and negative markers.

### Supp Figure 4

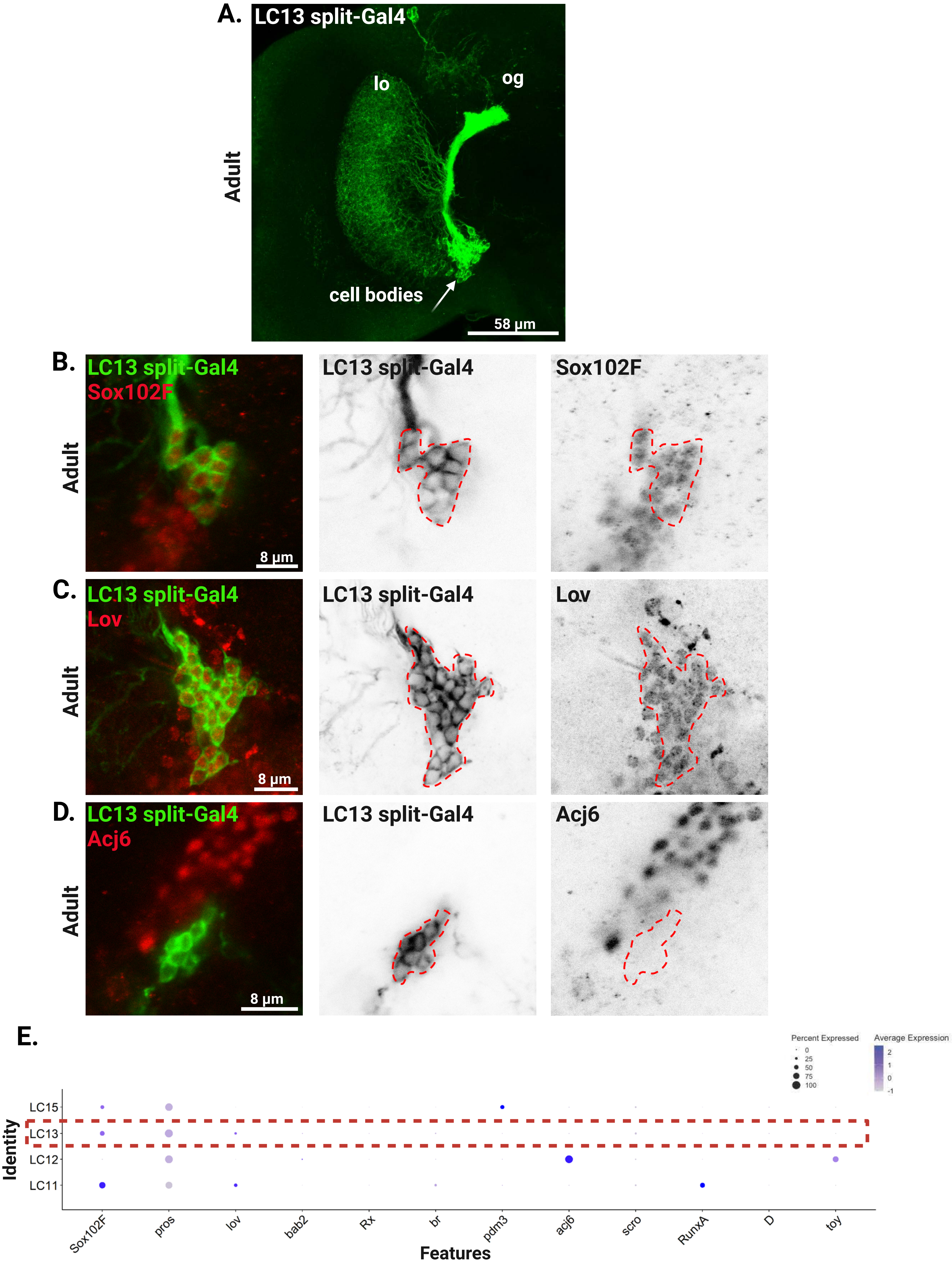

**Supp Figure 4: LC13 cluster annotation.**

A. Expression pattern of LC13 split-Gal4 (green) in the adult brain. Note the projection within the lobula (Lo), and the optic glomerulus (og) in the central brain. Arrow points to the cell bodies of LC13.
B. Expression pattern of LC13 split-Gal4 (green) with Sox102F (red) in the adult brain. Red dashed line delineates LC13 cell bodies. LC13 are Sox102F positive.
C. LC13 (green) express Lov (red). Red dashed line delineates LC13 cell bodies.
D. LC13 (green) do not express Acj6 (red). Red dashed line delineates LC13 cell bodies.
E. Dot plot showing the annotated cluster of LC13 (dashed line) showing several positive and negative markers.

#### Supp Figure 5

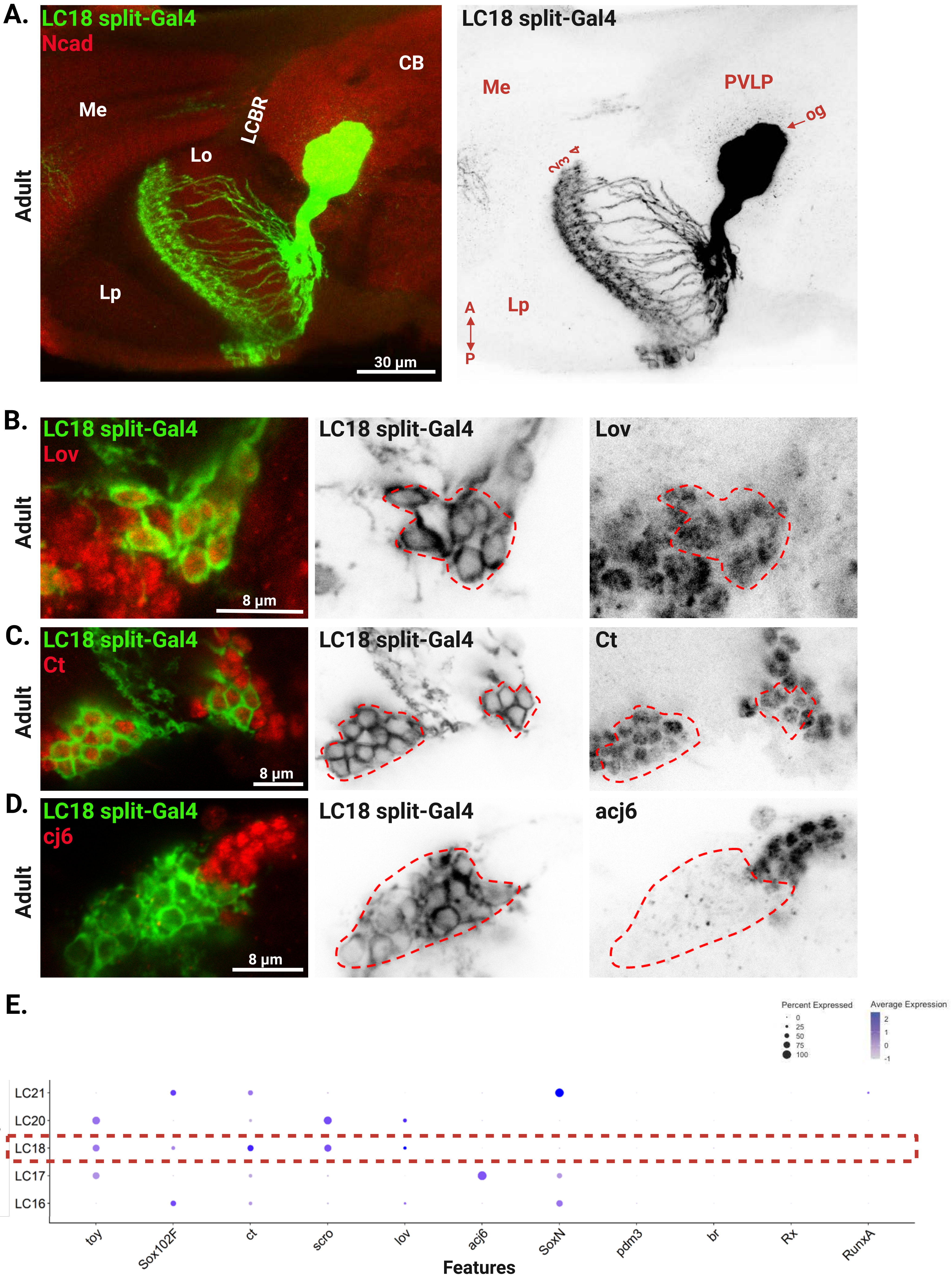

**Supp Figure 5: LC18 cluster annotation.**

A. Expression pattern of LC18 split-Gal4 (green) with Ncad (red) in the adult brain. Note the projection within the lobula (Lo) in layers 2, 3, and 4, and the optic glomerulus (og) in the central brain (arrow). Me: medulla; Lo: lobula; Lp: lobula plate; CB: central brain; LCBR: lateral cell body rind; PVLP: posterior ventrolateral protocerebrum.

B. Expression pattern of LC18 split-Gal4 (green) with Lov (red) in the adult brain. Red dashed line delineates LC18 cell bodies. LC18 are Lov positive.

C. LC18 (green) express Ct (red). Red dashed line delineates LC18 cell bodies.

D. LC18 (green) do not express Acj6 (red). Red dashed line delineates LC18 cell bodies.

E. Dot plot showing the annotated cluster of LC18 (dashed line) showing several positive and negative markers.

### Supp Figure 6

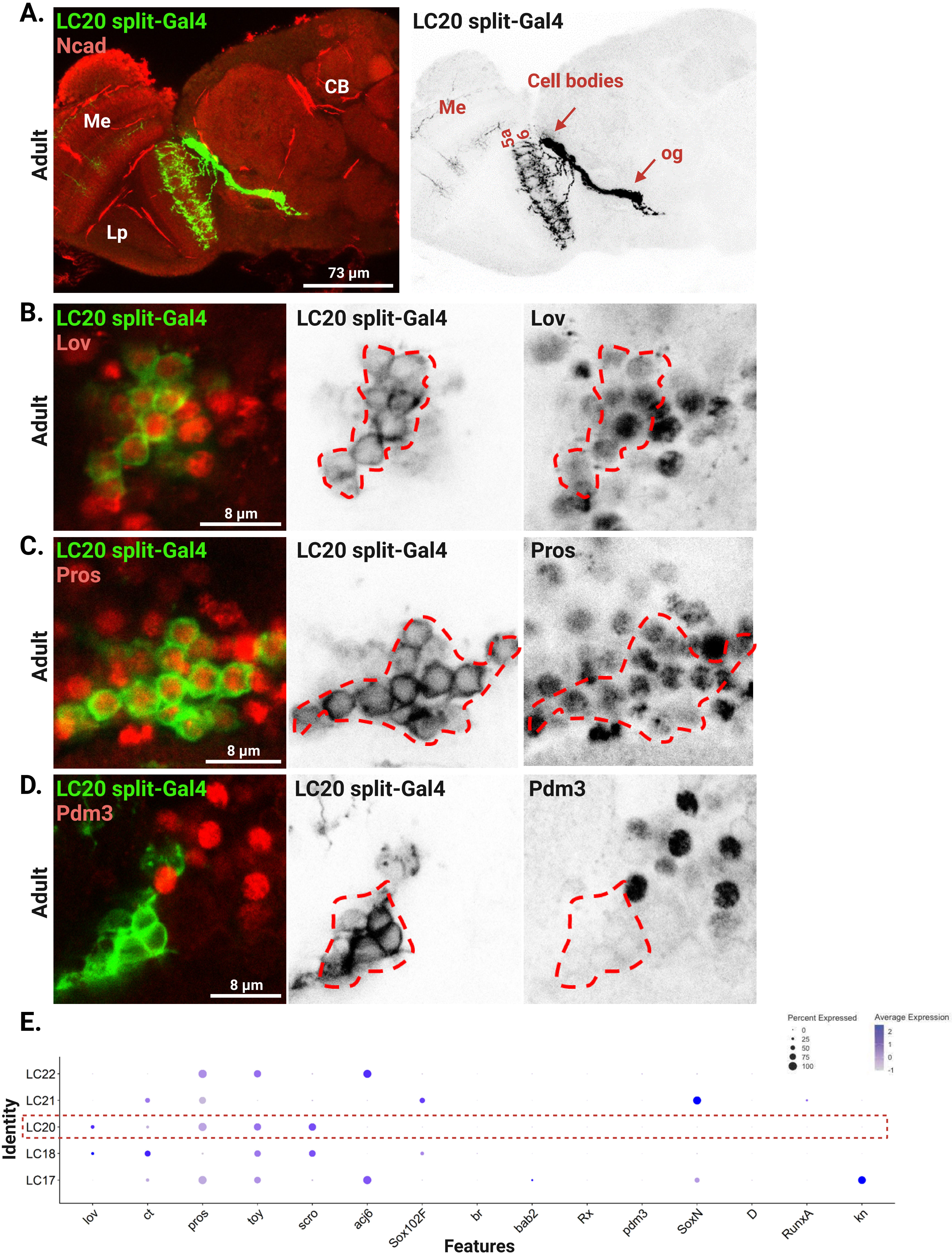

**Supp Figure 6: LC20 cluster annotation.**

A. Expression pattern of LC20 split-Gal4 (green) with Ncad (red) in the adult brain. Note the projection within the lobula in layers 5a and 6, and the optic glomerulus (og) in the central brain (arrow). Me: medulla; Lp: lobula plate; CB: central brain.

B. Expression pattern of LC20 split-Gal4 (green) with Lov (red) in the adult brain. Red dashed line delineates LC20 cell bodies. LC20 are Lov positive.

C. LC20 (green) express Pros (red). Red dashed line delineates LC20 cell bodies.

D. LC20 (green) do not express Pdm3 (red). Red dashed line delineates LC20 cell bodies.

E. Dot plot showing the annotated cluster of LC20 (dashed line) showing several positive and negative markers.

### Supp Figure 7

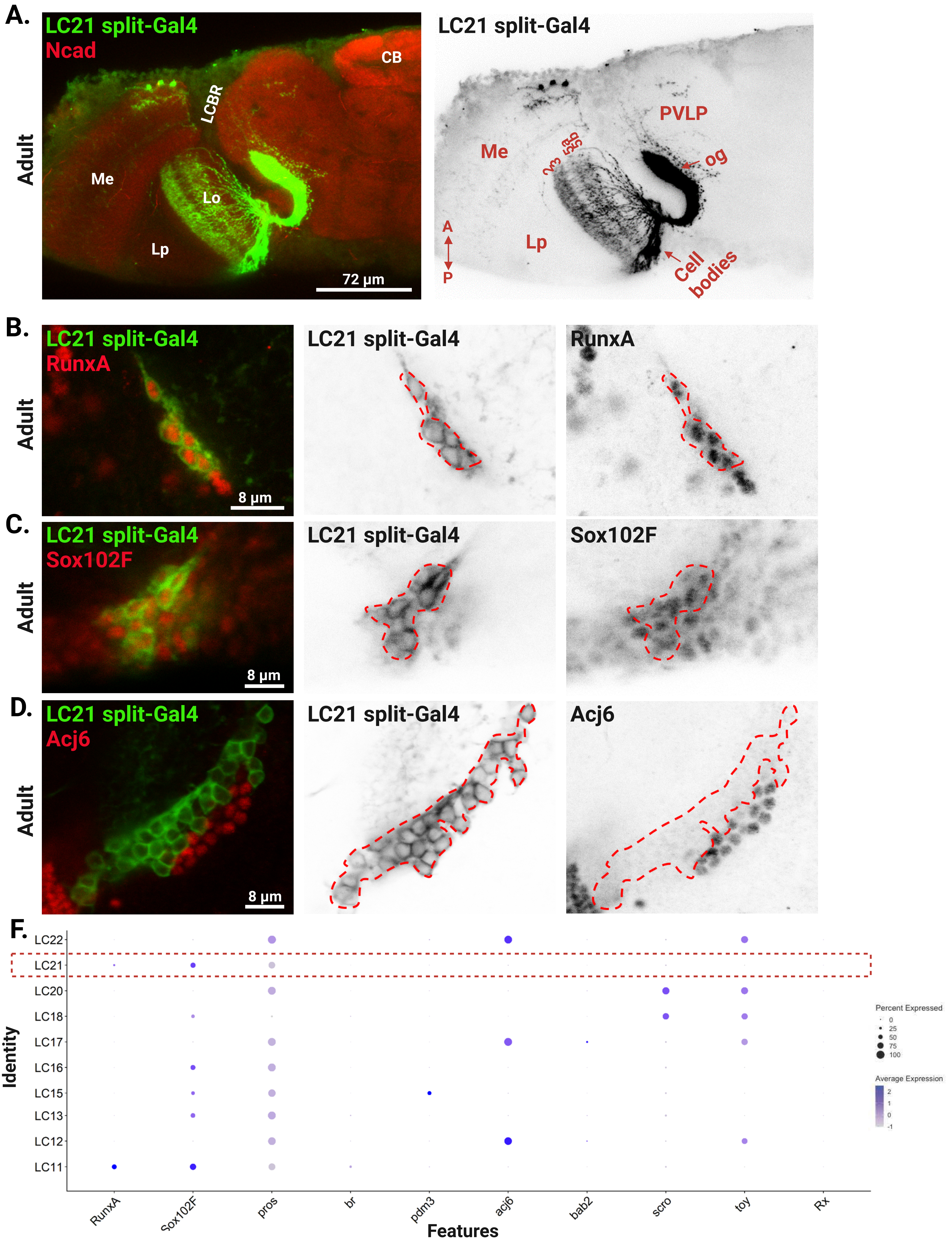

**Supp Figure 7: LC21 cluster annotation.**

A. Expression pattern of LC21 split-Gal4 (green) with Ncad (red) in the adult brain. Note the projection within the lobula (Lo) in layers 2,3,5a and 5b, and the optic glomerulus (og) in the central brain (arrow). Me: medulla; Lp: lobula plate; CB: central brain; LCBR: lateral cell body rind; PVLP: posterior ventrolateral protocerebrum.

B. Expression pattern of LC21 split-Gal4 (green) with RunxA (red) in the adult brain. Red dashed line delineates LC21 cell bodies, which are RunxA positive.

C. LC21 (green) express Sox102F (red). Red dashed line delineates LC21 cell bodies.

D. LC21 (green) do not express Acj6 (red). Red dashed line delineates LC21 cell bodies.

E. Dot plot showing the annotated cluster of LC21 (dashed line) showing several positive and negative markers.

**Supp Figure 8**

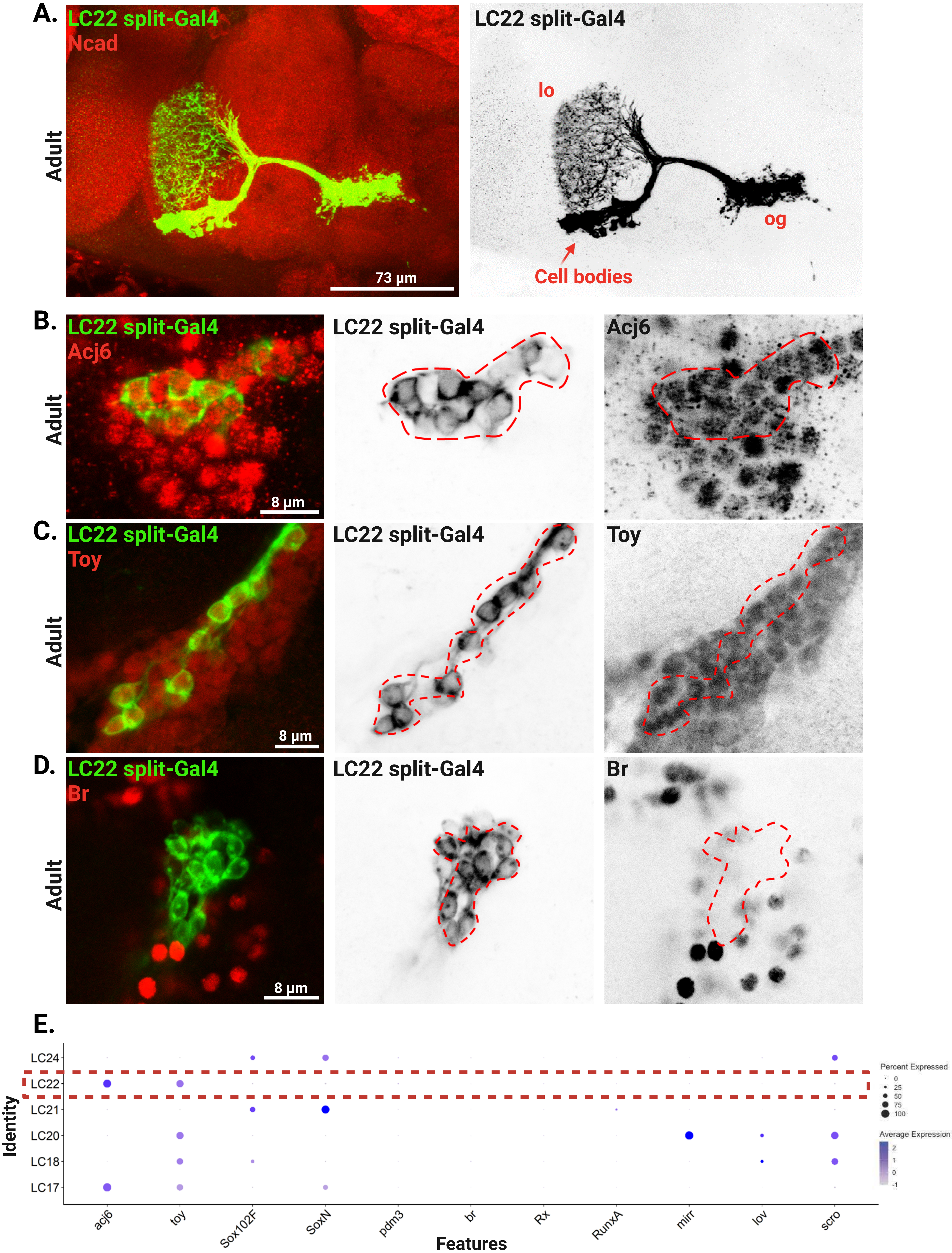

**Supp Figure 8: LC22 cluster annotation.**

A. Expression pattern of LC22 split-Gal4 (green) with Ncad (red) in the adult brain. Note the projection within the lobula (Lo), and the optic glomerulus (og) in the central brain. Arrow denotes the cell bodies of LC22.

B. Expression pattern of LC22 split-Gal4 (green) with Acj6 (red) in the adult brain. Red dashed line delineates LC22 cell bodies, which are Acj6 positive.

C. LC22 (green) express Toy (red). Red dashed line delineates LC22 cell bodies.

D. LC22 (green) do not express Br (red). Red dashed line delineates LC22 cell bodies.

E. Dot plot showing the annotated cluster of LC22 (dashed line) showing several positive and negative markers.

### Supp Figure 9

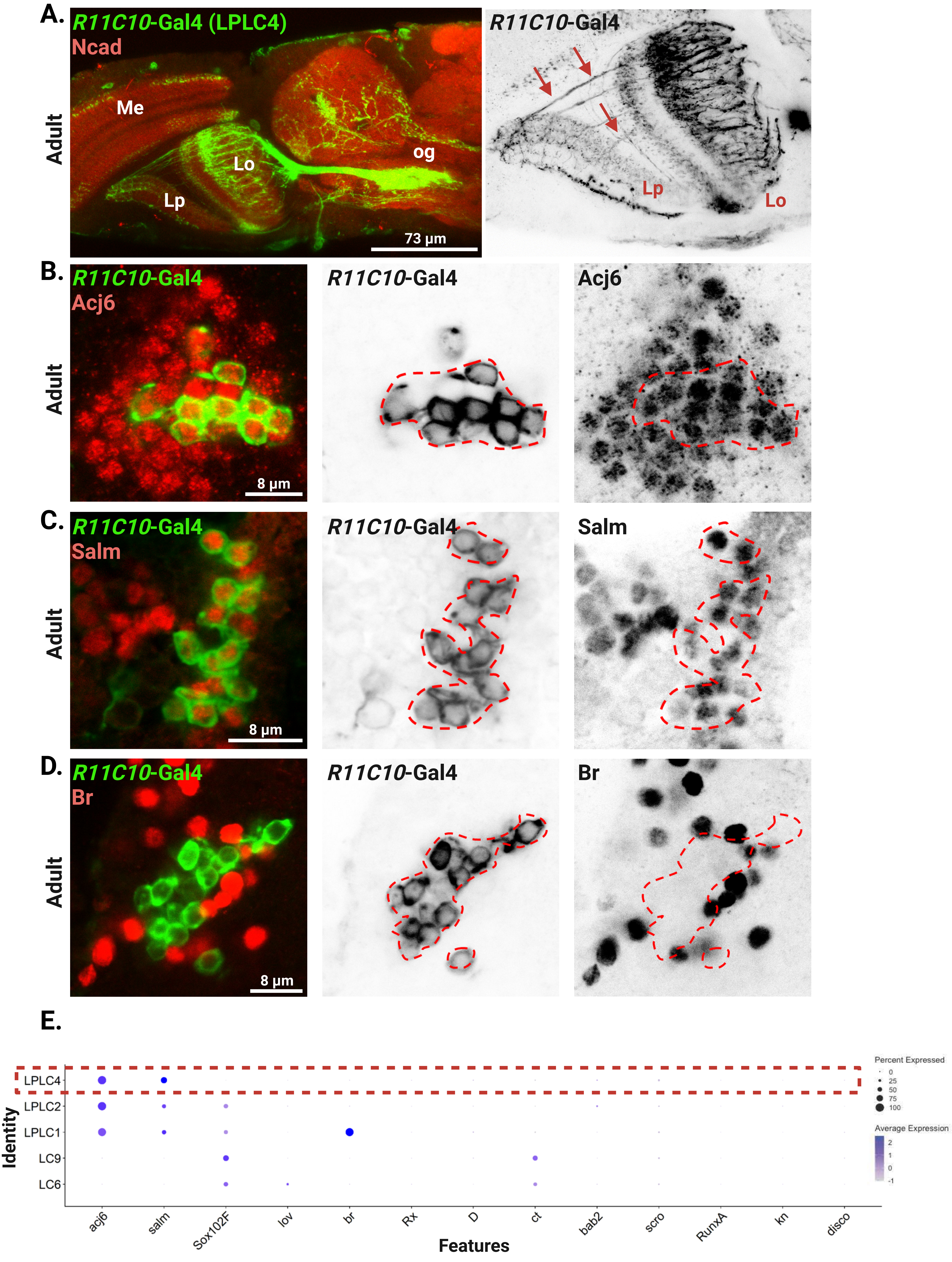

**Supp Figure 9: LPLC4 cluster annotation.**

A. Expression pattern of *R11C10*-Gal4 (green) with Ncad (red) in the adult brain, showing the presence of LPLC4 neurons. Right panel: Note the projection from the lobula (Lo) extending to the lobula plate (Lp) (arrows); and the optic glomerulus (og) in the central brain. Me: medulla.

B. Expression pattern of *R11C10*-Gal4 (green; LPLC4) with Acj6 (red) in the adult brain. Red dashed line delineates LPLC4 cell bodies, which are Acj6 positive.

C. LPLC4 (green) express Salm (red). Red dashed line delineates LPLC4 cell bodies.

D. LPLC4 (green) do not express Br (red). Red dashed line delineates LPLC4 cell bodies.

E. Dot plot showing the annotated cluster of LPLC4 (dashed line) showing several positive and negative markers.

#### Supp Figure 10

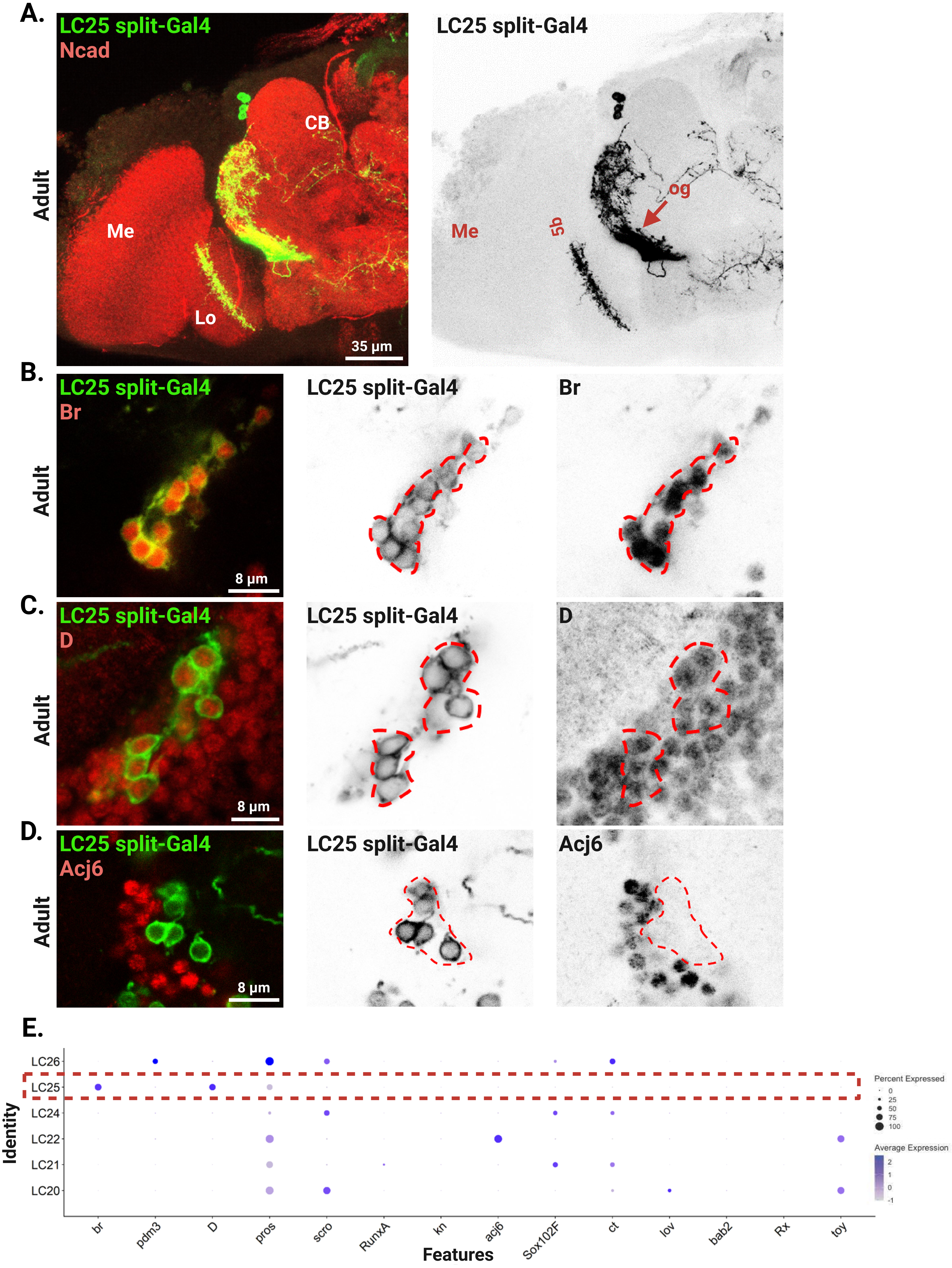

**Supp Figure 10: LC25 cluster annotation.**

A. Expression pattern of LC25 split-Gal4 (green) with Ncad (red) in the adult brain. Note the projection within the lobula (Lo) layer 5b, and the optic glomerulus (og) in the central brain (CB) (arrow). Me: medulla.

B. Expression pattern of LC25 split-Gal4 (green) with Br (red) in the adult brain. Red dashed line delineates LC25 cell bodies, which are Br positive.

C. LC25 (green) express D (red). Red dashed line delineates LC25 cell bodies.

D. LC25 (green) do not express Acj6 (red). Red dashed line delineates LC25 cell bodies.

E. Dot plot showing the annotated cluster of LC25 (dashed line) showing several positive and negative markers

#### Supp Figure 11

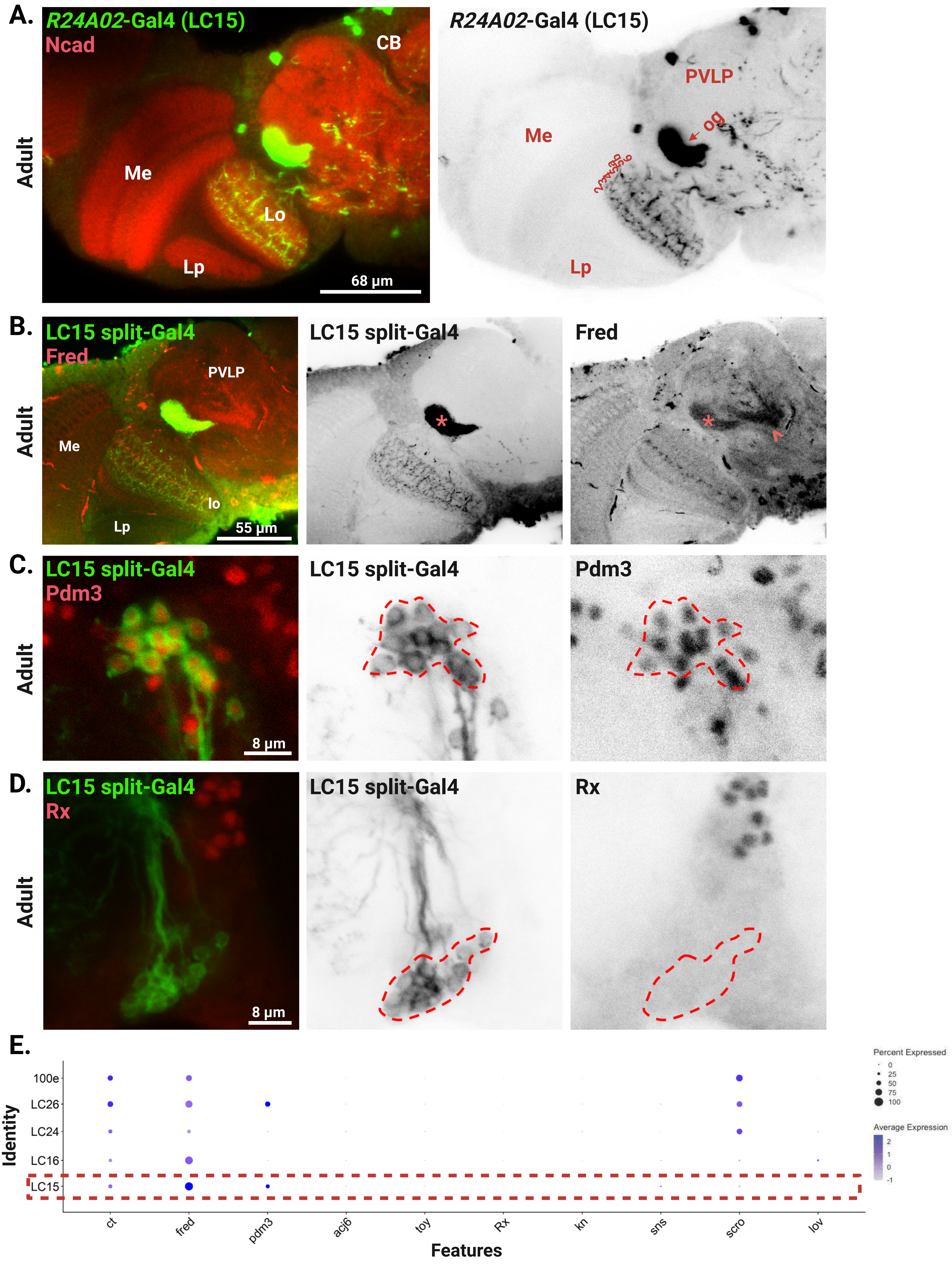

**Supp Figure 11: LC15 cluster annotation.**

A. Expression pattern of *R42A02*-Gal4 (green) with Ncad (red) in the adult brain, showing the presence of LC15. Note the projection throughout different lobula (Lo) layers, and the optic glomerulus (og) in the central brain (CB) (arrow). Me: medulla; Lp: lobula plate; PVLP: posterior ventrolateral protocerebrum.

B. Expression pattern of LC15 split-Gal4 (green) with Fred (red) in the adult brain. Asterisk delineates LC15 optic glomerulus. Note the neighboring optic glomerulus (arrowhead) is also Fred positive.

C. LC15 (green) express Pdm3 (red). Red dashed line delineates LC15 cell bodies.

D. LC15 (green) do not express Rx (red). Red dashed line delineates LC15 cell bodies.

E. Dot plot showing the annotated cluster of LC15 (dashed line) showing several positive and negative markers.

#### Supp Figure 12

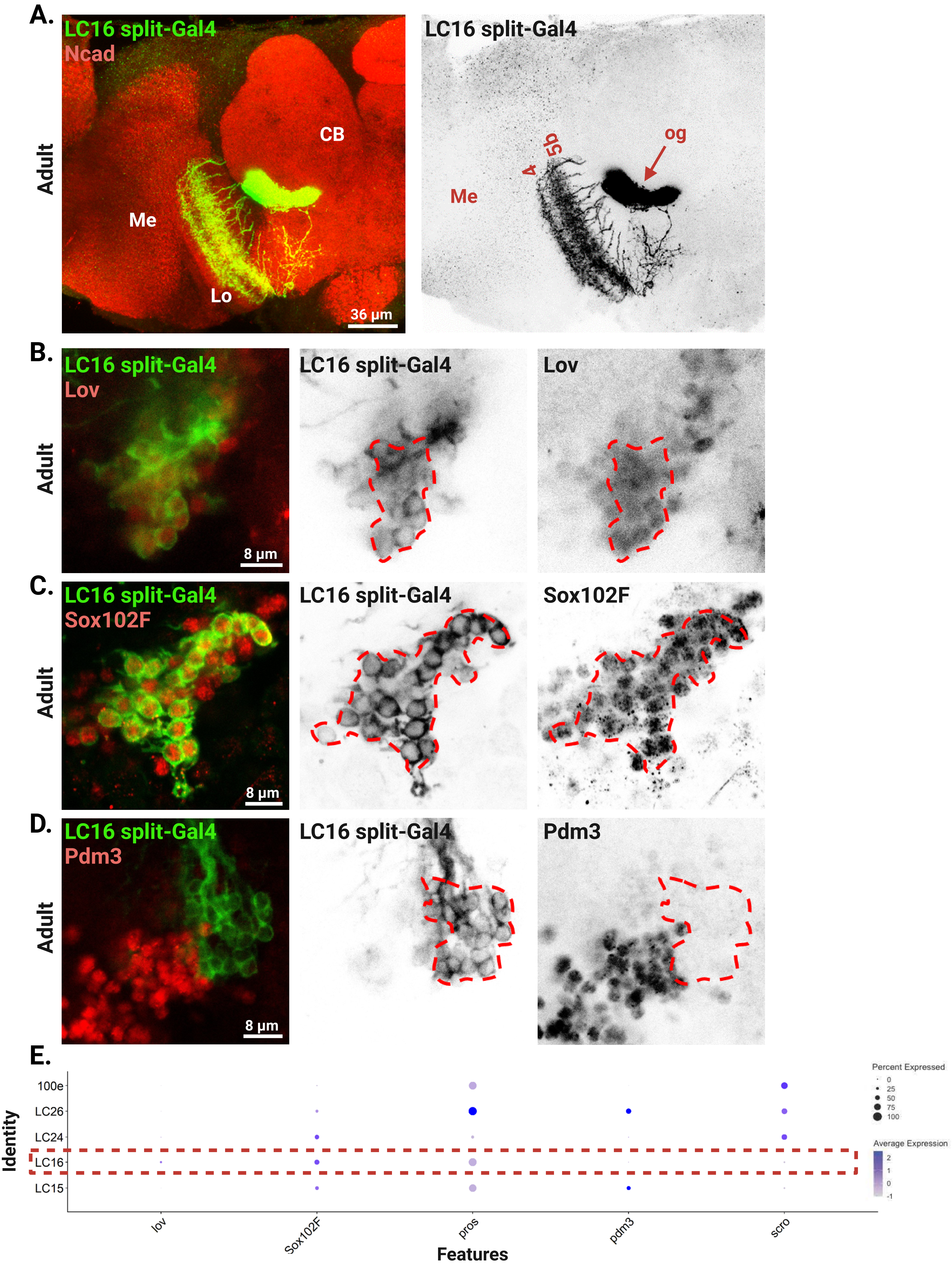

**Supp Figure 12: LC16 cluster annotation.**

A. Expression pattern of LC16 split-Gal4 (green) with Ncad (red) in the adult brain. Note the projection in lobula (Lo) layers 4 and 5b, and the optic glomerulus (og) in the central brain (CB) (arrow). Me: medulla.

B. Expression pattern of LC16 split-Gal4 (green) with Lov (red) in the adult brain. Red dashed line delineates LC25 cell bodies, which are Lov positive.

C. LC16 (green) express Sox102F (red). Red dashed line delineates LC16 cell bodies.

D. LC16 (green) do not express Pdm3 (red). Red dashed line delineates LC16 cell bodies.

E. Dot plot showing the annotated cluster of LC16 (dashed line) showing several positive and negative markers.

### Supp Figure 13

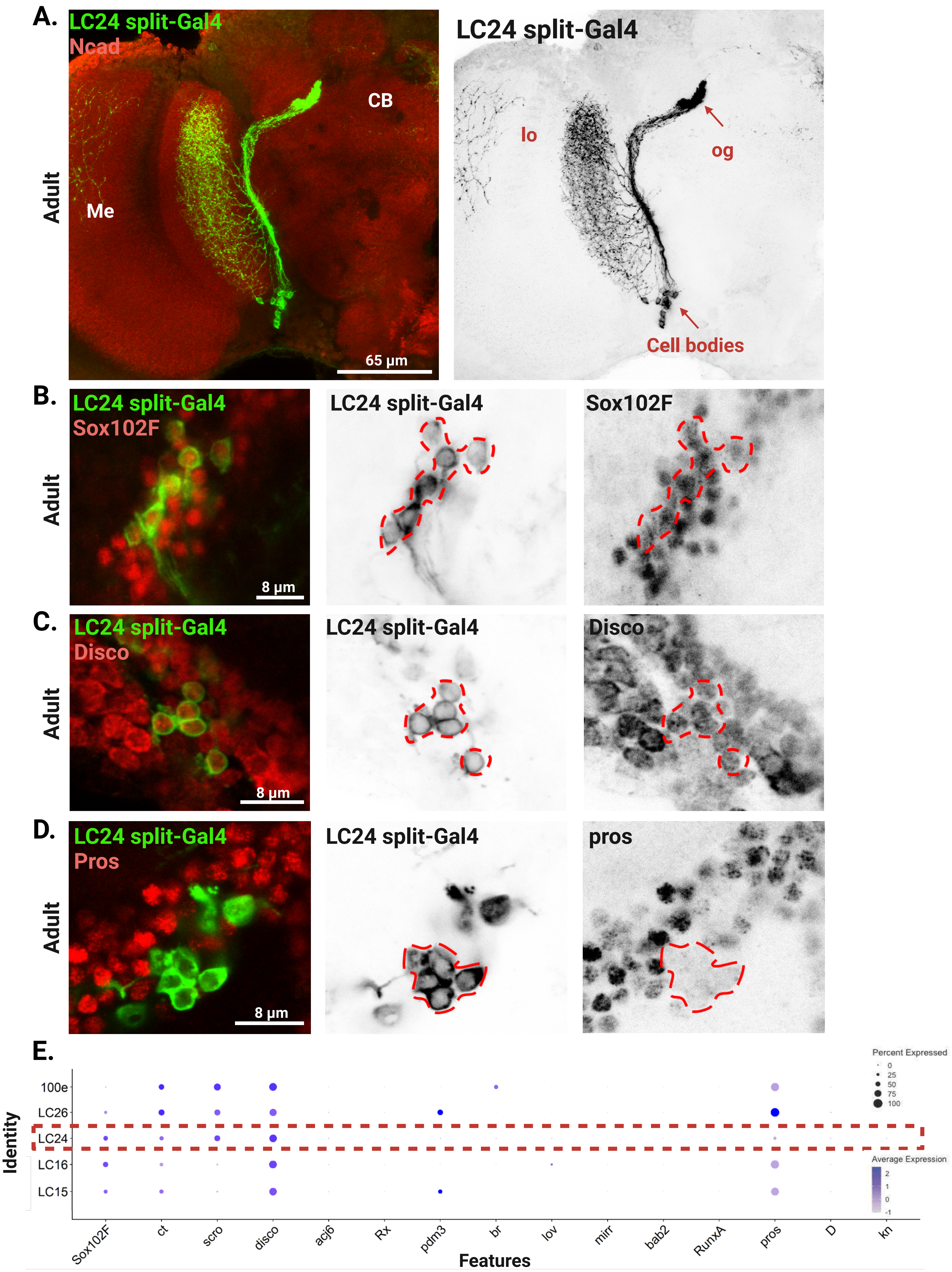

**Supp Figure 13: LC24 cluster annotation.**

A. Expression pattern of LC24 split-Gal4 (green) with Ncad (red) in the adult brain. Note the projection in the lobula (Lo) and the optic glomerulus (og) in the central brain (CB). Arrow points to LC24 cell bodies. Me: medulla.

B. Expression pattern of LC24 split-Gal4 (green) with Sox102F (red) in the adult brain. Red dashed line delineates LC24 cell bodies, which are Sox102F positive.

C. LC24 (green) express Disco (red). Red dashed line delineates LC24 cell bodies.

D. LC24 (green) do not express Pros (red). Red dashed line delineates LC24 cell bodies.

E. Dot plot showing the annotated cluster of LC24 (dashed line) showing several positive and negative markers.

### Supp Figure 14

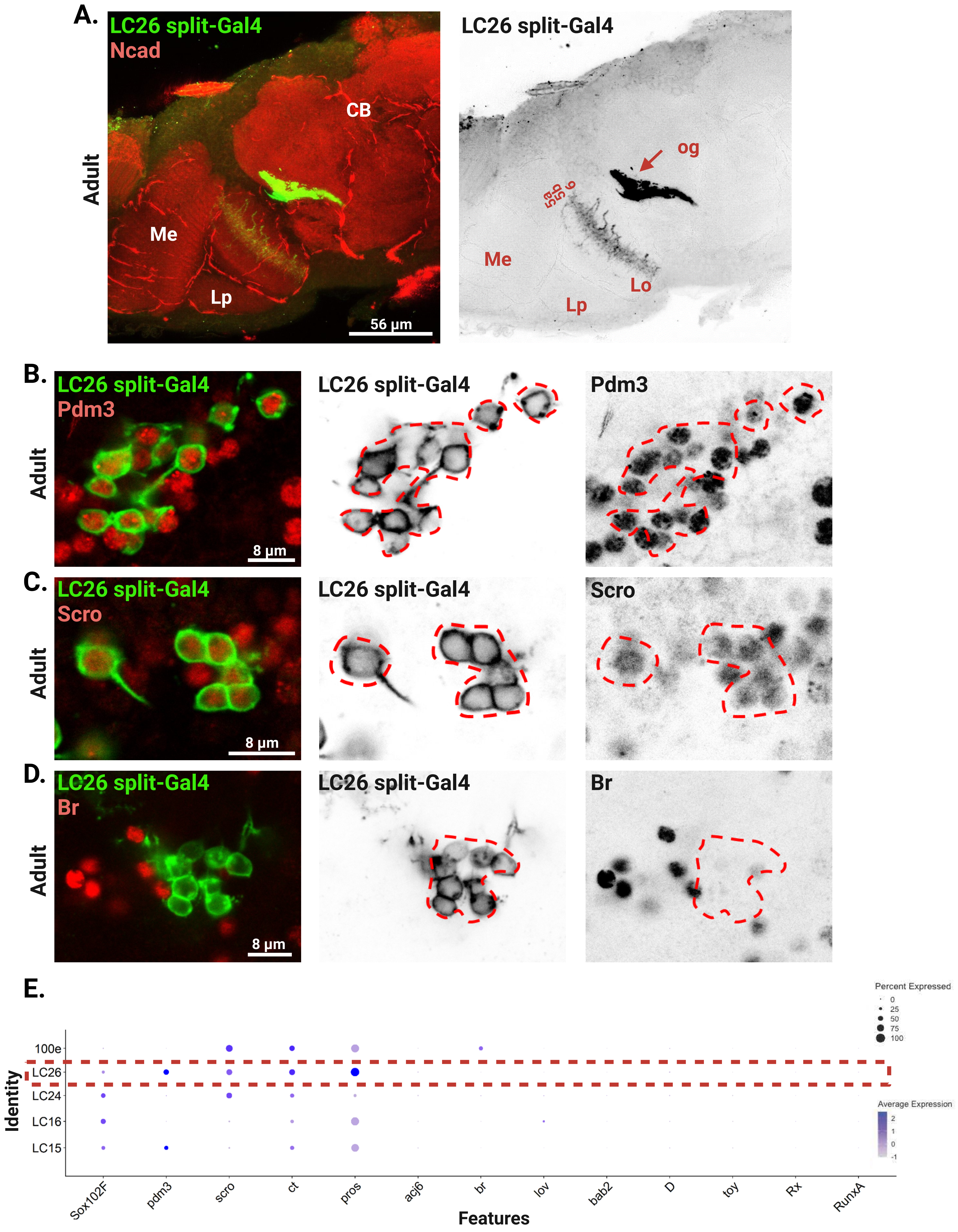

**Supp Figure 14: LC26 cluster annotation.**

A. Expression pattern of LC26 split-Gal4 (green) with Ncad (red) in the adult brain. Note the projection in the lobula (Lo) layers 5a, 5b and 6, and the optic glomerulus (og) in the central brain (CB) arrow. Me: medulla; Lp: lobula plate.

B. Expression pattern of LC26 split-Gal4 (green) with Pdm3 (red) in the adult brain. Red dashed line delineates LC26 cell bodies, which are Pdm3 positive.

C. LC26 (green) express Scro (red). Red dashed line delineates LC26 cell bodies.

D. LC26 (green) do not express Br (red). Red dashed line delineates LC26 cell bodies.

E. Dot plot showing the annotated cluster of LC26 (dashed line) showing several positive and negative markers

### Supp Figure 15

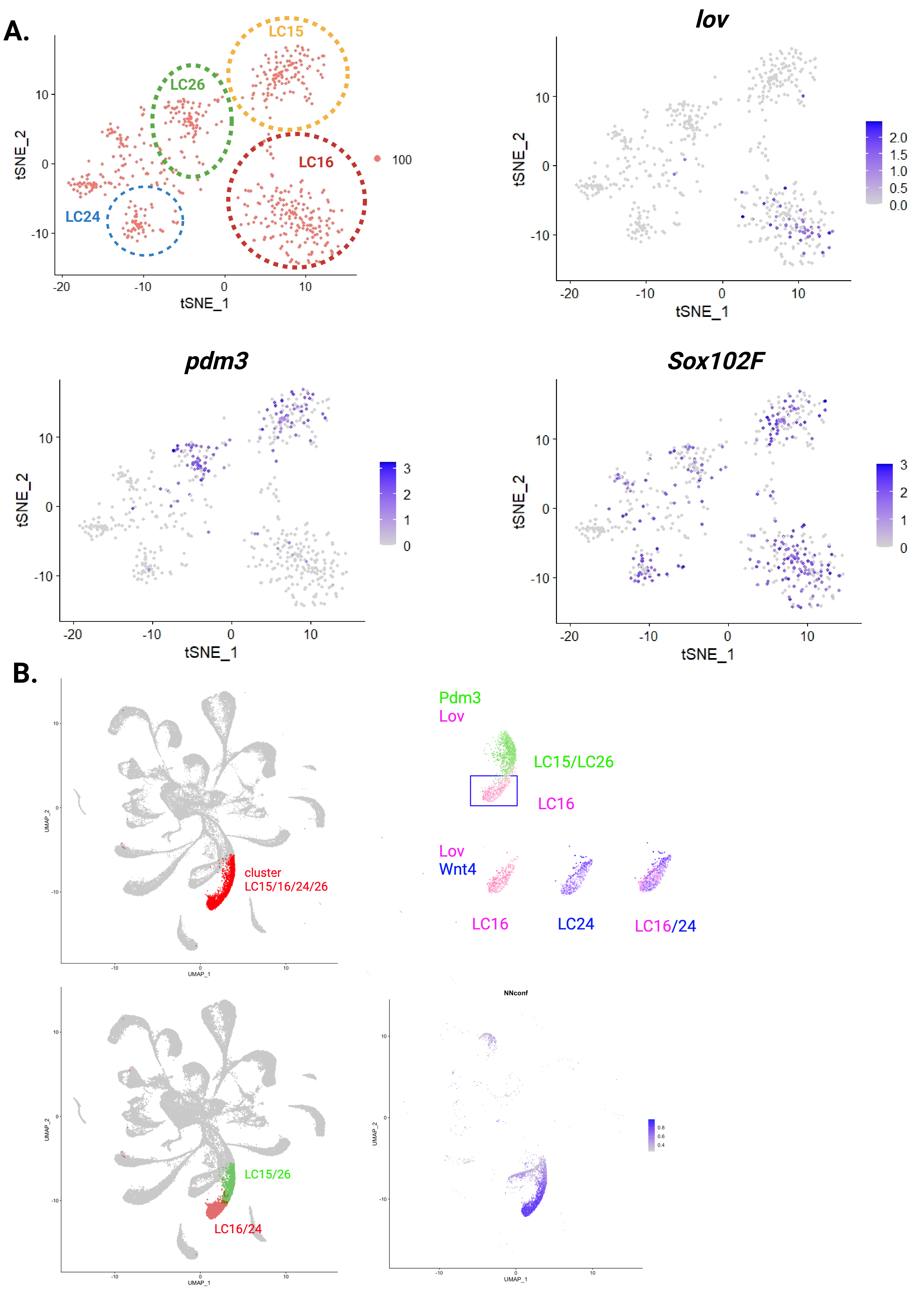

**Supp Figure 15: Reclustering cluster 100: LC15, LC16. LC24 and LC26.**

A. Left top panel: tSNE plots showing 5 sub-clusters contained within cluster 100. LC15, yellow dashed line; LC16: red dashed line; LC24: blue dashed line and LC26: green dashed line. LC16 is the only sub-cluster expressing Lov (right top panel). LC15 and LC26 clusters express Pdm3 (left bottom panel); while all clusters express Sox102F (right bottom panel).

B. UMAP plots of single cells coming from developing Wg region. We were able to further separate the cluster corresponding to LC15/LC16/LC24/LC26 (upper panel, marked in red) into two distinct subclusters: LC15/LC26 (lower panel, green; Pdm3-positive cells) and LC16/LC24 (lower panel, red color; Wnt4-positive and Lov-positive cells). Further distinction within this subcluster reveals that LC16 cells are marked by Lov expression (pink), whereas LC24 cells are defined by their unique expression profile: they express Wnt4, but lack the expression of Pdm3 and Lov (purple).

#### Supp Figure 16

A.

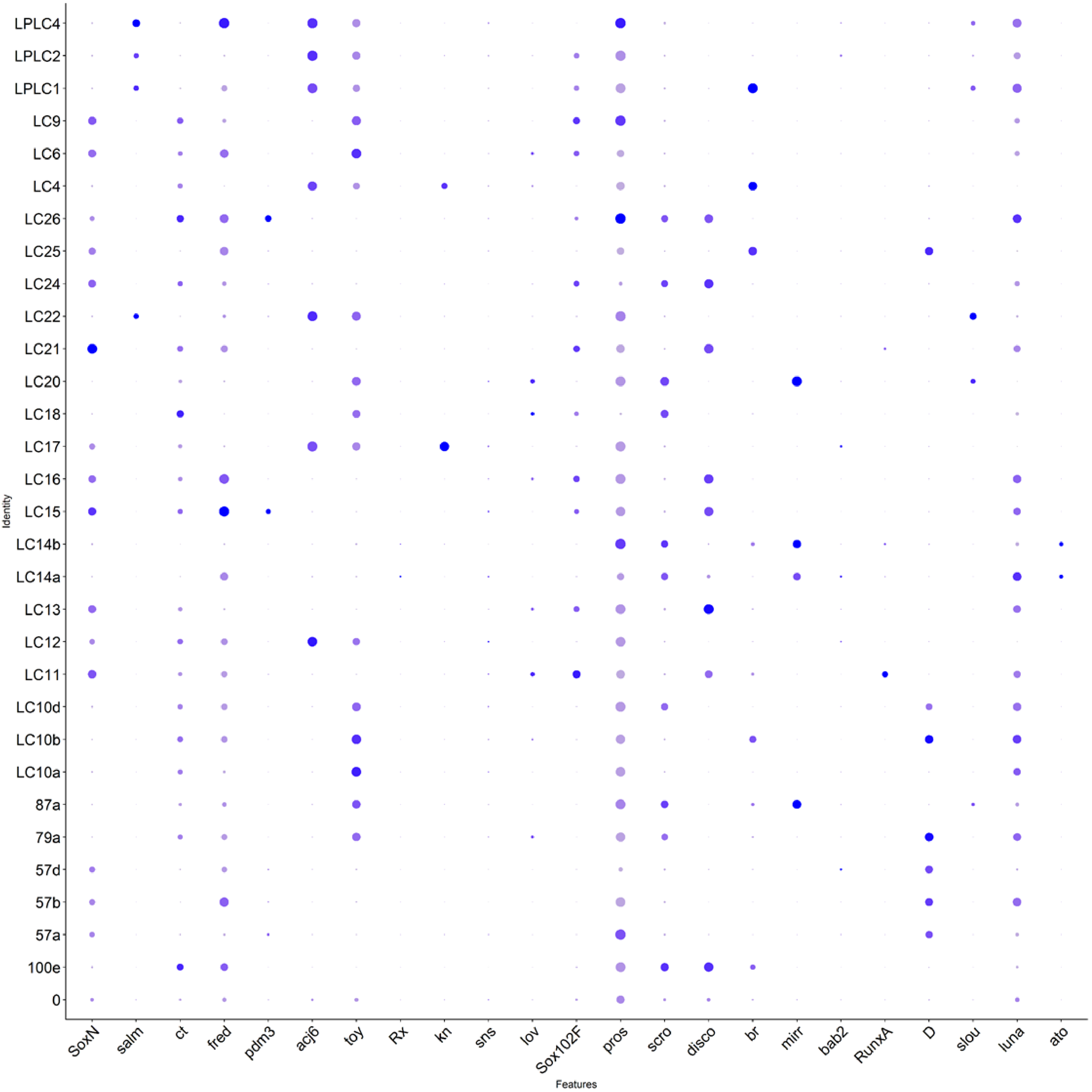


B.

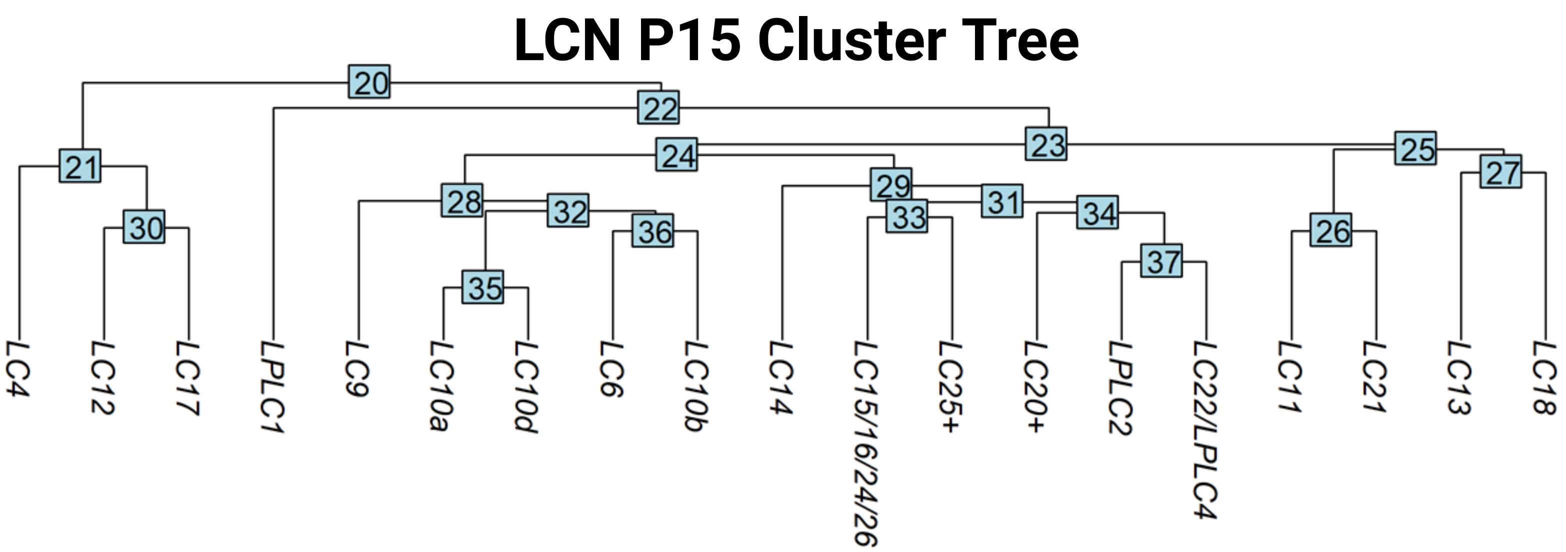

**Supp Figure 16: LCN markers.**

A. Dot plot showing different markers in LCN adult clusters. Note that each cluster has a unique combination of genes.

B. Cluster tree analysis of LCNs at P15. Note that LCNs that originate from a similar origin cluster together. For example, LC4, LC12 and LC17 have a similar origin from the vtIPC; while LC6, LC9 and LC10s seem to have closely related origins as well.

### Supp Figure 17

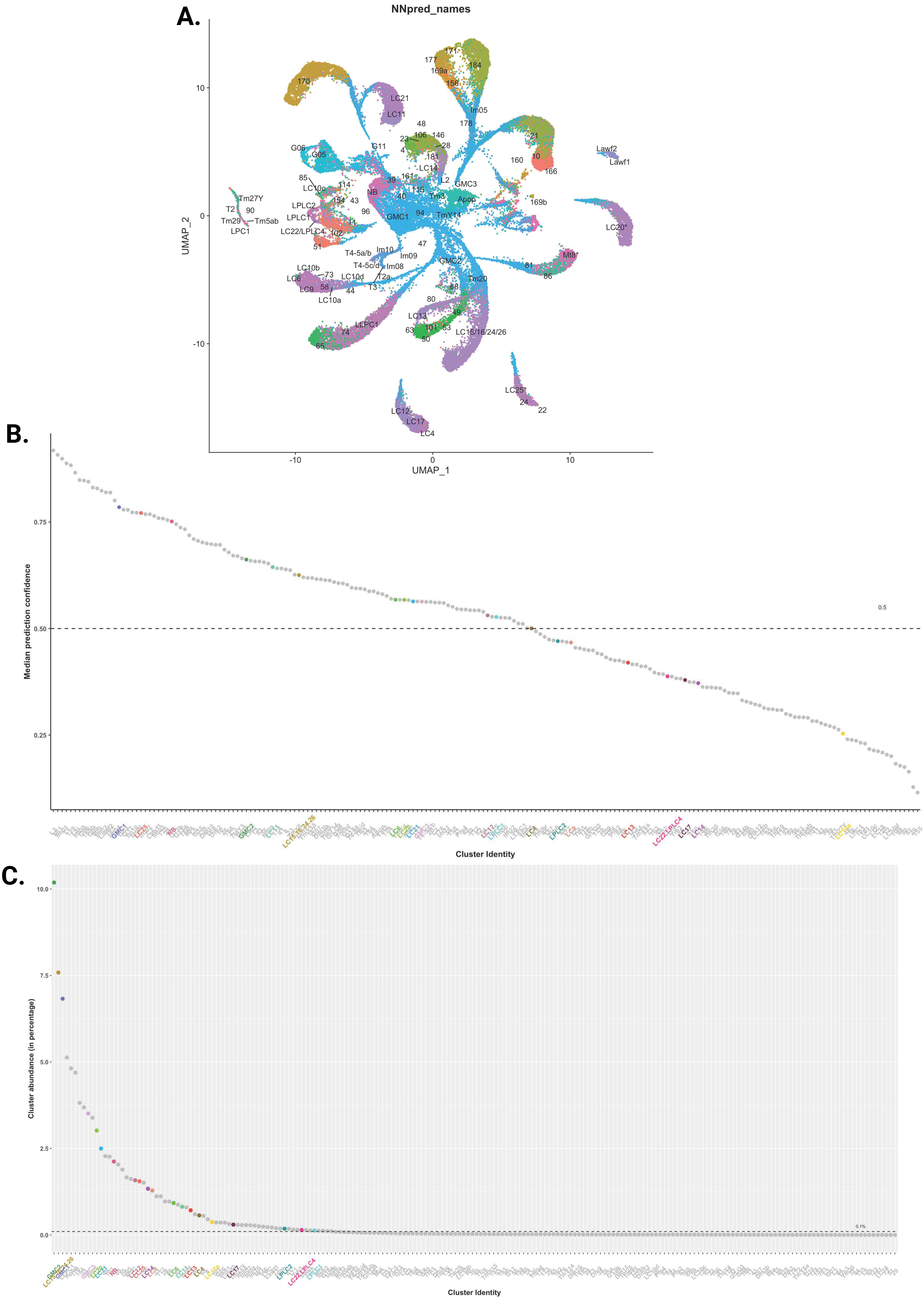

**Supp Figure 17 Neural Network (NN) annotation of the L3 *wg*-Gal4 scRNAseq dataset.**
A. Two-dimensional UMAP plot for all NN predicted clusters.
B. Median prediction confidence of the NN classified dataset highlighting LCN clusters from Supplementary Table 1.
C. Plot showing the cluster abundance (in percentage) of LCN clusters in the dataset from Supplementary Table 1.

### Supp Figure 18

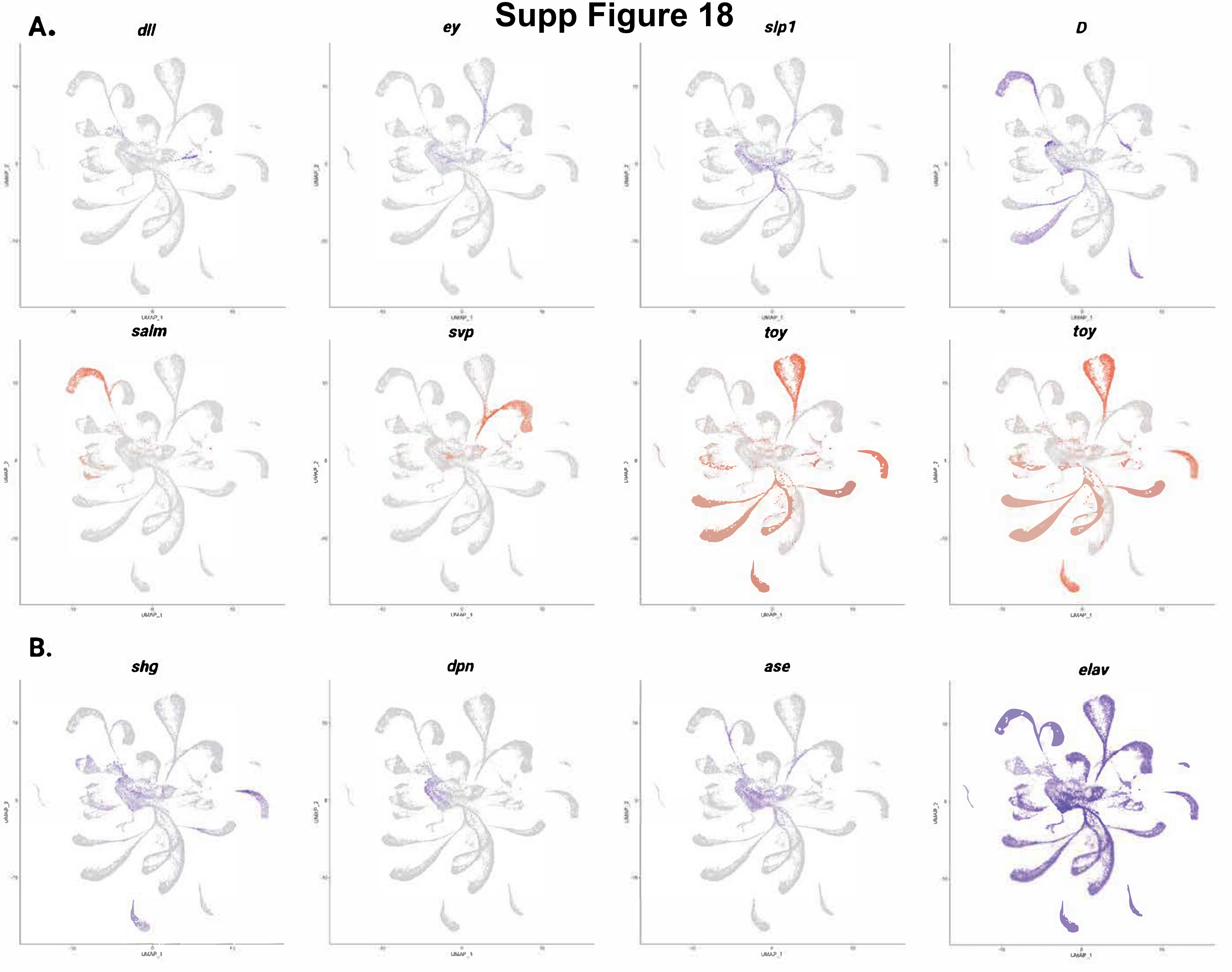

**<u>Supp Fig 18: tTFs and markers expression pattern in the tOPC neuroblasts.</u>**

A. tOPC neuroblasts sequentially express *dll*, *ey*, *slp* and *D* (blue). Each of these time windows produces distinct neuronal subtypes, marked by the expression of specific genes (red; *salm, svp, toy and toy*).

B. UMAP plots showing the expression of different cell type markers: *shg* is expressed in the neuroepithelium and young neuroblasts; *dpn* is expressed in the neuroepithelium and neuroblasts; *ase* is expressed in neuroblasts and GMCs, while *elav* is primarily expressed in neurons, although its expression can already be observed in GMCs.

### Supp Figure 19

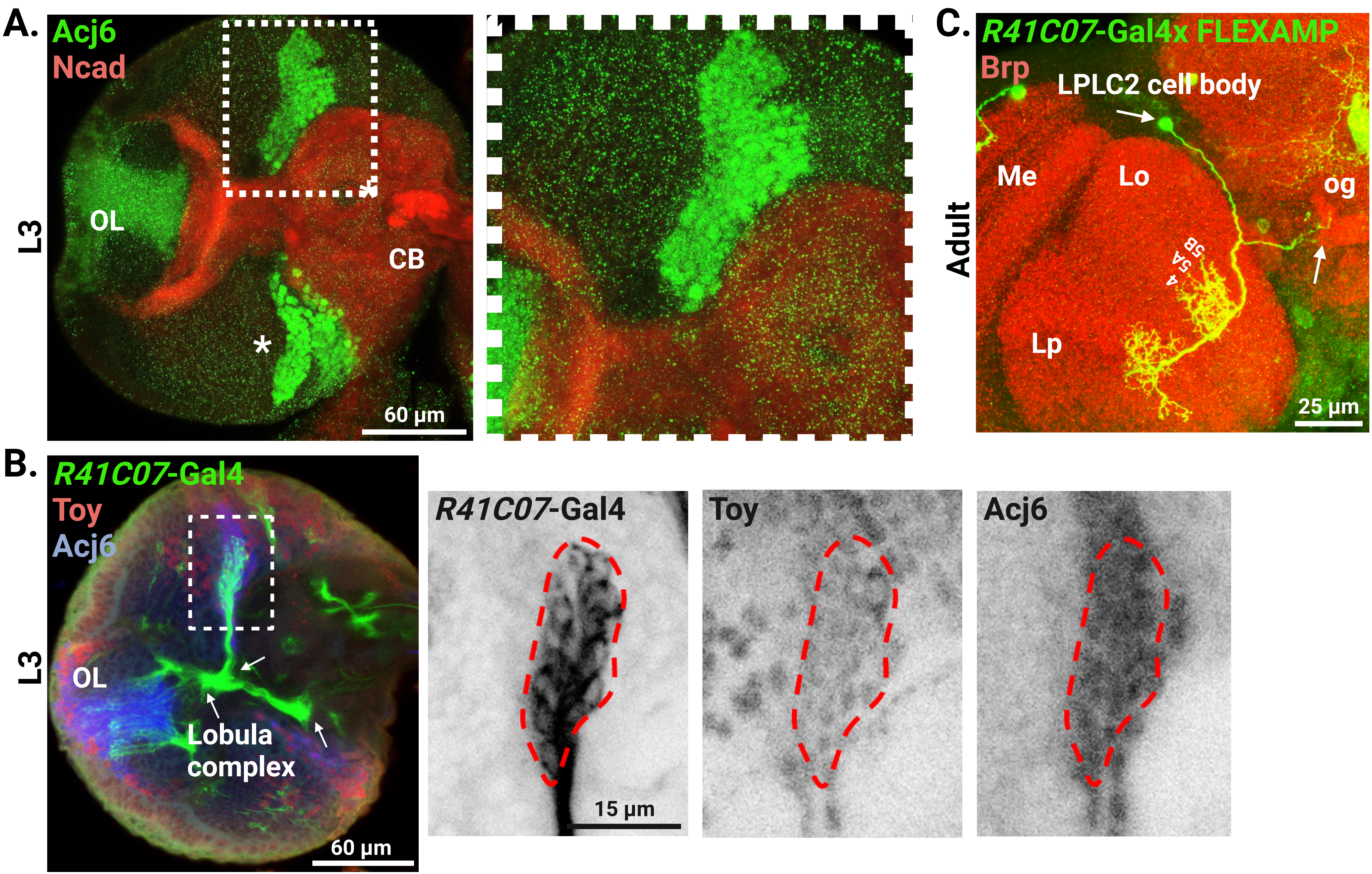

**Supp Figure 19: Dorsal central brain LCNs population.**

A. Expression pattern of Acj6 (green) with Ncad (red) in L3 brain. Inset showing a population of Acj6+ neurons in the dorsal central brain (CB) which correspond to LPLC1, LPLC2, LPLC4/LC22 (see Figure 7). Asterisk depicts vtIPC Acj6+ population (LC4, LC12 and LC17- see Figure 6). OL: optic lobe.

B. Expression pattern of *R41C07*-Gal4 (green) with Toy (red) and Acj6 (blue) in L3 larval brain. Note the LPLC2 population with projections to the Lobula complex and central brain (arrows). Inset shows a magnified view of LPLC2 cell bodies co-expressing Acj6 and Toy.

C. FLEXAMP in *R41C07*-Gal4 (green) with Brp (red) confirms the presence of LPLC2 neurons with projections in the lobula plate (Lp); lobula (Lo) layers 4, 5a and 5b; as well projection to the optic glomerulus (og) in the central brain. Me: medulla.
