## Supplementary Table 2 for "Morphological and functional convergence of visual projections neurons from diverse neurogenic origins in *Drosophila*"

| Reagent | Source | Identifier |
|---|---|---|
| **Fly strains** | | |
| Drosophila, OL0077B (LC6 split-Gal4) | BDSC (Wu et al., 2016) | 68247 |
| Drosophila, SS02651 (LC9 split-Gal4) | BDSC (Wu et al., 2016) | 68342 |
| Drosophila, OL0015B (LC11 split-Gal4) | BDSC (Wu et al., 2016) | 68362 |
| Drosophila, OL0027B (LC13 split-Gal4) | BDSC (Wu et al., 2016) | 68257 |
| Drosophila, OL0042B (LC15 split-Gal4) | BDSC (Wu et al., 2016) | 68258 |
| Drosophila, OL0046B (LC16 split-Gal4) | BDSC (Wu et al., 2016) | 68331 |
| Drosophila, OL0010B (LC18 split-Gal4) | BDSC (Wu et al., 2016) | 68358 |
| Drosophila, SS00343 (LC20 split-Gal4) | BDSC (Wu et al., 2016) | 68260 |
| Drosophila, OL0044B (LC21 split-Gal4) | BDSC (Wu et al., 2016) | 68351 |
| Drosophila, OL0001B (LC22 split-Gal4) | BDSC (Wu et al., 2016) | 68357 |
| Drosophila, SS02638 (LC24 split-Gal4) | BDSC (Wu et al., 2016) | 68340 |
| Drosophila, SS02650 (LC25 split-Gal4) | BDSC (Wu et al., 2016) | 68341 |
| Drosophila, SS02445 (LC26 split-Gal4) | BDSC (Wu et al., 2016) | 68333 |
| Drosophila, R93G05-Gal4 (LC4-Gal4) | BDSC | 40662 |
| Drosophila, R65B05-Gal4 (LC12/17-Gal4) | BDSC (Aptekar et al. 2015) | 49610 |
| Drosophila, R50C03-Gal4 (LPLC2-Gal4) | BDSC | 38734 |
| Drosophila, *wg*-Gal4 | ND382, K. Basler (Bertet et al., 2014) | |
| Drosophila, *UAS-Red.Stinger* | BDSC | 8547 |
| Drosophila, Ac76E MiMIC | BDSC | 59400 |
| Drosophila, Kn MiMIC | BDSC (Özel et al., 2021) | |
| Drosophila, sns MiMIC | BDSC | 59801 |
| Drosophila, Beat-Ic MiMIC | (Özel et al., 2021) | |

**Supplementary Table 2:**

| | | |
|---|---|---|
| Drosophila, slp-Gal4 | R35A08, Janelia Gal4 collection (Bertet et al., 2014) | |
| Drosophila, R80G09-Gal4 (ham-Gal4) | BDSC (Aptekar et al., 2015) | 40089 |
| Drosophila, RunxA MiMIC | BDSC | 76689 |
| Drosophila, R93G05-Lex A (LC4-LexA) | BDSC | 54310 |
| Drosophila, vtIPC-Gal4 | Desplan lab, Filipe Pinto-Teixeira | |
| Drosophila, R41C07-Gal4 | BDSC | 48145 |
| Drosophila, FLEXAMP y,w,UAS-FLP; If/CyO; act>y[+]>LHV2-86Fb,13XlexAop2-myr::GFP/TM6B | Bertet et al., 2014 | |
| **Antibodies** | | |
| mouse anti-acj6 | DSHB | anti-Acj6 |
| rat-anti bab2 | Laski lab; Couderc et al., 2002 | N/A |
| mouse-anti br | DSHB | 25E9.D7 |
| mouse-anti brp | DSHB | nc82 |
| rabbit-anti caps | Claude Desplan lab | N/A |
| mouse-anti con | DSHB | C1.427 |
| mouse-anti ct | DSHB | 2B10-S |
| guinea pig-anti D | Claude Desplan lab; Özel, Simon et al., 2020 | N/A |
| guinea pig-anti disco | Erclik lab; Valentino et al., 2022 | N/A |
| rat-anti E-cad | DSHB | DCAD2 |
| guinea pig-anti-Fred | Tanya Wolf lab; Fetting et al., 2001 | N/A |
| rabbit-anti GFP | Invitrogen | A11122 |
| sheep-anti GFP | Biorad | 4745-1051 |
| guinea pig-anti kn | Claude Desplan lab; Kostantinides et al., 2022 | N/A |
| guinea pig-anti lov | Kate Beckingham lab; Zhou et al., 2016 | N/A |
| rabbit-anti mirr | Helen McNeill lab | N/A |
| rat-anti Ncad | DSHB | DN-ex8 |
| rabbit-Olid2 | Claude Desplan lab; Erclik et al., 2017 | N/A |
| rat-anti pdm3 | Claude Desplan lab | N/A |
| mouse-anti pros | DSHB | MR1A |

**Supplementary Table 2:**

| | | |
|---|---|---|
| mouse-anti RFP | MBL | M155-3 |
| rabbit-anti RFP | Invitrogen | R10367 |
| rabbit-anti RunxA | Claude Desplan lab | N/A |
| guinea pig-anti Rx | Claude Desplan lab | N/A |
| guinea pig- anti salm | Claude Desplan lab | N/A |
| guinea pig-anti scro | Claude Desplan lab; Kostantinides et al., 2022 | N/A |
| rabbit-anti Sox102F | Claude Desplan lab; Kostantinides et al., 2022 | N/A |
| rabbit-anti SoxN | Claude Desplan lab | N/A |
| mouse-anti Ten-m | DHSB | MAb20 |
| rabbit-anti toy | Claude Desplan lab;<br>Özel, Simon et al., 2020 | N/A |
| rat anti-toy | Claude Desplan lab; Kostantinides et al., 2022 | N/A |
| Donkey-anti guinea pig Alexa Fluor 555 | Sigma-Aldrich | SAB4600297-5OUL |
| Donkey-anti guinea pig Alexa Fluor 647 | Jackson ImmunoResearch | 706-605-148 |
| Donkey-anti mouse Alexa Fluor 555 | ThermoFisher Scientific | A31570 |
| Donkey-anti mouse Alexa Fluor 647 | Jackson ImmunoResearch | 715-605-151 |
| Donkey-anti rabbit Alexa Fluor 488 | ThermoFisher Scientific | A31570 |
| Donkey-anti rabbit Alexa Fluor 555 | Invitrogen | A31572 |
| Donkey-anti rat Alexa Fluor 405 | Jackson ImmunoResearch | 712-475-153 |
| Donkey-anti rat Alexa Fluor 555 | Abcam | Ab150154 |
| Donkey-anti rat Alexa Fluor 647 | Jackson ImmunoResearch | 712-605-153 |
| Donkey-anti rat cy5 | Jackson ImmunoResearch | 712-175-153 |
| Donkey-anti sheep Alexa Fluor 488 | Jackson ImmunoResearch | 713-545-147 |
